# Dual-color Fluorescence Cross-Correlation Spectroscopy to study Protein-Protein Interaction and Protein Dynamics in Live Cells

**DOI:** 10.1101/2021.09.10.459760

**Authors:** Katherina Hemmen, Susobhan Choudhury, Mike Friedrich, Johannes Balkenhol, Felix Knote, Martin Lohse, Katrin G. Heinze

## Abstract

We present a protocol and workflow to perform live cell dual-color fluorescence crosscorrelation spectroscopy (FCCS) combined with Förster Resonance Energy transfer (FRET) to study membrane receptor dynamics in live cells using modern fluorescence labeling techniques. In dual-color FCCS, where the fluctuations in fluorescence intensity represents the dynamical “fingerprint” of the respective fluorescent biomolecule, we can probe co-diffusion or binding of the receptors. FRET, with its high sensitivity to molecular distances, serves as a well-known “nanoruler” to monitor intramolecular changes. Taken together, conformational changes and key parameters such as local receptor concentrations, and mobility constants become accessible in cellular settings.

Quantitative fluorescence approaches are challenging in cells due to high noise levels and the vulnerable sample itself. We will show how to perform the experiments including the calibration steps. We use dual-color labeled β2-adrenergic receptor (β_2_AR) labeled (eGFP and SNAPtag-TAMRA). We will guide you step-by-step through the data analysis procedure using open-source software and provide templates that are easy to customize.

Our guideline enables researchers to unravel molecular interactions of biomolecules in live cells *in situ* with high reliability despite the limited signal-to-noise levels in live cell experiments. The operational window of FRET and particularly FCCS at low concentrations allows quantitative analysis near-physiological conditions.

Link to accompanying video: https://tr240.uni-wuerzburg.de/vippclass/index.php/s/TL8aWmwE9RjGfLE

## Introduction

Fluorescence spectroscopy is one of the main methods to quantify protein dynamics and protein-protein interactions with minimal perturbation in a cellular context. Confocal fluorescence correlation spectroscopy (FCS) is one powerful method to analyze molecular dynamics as it is single molecule sensitive, highly selective, and live-cell compatible ^1^. Compared with other dynamics orientated approaches, FCS has a broader measurable time range spanning from ∼ ns to ∼ s, most importantly covering the fast time scales that are often inaccessible by imaging-based methods. Moreover, it also provides spatial selectivity so that membrane, cytoplasmic and nucleus molecular dynamics can be easily distinguished ^2^. Thus, molecular blinking, the average local concentration and the diffusion coefficient can be quantitatively analyzed with FCS; intermolecular dynamics such as binding become easily accessible when probing co-diffusion of two molecular species in fluorescence cross-correlation spectroscopy (FCCS) analysis ^3-5^ in a dual color approach. The main underlying principle in correlation spectroscopy is the statistical analysis of intensity fluctuations emitted by fluorescently labeled biomolecules diffusing in and out of a laser focus (Figure 1A). The resulting auto- or cross-correlation functions then can be further analyzed by curve fitting to eventually derive the rate constants of interest. In other words: The statistical methods FCS and FCCS do not provide single molecule traces like in single particle tracking, but a dynamic pattern or “fingerprint” of a probed specimen with high temporal resolution. Combined with Förster resonance energy transfer (FRET), also *intra*molecular dynamics such conformational changes can be monitored at the same time in a common confocal setup ^5,6^. FRET probes the distance of two fluorophores and is often referred to as a molecular “nano ruler”. Energy transfer takes place only when i) the molecules in close vicinity (3-10 nm), ii) the emission spectrum of the donor significantly overlaps with absorption spectrum of the acceptor molecule, and iii) the dipole orientation of donor and acceptor is (almost) parallel. Thus, the combination of FRET and FCCS provides a technique with very high spatio-temporal resolution.

**Figure 1.**
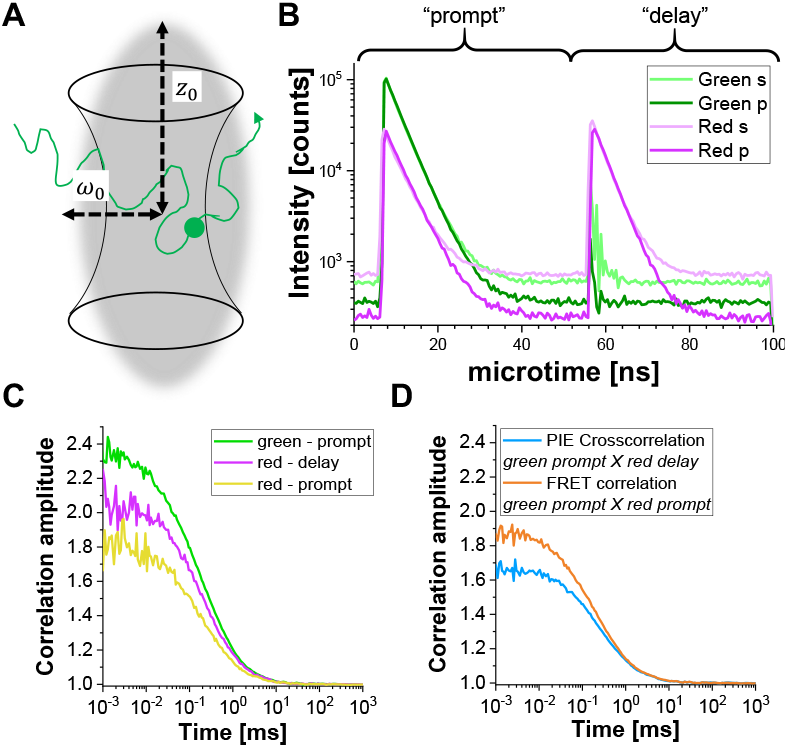
Pulsed-interleaved excitation (PIE) based fluorescence (cross) correlation spectroscopy (F(C)CS). (A) In FCS fluorescently labeled molecules diffuse freely in and out of a confocal detection volume, in which they are excited by a laser pulse and their emitted fluorescence is collected. The resulting intensity fluctuations are correlated and provide information on e.g. the mobility of the molecules. (B) In PIE, two different laser lines (“prompt” and “delay”) are used to excite the sample labeled with two different fluorophores (“green” and “red”). The time difference between both excitation pulses is chosen based on the fluorescence lifetime of the selected fluorophores: Their respective fluorescence should have decayed. In the shown double-labeled sample, both fluorophores are close enough together to undergo Förster Resonance Energy Transfer (FRET), and the “green” or donor fluorophore transfer part of its energy to the “red” or acceptor fluorophore. Thus, red fluorescence emission occurring in the “prompt” time-window, i.e. after green excitation. In the used setup (Supplementary Note 2), two detectors are used for each color, one oriented parallel to the excitation beam orientation (denoted “p”) and the second perpendicular (denoted “s”). (C) Three different autocorrelation functions can be determined in a PIE experiment: Correlation of the signal collected in the green channels in the prompt time window (*ACF*_*gp*_), signal collected in the red channels in the prompt time window (*ACF*_*rp*_) and the signal collected in the red channels in the delay time window (*ACF*_*rd*_). (D) Two different crosscorrelation functions can be constructed: In the “PIE” crosscorrelation, *CCF*_*PIE*_, the signal collected in the green channels in the prompt time window is correlated with the signal collected in the red channels in the delay window. The amplitude of this curve is related to the co-diffusion of fluorophores. In the “FRET” crosscorrelation, *CCF*_*FRET*_, the signal collected in the green channels in the prompt time window is correlated with the signal collected in the red channels in the same prompt window. The shape of this curve at times faster than diffusion is related to the FRET-induced intensity changes.

When spatial selectivity, sensitivity as well as live-cell compatibility is required, FRET-FCCS has obvious advantageous over other methods such as ITC ^7^, SPR ^8^ or NMR ^9,10^ when it comes to measuring protein dynamics and interactions.

Despite the capabilities and promise of dual color fluorescence cross-correlation spectroscopy (dc-FCCS), performing dc-FCCS in live cells is technically challenging due to spectral bleed-through or cross-talk between the channels ^3,4^, difference in the confocal volumes due to the spectrally distinct laser lines ^3,4,11^, background signal and noise or limited photostability of the samples ^12-15^. Introduction of pulse interleaved excitation (PIE) to FCCS was an important “tweak” to temporally decouple the different laser excitations to reduce the spectral crosstalk between the channels ^16^. Other correction methods to counter spectral bleed-through ^17-19^and background have also been well-accepted ^17-19^. For details and basics on FCS, PIE or FRET the reader is referred to the following references ^2,4,6,16,20-24^.

Here, first all necessary calibration experiments were performed and analyzed before experimental results of a prototypical G-protein coupled receptor, β_2_-adrenergic receptor (β_2_AR), for three different scenarios are shown: (1) Single-labeled molecules carrying either a “green” (eGFP) or a “red” (SNAP tag-based labeling) ^25^ fluorophore. (2) A double-labeled construct, which carries an N-terminal SNAP tag and intracellular eGFP (NT-SNAP). In this case, both labels are at the same protein, thus 100 % co-diffusion was expected. (3) A double-labeled sample, where both fluorophores are on the same side of the cell membrane (CT-SNAP). It carries a C-terminal SNAP tag and an intracellular eGFP. Here, again both labels are at the same protein with again 100 % co-diffusion expected. As both labels are very close to each other, on the same side of the cell membrane, it shows the potential to observe FRET and anticorrelated behavior. All constructs were transfected in Chinese Hamster Ovary (CHO) cells and later labeled with a red fluorescent substrate which is membrane-impermeable for the NT-SNAP construct and a membrane-permeable substrate for the CT-SNAP construct. Finally, simulated data exemplifies the influence of experimental parameter on the FRET-induced anticorrelation, and the effect of protein-protein interactions on the co-diffusion amplitude.

Thus, this protocol provides a complete guide to perform the combined approach of FRET-FCCS in living cells to understand protein dynamics and protein-protein interactions while making aware of technical / physical artifacts, challenges and possible solutions.

## Protocol

### 1. Experimental protocol

#### 1.1. Sample preparation

##### 1.1.1. Cell seeding

Important note: Cell seeding and transfection need to be performed under sterile conditions.

1. Place a cleaned coverslip per well onto a 6-well culture plate and wash three times with sterile phosphate-buffered saline (PBS). Note: The coverslip cleaning protocol is detailed in **Supplementary Note 1**.
2. Add 2 ml of cell culture medium with phenol red supplemented with 10 % fetal bovine serum (FBS), 100 μg/ml penicillin and 100 μg/mL streptomycin to each well and keep it aside.
3. Take the CHO cells, which are cultured in the same medium with phenol red at 37 °C, in 5 % CO_2_ and wash them with 5 ml PBS to remove the dead cells.
4. Add 2 mL of trypsin and incubate for 2 min at room temperature (RT).
5. Dilute the detached cells with 8 mL of medium with phenol red and mix carefully by pipetting.
6. Count the cells in a Neubauer chamber and seed the CHO cells at a density of 1.5 × 10^5^ cells/well in the 6-well cell culture plate containing the coverslips.
7. Let the cells grow in an incubator (37 °C, 5 % CO_2_) for 24 hr in order to achieve approx. 80 % confluency.

##### 1.1.2. Transfection

1. Dilute 2 µg of the desired vector DNA (e.g. CT-SNAP or NT-SNAP) and 6 µL of the transfection reagent in two separate tubes each containing 500 µL reduced-serum medium for each well and incubate them for 5 min at RT.
2. Mix the two solutions together to obtain the transfection mixture and incubate it for further 20 min at RT.
3. In the meantime, wash the seeded CHO cells once with sterile PBS.
4. Replace the PBS with 1 mL/well of phenol red-free medium supplemented with 10 % FBS and no antibiotics.
5. Add the entire transfection mixture of 1 mL dropwise to each well and incubate the cells overnight at 37 °C, in 5% CO_2._

##### 1.1.3. Labeling of SNAP constructs

1. Dilute the appropriate SNAP substrate stock solution in 1 mL medium supplemented with 10 % FBS to obtain a final concentration of 1 µM.
2. Wash the transfected cells once with PBS and add 1 mL per well of 1 µM SNAP substrate solution.
3. Incubate the cells for 20 min at 37 °C in 5 % CO_2_.
4. Wash the cells thrice with phenol red-free medium and add 2 mL per well phenol red-free medium.
5. Incubate the cells for 30 min at 37 °C in 5 % CO_2_.

##### 1.1.4. Transfer to measurement chamber

1. Transfer the coverslips of all samples subsequently into the imaging chamber and wash with 500 µL imaging buffer.
2. Add 500 µL imaging buffer before moving to the FRET-FCS setup.

#### 1.2. Calibration Measurements

The FRET-FCS setup is equipped with a confocal microscope water objective, two laser lines, a Time-Correlated Single Photon Counting (TCSPC) system, two hybrid photomultiplier tubes (PMT) and two avalanche photodiodes (APD) for photon collection and the data collection software. It is very crucial to align the setup every time before measurements in live cells.

Note: The detailed setup description can be found in **Supplementary Note 2**.

Important note: Both lasers and all detectors (two PMTs and two APDs) are always on during the measurements, as all measurements need to be conducted under identical conditions. Use a coverslip from the same lot on which the cells were seeded, this decreases the variation in collar ring correction.

##### 1.2.1. Adjusting focus, pinhole and collar ring position

1. Place 2 nM green calibration solution on a glass coverslip and switch on the 485 nm and 560 nm laser operated in Pulsed Interleaved Excitation (PIE) mode ^16^.
2. Focus on the solution and adjust the pinhole and collar ring position such that the highest count rate and smallest confocal volume are obtained to get the maximum molecular brightness.
3. Repeat this process for the red channels with 10 nM red calibration solution and a mixture of both.

##### 1.2.2. Optimizing the confocal overlap volume

1. Place the 10 nM DNA solution on glass coverslip and adjust the focus, pinhole, and collar ring position such that the cross correlation between the green and red detection channels is highest, i.e. shows the highest amplitude.

Notes:

Steps 1.2.1 and 1.2.2 might have to be repeated back and forth to find the optimal alignment. Take 3-5 measurements from each calibration solution for 30 s – 120 s after the focus, pinhole, and collar ring position have been aligned optimally for the green and red detection channels and the confocal overlap volume.

Measure a drop of ddH2O, the imaging medium and a non-transfected cell for 3-5 times each for 30 s – 120 s to determine the background count rates.

##### 1.2.3. Optional step: Instrument response function

Optional but highly recommended: Collect the instrument response function with 3-5 measurements for 30 s – 120 s.

#### 1.3. Measurements in live cells

1. Find a suitable cell by illuminating with the Mercury lamp and observing through the ocular. Note: Suitable cells are alive and the fluorescence of the protein of interest, here a surface receptor, is visible all over the surface. Less bright cells are more suitable than brighter ones due to the better contrast in FCS when a low number of molecules is in the focus.
2. Switch on both lasers in PIE mode and focus on the membrane by looking for the maximum counts per second. Note: The laser power might need to be reduced for the cell samples (less than 5 µW at objective). This depends highly upon the used fluorophores and the setup.
3. Observe the auto- and cross-correlation curves of the β_2_AR bound to eGFP and / or SNAP-tagged probes in the online preview of the data collection software and collect several short measurements (2 −10) with an acquisition time between 60 −180 s Important note: Do not excite the cells for long time continuously as the fluorophores may bleach. However, it will depend on the brightness of each cell, how long the measurements can be and how many measurements in total can be performed.

### 2. Data analysis

#### 2.1. Data Export

Export the correlation curves G(*t*_*c*_) and count rates CR from all measurements. Please take care here to correctly define the “prompt” and “delay” time windows and use the “microtime gating” option in the data correlation software. In total, three different correlations are required: (i) autocorrelation of the green channel in prompt time window (*ACF*_*gp*_), (ii), autocorrelation of the red channels in the delay time window (*ACF*_*rd*_), and finally (iii) the cross-correlation of the green channel signal in the prompt time window with the red channel signal in the delay time window (*CCF*_*PIE*_).

Note: The data export is shown step-by-step for different software in **Supplementary Note 3**.

#### 2.2. Calibration measurements

##### 2.2.1. Determination of the confocal detection volume

1. Use the autocorrelation functions of the green (*ACF*_*gp*_) and red (*ACF*_*rd*_) fluorophore solutions, and fit them to a 3D diffusion model with an additional triplet term if required (eq. 1) to calibrate the shape and size of the confocal detection volume for the two used color channels:

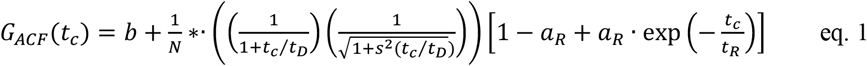

where *b* is the baseline of the curve, *N* the number of molecules in focus, *t*_*D*_ the diffusion time (in ms) and *s* = *z*_*0*_/*ω*_*0*_ the shape factor of the confocal volume element. The triplet blinking and photophysics is described by its amplitude *a*_*R*_ and relaxation time *t*_*R*_ Note: All variables and symbols used within the protocol are listed in Table 1.
2. Use the known diffusion coefficients *D* for the green ^26^ and red calibration standard ^27^ and the obtained shape factors *s*_*green*_ and *s*_*red*_ to determine the dimensions (width ω_0_ and height z_0_) and volume *V*_*eff*_ of the confocal volume element is determined (eq. 2a-c).

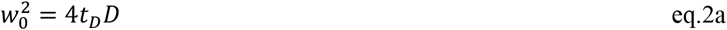

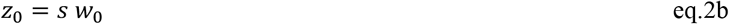

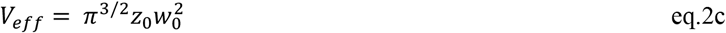

Note: Templates for calculation of the calibration parameter are provided as Supplementary files (S7).

**Table 1.**
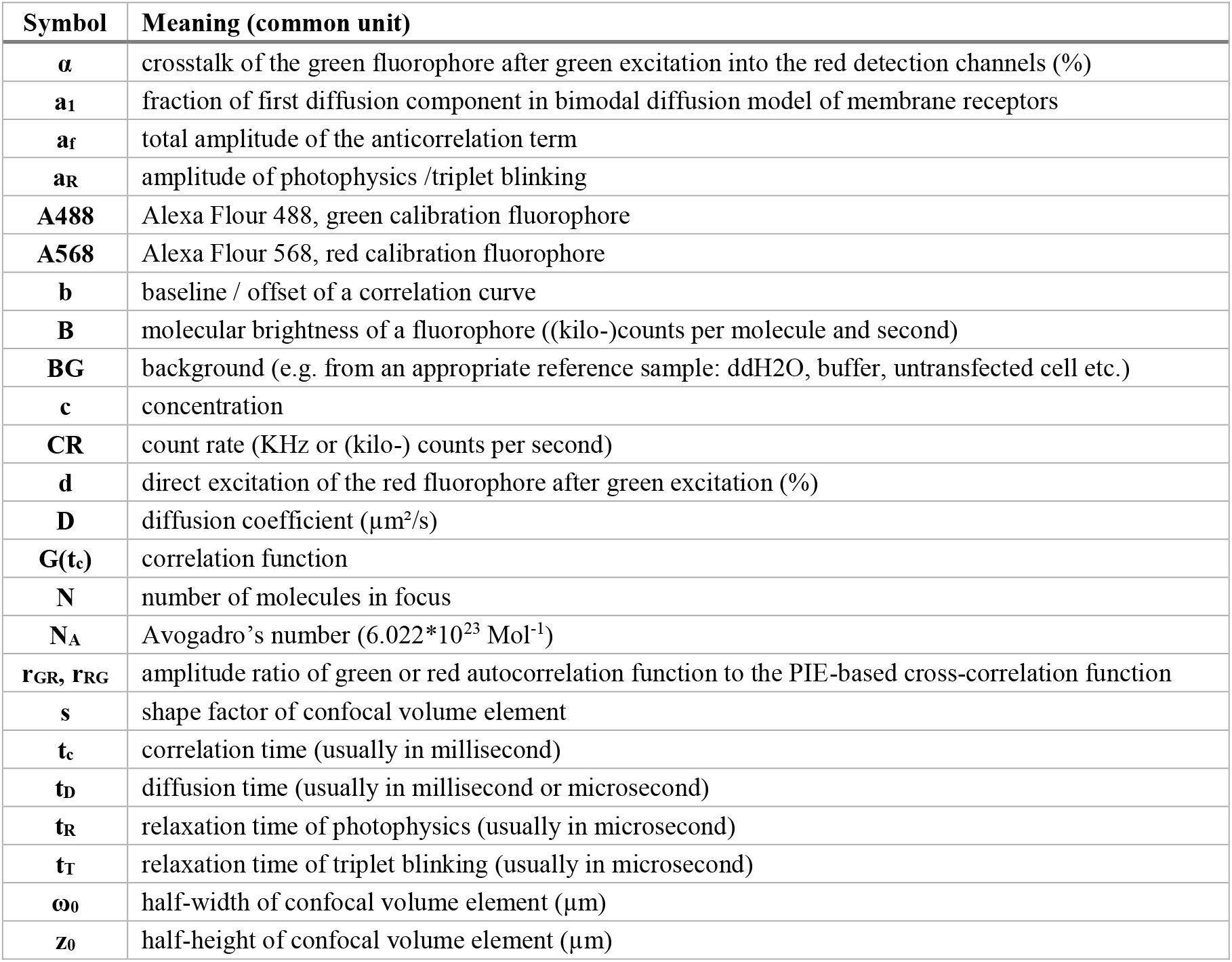
List of variables and abbreviations. For the use of symbols and definition in fluorescence and FRET experiments, the guidelines of the FRET community ^51^ are recommended.

##### 2.2.2. Spectral crosstalk α of green into red channel

Calculate the spectral crosstalk *α* of the green fluorescence signal (collected in channels 0 and 2) into the red detection channels (channel number 1 and 3) is as a ratio of the background-corrected (BG) signals (eq. 3).

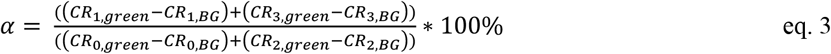

##### 2.2.3. Direct acceptor excitation δ

Determine the direct excitation of the acceptor fluorophore *δ* by the donor excitation wavelength by the ratio of the background-corrected count rate of the red calibration measurements in the “prompt” time window (excitation by green laser) to the background-corrected count rate in the “delay” time window (excitation by red laser) are used (eq. 4).

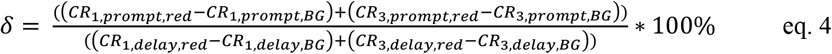

##### 2.2.4. Molecular brightness *B*

Calculate the molecular brightness *B* of both the green and red fluorophores (eq. 5a-b) based on the background-corrected count rates and the obtained number of molecules in focus, *N*, from the 3D diffusion fit (eq. 1):

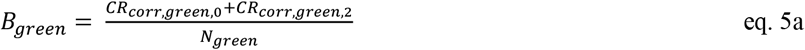

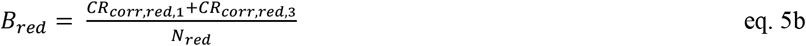

##### 2.2.5. Overlap of green and red confocal detection volume

1. Fit both *ACF*_*gp*_ and *ACF*_*rd*_ as well as *CCF*_*gp-rd*_ of the double-labeled DNA to the 3D diffusion model (eq. 1). Keep the obtained shape factors, *s*_*green*_ and *s*_*red*_, constant for *ACF*_*gp*_ and *ACF*_*rd*_, respectively. The shape factor for the *CCF*_*gp-rd*_, *s*_*PIE*,_ is usually in between these two values. Note: In an ideal setup, both *V*_*eff, green*_ and *V*_*eff, red*_ would have the same size and overlap perfectly.
2. Determine the amplitude at zero correlation time, *G*_*0*_(*t*_*c*_), based on the found values of the apparent number of molecules in focus (*N*_*green*_, *N*_*red*_ and *N*_*PIE*_).
3. Calculate the amplitude ratio *r*_*GR*_ and *r*_*RG*_ for a sample with 100 % co-diffusion of green and red fluorophores (eq. 6): Note: Be aware that *N*_*PIE*_ is not reflecting the number of double-labeled molecules in the focus but reflects only the 1/*G*_*0*_(*t*_*c*_).

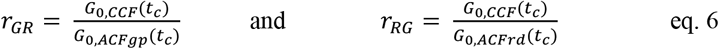

#### 2.3. Live cell experiments

##### 2.3.1. Single-labeled constructs

1. Fit the cell samples to an appropriate model. For the shown membrane receptor, diffusion occurs in a bimodal fashion with a short and a long diffusion time. Additionally, the photophysics and blinking of the fluorophores have to be considered:

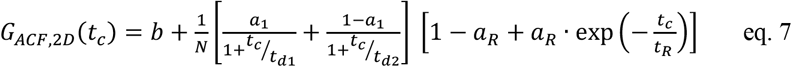

where *t*_*d1*_ and *t*_*d2*_ are the two required diffusion times and *a*_*1*_ is the fraction of the first diffusion time.. Important note: In contrast to the calibration measurements, in which the free dyes and DNA strands freely diffuse in all directions, membrane receptor shows only 2D diffusion along the cell membranes. This difference between 3D and 2D diffusion is reflected by the modified diffusion term (compare eq. 1), where *t*_*D*_ in the 2D case does not depend upon the shape factor *s* of the confocal volume element.
2. Calculate the concentration *c* of green or red labeled proteins from the respective *N* and *V*_*eff*_ using basic math (eq. 8):

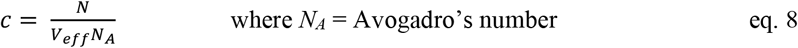

##### 2.3.2. N-terminal SNAP label and intracellular eGFP

1. Fit the two autocorrelations (*ACF*_*gp*_ and *ACF*_*rd*_) of the double-labeled sample using the same model as for the single-labeled constructs for the ACFs (eq. 7) and the *CCF*_*PIE*_ using a bimodal diffusion model (eq. 9):

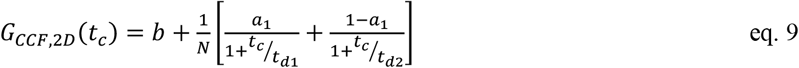

Note: For a global description of the system, all three curves have to be fit jointly: The diffusion term is identical for all three curves and the only difference is the relaxation term for the *CCF*_*PIE*_. As photophysics of two fluorophores is usually unrelated no correlation term is required. This absence of relaxation terms results in a flat *CCF*_*PIE*_ at short correlation times. However, crosstalk and direct excitation of the acceptor due to the donor fluorophore might show false-positive amplitudes and should be carefully checked for using the calibration measurements.
2. Calculate the concentration *c* of green or red labeled proteins from the respective *N* and *V*_*eff*_ using basic math (eq. 8).
3. Estimate the fraction or concentration, *c*_*GR*_ or *c*_*RG*_, of interacting green and red labeled proteins from the cell samples using the correction factors obtained from the DNA samples, the amplitude ratios *r*_*GR*_ and *r*_*RG*_ of the cell sample and their respective obtained concentrations (eq. 10).

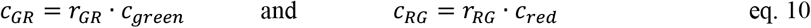

##### 2.3.3. C-terminal SNAP label and intracellular eGFP

Fit the two autocorrelations (*ACF*_*gp*_ and *ACF*_*rd*_) of the FRET sample as the single-labeled samples (eq. 7) and the *CCF*_*FRET*_ to a bimodal diffusion model containing an anticorrelation term (eq. 11):

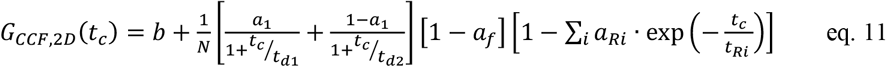

where a_*f*_ reflects the amplitude of the total anti-correlation and *a*_*R*_ and *t*_*R*_ the respective amplitude and relaxation time. Note: In case of anti-correlated fluorescence changes due to FRET one or several anti-correlation terms might be required (eq. 11) resulting in a “dip” of *CCF*_*FRET*_ at low correlation times coinciding with a rise in the two autocorrelations (*ACF*_*gp*_ and *ACF*_*rd*_). However, be aware that photophysics such as triplet blinking might mask the anti-correlation term by dampening the FRET-induced anti-correlation. A joint analysis supplemented with filtered FCS methods might help to unmask the anti-correlation term. Additionally, technical artifacts stemming from dead times in the counting electronics in the nanoseconds range should be excluded ^16^.

A more detailed step-by-step procedure on how to perform the analysis in ChiSurf ^28^ and excel templates e.g. for the calculation of confocal volume or molecular brightness are provided on our github repository (https://github.com/HeinzeLab/JOVE-FCS) and as supplementary files (Supplementary Note 4 and S6). Additionally, our python-scripts for batch export of data acquired with the Symphotime software in .ptu format can be found there.

## Result

Below, exemplary the results of calibration and live-cell measurements are shown. Additionally, the effect of FRET on the cross-correlation curves is shown based on simulated data next to exemplifying the effect of protein-protein-interaction increasing the *CCF*_*PIE*_ amplitude.

### 1. PIE-based FCS data export

In PIE experiments, data are collected in the time-tag time-resolved mode (TTTR) ^29,30^. Figure 1B shows exemplary the photon arrival time histograms of a PIE measurement of a double-labeled DNA strand on the described setup (Supplementary Note 1). The setup contains in total four detection channels, two for the green fluorescence signal and two for the red fluorescence signal. Additionally, the photons are sorted by their polarization – either parallel or perpendicular to the excitation direction. In the “prompt” time window, the green fluorophore gets excited and signal is detected in both the green and – due to FRET - red channels, while in the delay time window the red fluorophore gets excited and signal is only detected in the red channels. Based on the detection channels and time windows at least five different correlation curves can be reconstructed (Figure 1C-D). The autocorrelations (ACF) of the green signal in the prompt time window (*ACF*_*gp*_), of the red signal in the prompt time window (in case of FRET, *ACF*_*rp*_) and lastly of the red signal in the delay time window (*ACF*_*rd*_) report on the protein mobility, photophysics (e.g. triplet blinking) and other time-correlated brightness changes in the fluorophores (e.g. due to FRET). The PIE-based crosscorrelation *CCF*_*PIE*_ of the green signal in the prompt time window with the red signal in the delay time window allows determining the fraction of co-diffusion of the green and red fluorophore ^16^. The FRET-based crosscorrelation *CCF*_*FRET*_ of the green with the red signal in the prompt time window is related to FRET-induced, anticorrelated brightness changes in the green and red signals ^31-33^.

### 2. Calibration

Figure 2A-B shows a calibration measurement of the singly diffusing green and red fluorophores, respectively. Based on a fit with eq. 1 and the known diffusion coefficient D_green_ ^26^ and D_red 27_ the shape (*z*_*0*_ and *ω*_*0*_) and size (*V*_*eff*_) of the detection volume is calculated using eq. 2a-c. The fit results from the *ACF*_*gp*_ from the green fluorophore and *ACF*_*rd*_ from the red fluorophore are summarized in Figure 2C. Both fluorophores show an additional relaxation 8.6 µs (18 %) and 36 µs (15 %) for, respectively. The molecular brightness (eq. 5a-b) of the green and red fluorophore amounts to 12.5 kHz per molecule and 2.7 kHz per molecule, respectively, for the used excitation conditions.

For a reliable estimation of the confocal volume size and shape as well as the molecular brightness, it is recommended to perform 3-5 measurements per calibration experiments and to perform a joint (or global) fit of all repeats.

The crosstalk α (Figure 2D, eq. 3) and the direct excitation of acceptor by the green laser δ (Figure 2E, eq. 4) for this fluorophore pair lies at ∼15 % and ∼ 38 %.

To determine and calibrate the overlap of the green and red excitation volume a double-labeled double DNA strand is used (Figure 3A) as described above. Here, the fluorophores are spaced 40 bp apart such that no FRET can occur between the green and red fluorophores attached to the ends of the DNA double strands. Figure 3B shows the autocorrelations from both fluorophores in green (*ACF*_*gp*_) and magenta (*ACF*_*rd*_) and the PIE-crosscorrelation, *CCF*_*PIE*_, in cyan. As a reminder, for *CCF*_*PIE*_ the signal in the green channels in the prompt time window is correlated with the signal in the red channels in the delay time window ^16^.

Here, we obtain an average diffusion coefficient for the DNA strand of *D*_*DNA*_ = 77 µm^2^/s is obtained (More details on the calculation can be found in the step-by-step protocol, Supplementary Note 4). This value is found when inserting the calibrated green and red detection volumes size (Figure 2) and the respective diffusion times of *ACF*_*gp*_ and *ACF*_*rd*_ of the DNA strand (Figure 3C) into eq. 2a. Next, using the obtained correction values *r*_*GR*_ and *r*_*RG*_, determined from eq. 6 later on the amount of co-diffusing, i.e. double labeled molecules (or protein complexes in case of co-transfection of two different proteins) can be determined from the cell samples.

**Figure 2.**
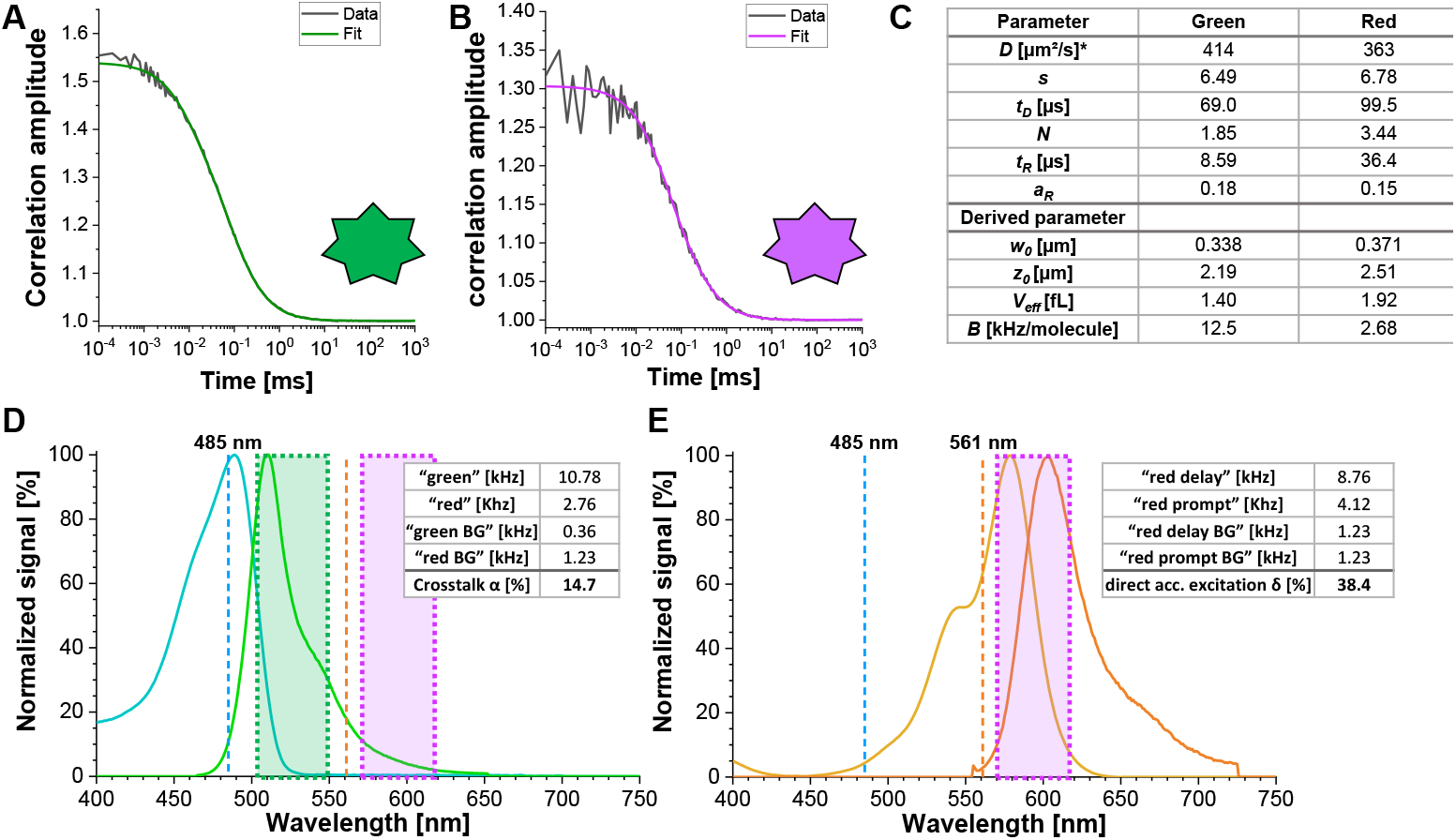
Calibration measurements of freely diffusing green and red calibration standard. (A-B) Representative 60 s measurement of a 2 nM green (A) and a 10 nM red (B) calibration standard measurement fitted to the 3D Diffusion model including an additional relaxation time (**eq. 1**). The table in panel (C) shows the fit results and the derived parameter based on **eq. 2a-c** and **eq. 5a-b**. ^*^Diffusion coefficients were taken from literature ^26,27^. (D) Determination of the crosstalk α of the green signal into the red channels (**eq. 3**). The excitation spectrum of green standard is shown in cyan, in green the emission spectrum. The excitation laser lines at 485 nm (blue) and 561 nm (orange) are shown as dashed lines. Transparent green and magenta boxes show the collected emission range (**Supplementary Note 2**). (E) Determination of the direct excitation δ of the red fluorophore by the 485 nm laser (eq. 4). Color code is identical to (D), light and dark orange show the excitation and emission spectrum of the red standard, respectively.

**Figure 3.**
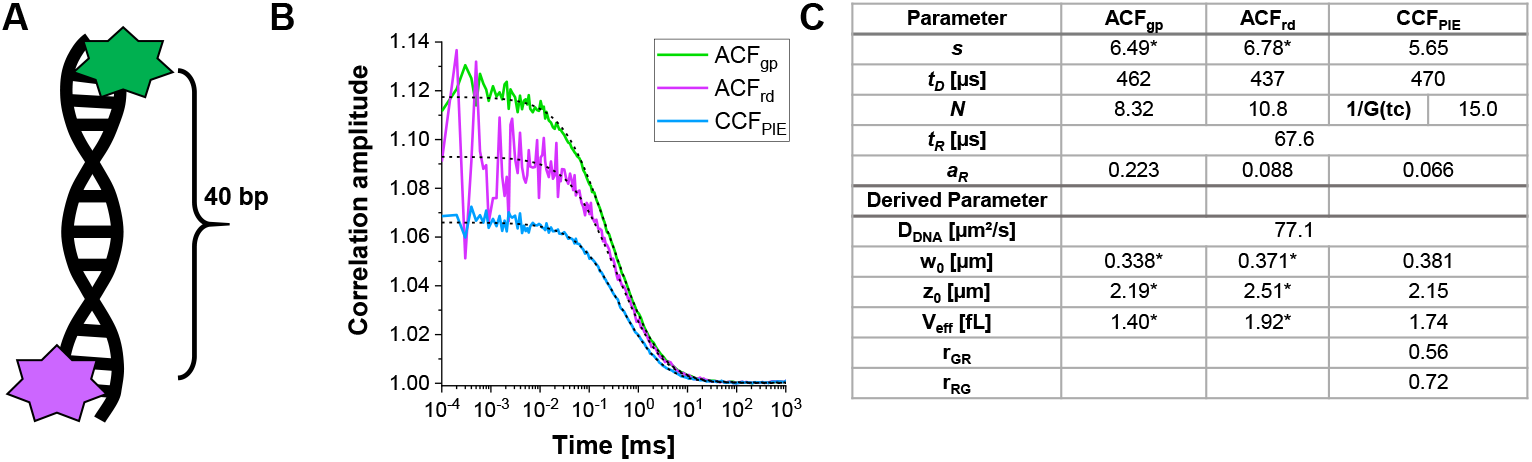
Calibration of the green-red overlap volume using a DNA sample. (A) The DNA strand used for calibration carries a the green and an red calibration fluorophore, with a distance of 40 bp inbetween. The interdye distance must be sufficiently large to exclude FRET between the fluorophores. (B) Representative 60 s measurement of a 10 nM DNA solution. Autocorrelations from both fluorophores in green (*ACF*_*gp*_, green standard) and magenta (*ACF*_*rd*_, red standard) and the PIE-crosscorrelation, *CCF*_*PIE*_, in blue. The table in panel (C) shows the fit results based on the 3D Diffusion model including an additional relaxation term (**eq. 1**) and the derived parameter diffusion coefficient of DNA, *D*_*DNA*_, (**eq. 2a**), the size and shape of the overlap volume (**eq. 2a-c**) and the correction ratios r_GR_ and r_RG_ (**eq. 6**). Please note the values for the green and red detection volume (labeled with ^*^) were taken from the fit of the individual fluorophores shown in Figure 2.

### 3. Live-cell experiments

In the following section, the analysis of live-cell experiments for different β_2_AR constructs are shown. As β_2_AR is a membrane protein, its diffusion is largely limited to a two-dimensional diffusion (Figure 4A) along the cell membrane (except of transport or recycling processes to or from the membrane) ^2^. This restriction to the 2D diffusion is reflected in the modified diffusion term of eq. 9 compared to eq. 1: it is independent from the shape factor.

**Figure 4.**
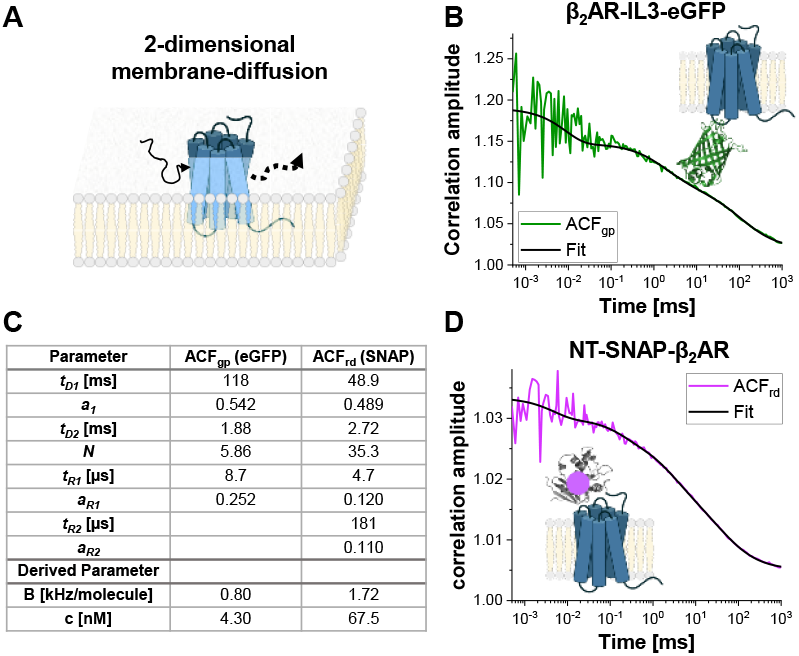
Representative measurement of single-label constructs. (A-B) In this study, the membrane receptor β_2_AR was used as a example. In contrast to the fluorophores and DNA strand used for calibration, which could freely float through the detection volume, membrane proteins diffuse mainly laterally along the membrane, described as 2-dimensional diffusion. (B,D) *ACF*_*gp*_ and *ACF*_*rd*_ of the single-label constructs β_2_AR-IL3-eGFP (B) and NT-SNAP-β_2_AR (D). Shown is the average of 4-6 measurements with each 120 – 200 s long. The table in panel (C) shows the fit results of the data to the bimodal two-dimensional diffusion model including additional relaxation terms (**eq. 7**).

#### 3.1. Single-labeled constructs: β_2_AR-IL3-eGFP and NT-SNAP-β_2_AR

Figure 4 shows exemplary measurements of the single-label constructs β_2_AR-IL3-eGFP (Figure 4B), where the eGFP is inserted into the intracellular loop 3, and NT-SNAP-β_2_AR (Figure 4C), where the SNAP tag is conjugated to the N-terminus of β_2_AR. The SNAP tag is labeled with a membrane-impermeable SNAP surface substrate. The shown curves represent the average of 4-6 repeated measurements each 120 - 200 s long. The respective autocorrelations *ACF*_*gp*_ and *ACF*_*rd*_ of the eGFP and SNAP signal are fitted to a bimodal, two-dimensional diffusion model shown in eq. 9. The *ACF*_*gp*_ from eGFP shows only the expected triplet blinking at *t*_*R1*_ ∼ 9 µs. Interestingly, the *ACF*_*rd*_ of the SNAP signal requires two additional relaxation times, one at the typical triplet blinking time of *t*_*R1*_ ∼ 5 µs and a second one at *t*_*R2*_ ∼ 180 µs.

The molecular brightness of eGFP and the SNAP substrate in live cells under the used excitation conditions amounts to 0.8 and 1.7 kHz per molecule, respectively (eq. 5a-b). The concentration of the labeled β_2_AR constructs incorporated into the cell membrane lies in the nano-molar range and can be determined based on the obtained number of molecules (eq. 9, Figure4C) and the size of the respective confocal volume for the green and red channel (Figure 2) using eq. 8.

#### 3.2. Double-labeled construct: NT-SNAP-β_2_AR-IL3-eGFP

In the double-labeled construct NT-SNAP-β_2_AR-IL3-eGFP (short NT-SNAP), eGFP is inserted into the intracellular loop 3 and additionally the SNAP tag is conjugated to the N-terminus of β_2_AR (Figure 5A). As both fluorophores are on different sides of the cell membrane, they cannot interact, i.e. FRET does not occur. In an ideal case, this construct would show 100 % co-diffusion of the green and red fluorophore. Figure 5B-D shows two measurements of the NT-SNAP in two cells on two different measurement days. Fitting the *ACF*_*gp*_ and *ACF*_*rd*_ of the “better” measurement shown in Figure 5B with eq. 7 and the *CCF*_*PIE*_ with eq. 9, reveals 50-60 molecules in focus for the *ACF*_*gp*_ and *ACF*_*rd*_, whereas *N*_*app*_, thus 1/G_0_(*t*_*c*_) ∼ 114 for the *CCF*_*PIE*_ (Figure 5C). The concentration of labeled receptors lies in the ∼100 nM range as determined with eq. 8. To determine the average concentration of double-labeled molecules, first the ratio of G_0_(*t*_*c*_) (represented by 1/*N*_*(app)*_) of the *CCF*_*PIE*_ to *ACF*_*gp*_ and *ACF*_*rd*_, respectively, is calculated (eq. 6). Next, these values, *r*_*GRcell*_ = 0.43 and *r*_*RGcell*_ = 0.53 are compared to the values obtained from the DNA measurement (*r*_*GR,DNA*_ = 0.51 and *r*_*RG,DNA*_ = 0.79 on this measurement day). Using the rule of proportions, a *r*_*GRcell*_ = 0.43 from the *ACF*_*gp*_ of the eGFP signal reflects a fraction of co-diffusion (*r*_*GRcell*_/*r*_*GR,DNA*_) of 0.84, where for the other case of *ACF*_*rd*_ of the SNAP substrate signal this value amounts to 0.67. The average concentration of double-labeled NT-SNAP construct can finally be calculated based on eq. 10. In contrast, in the measurement shown in Figure 5D from a different day the concentration of receptors is quite low and the data very noisy such that the fit range is limited up to ∼ 10 µs and also low amount of co-diffusion is observed (15 – 26 %).

**Figure 5.**
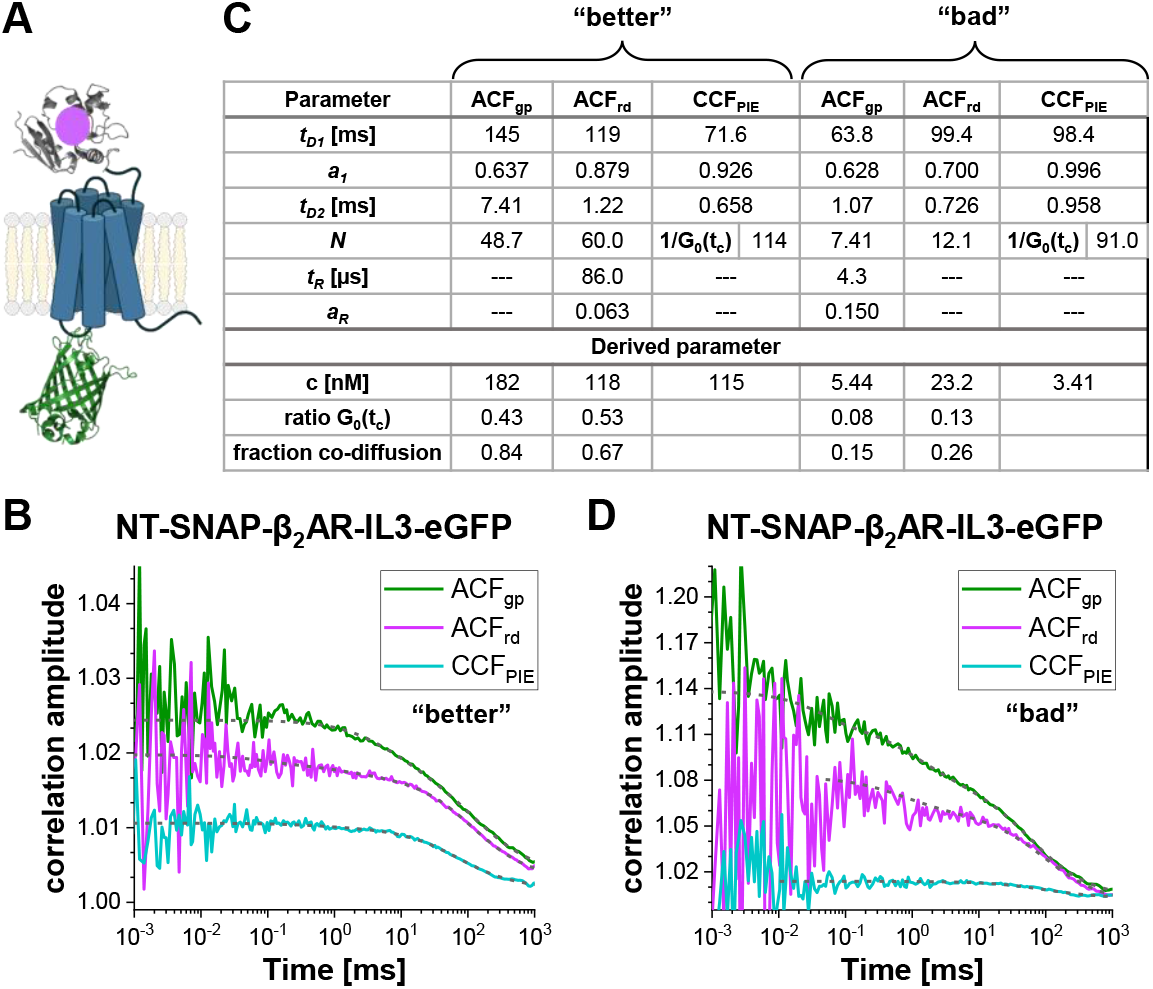
Double-labeled NT-SNAP-β_2_AR-IL3-eGFP construct. (A) In the double labeled construct, the eGFP is inserted into the intracellular loop 3 and the SNAP tag attached to the N-terminus of β_2_AR (NT-SNAP). (B, D) *ACF*_*gp*_, *ACF*_*rd*_ and *CCF*_*PIE*_ of two measurements of the double-labeled construct. The data is fit to a bimodal two-dimensional diffusion model (**eq. 9**, *CCF*_*PIE*_) and including additional relaxation terms (**eq. 7**, *ACF*_*gp*_ and *ACF*_*rd*_). The table in panel (C) shows the fit results and the derived parameter concentration (**eq. 8**), the ratio of the correlation amplitude at zero correlation time (G_0_(t_c_)) and the fraction of co-diffusing molecules (**eq. 10**). Please note that the measurement were acquired on different days, thus slightly different factor for the amplitude correction were used (B: *r*_*GR,DNA*_ = 0.51 and *r*_*RG,DNA*_ = 0.79; D: *r*_*GR,DNA*_ = 0.51 and *r*_*RG,DNA*_ = 0.56).

#### 3.3. Double-labeled construct undergoing FRET: β_2_AR-IL3-eGFP-CT-SNAP

In the double-labeled construct β_2_AR-IL3-eGFP-CT-SNAP (Figure 6A), the eGFP is inserted into the intracellular loop 3 identical to the NT-SNAP-β_2_AR-IL3-eGFP construct and the SNAP tag attached to the C-terminus. Thus, here both labels are on the same side of the cells’ plasma membrane, and due to the fluorophores’ closeness FRET is occurring as can be judged based on the quenched eGFP lifetime (Supplementary Note 5). Considering the general flexibility of relatively unstructured protein regions like the C-terminus ^34^ and the knowledge about the at least two different proteins conformations of GPCRs ^35^, dynamic changes in the FRET efficiency due to eGFP-SNAP distance changes would be expected and to show up as anticorrelated terms in the *CCF*_*FRET*_ (orange curve in Figure 6B). The joint (or global) fit of all five correlations shown in Figure 6B reveals ∼70 % of slowly diffusing molecules at ∼100 ms while the rest diffuses with ∼ 1 ms. All autocorrelations and *CCF*_*FRET*_ show relaxation terms at 37 µs and 3 µs, where the correlations dominated by red signal (*ACF*_*rp*_, *ACF*_*rd*_ and *CCF*_*FRET*_) show an additional slow component ∼ 50 ms (Figure 6C).

**Figure 6.**
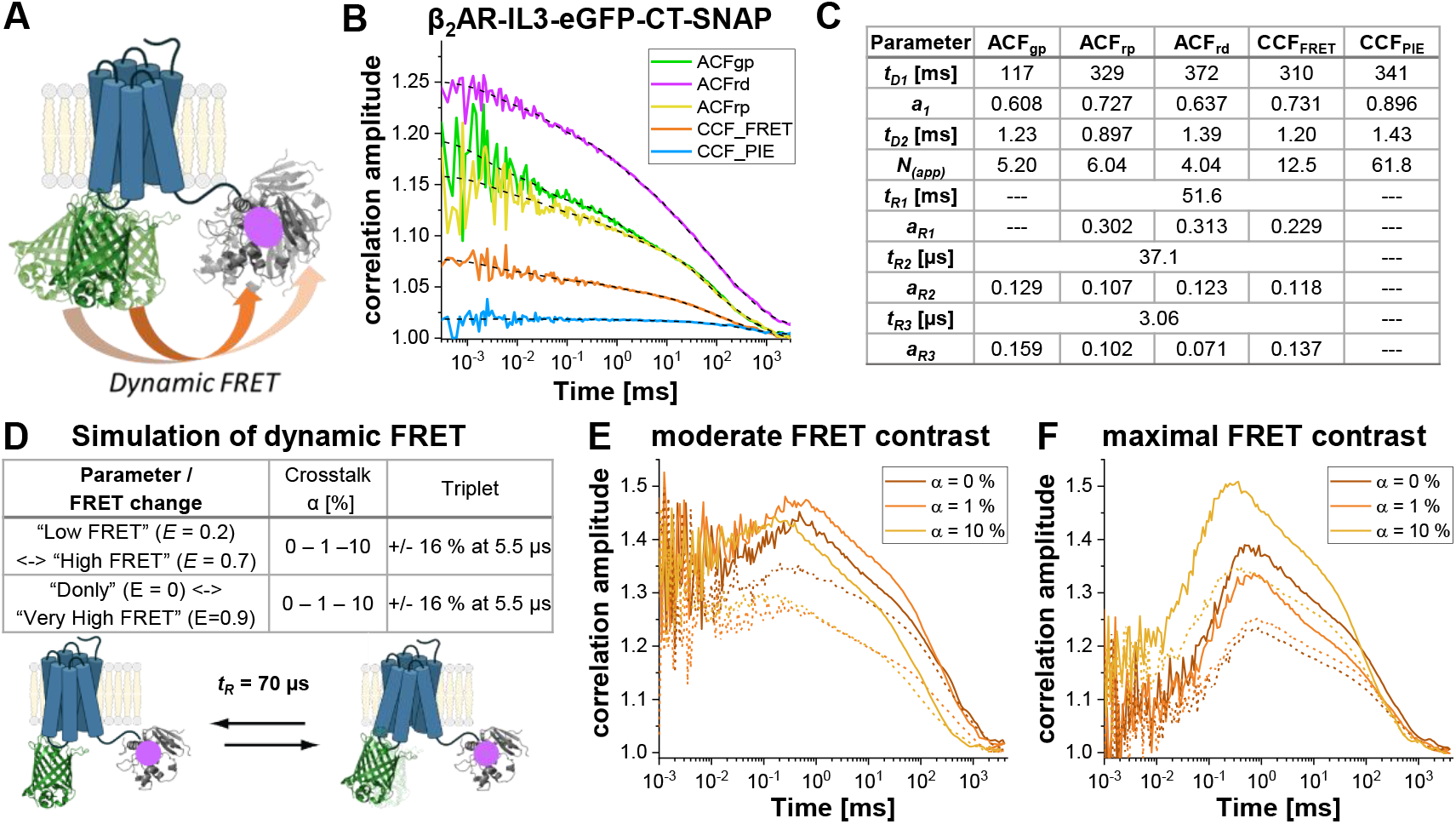
Simulation of double-labeled sample showing dynamic FRET. (A) Double-labeled β_2_AR with an eGFP inserted into the intracellular loop 3 and a C-terminal SNAP tag. Both fluorophores are close enough to undergo FRET and show changes in the FRET efficiency if the receptor undergoes protein dynamics. (B) Autocorrelation (*ACF*_*gp*_, *ACF*_*rp*_ and *ACF*_*rd*_, fit with **eq. 7**) and cross-correlation curves (*CCF*_*FRET*_ (**eq. 7**) and *CCF*_*PIE*_ (**eq. 9**)) of an example measurement. Table in panel (C) shows the fit results. (D-F) To show the influence of experimental parameter on the expected, FRET-induced anticorrelation term, 12 simulations were performed, in which the change in the FRET efficiency (small or large), different amount of donor crosstalk into the acceptor channels (0 %, 1 % or 10 %) and the absence and presence of triplet blinking were modeled. The equilibrium fraction of both FRET-states was assumed to 50:50 and their exchange rates adjusted such that the obtained relaxation time *t*_*R*_ = 70 µs. More details on the simulations see in the text. (E) *CCF*_*FRET*_ of the simulation results with a moderate FRET contrast and in the absence of crosstalk (dark orange), 1 % crosstalk (orange) and 10 % crosstalk (light orange). Solid lines show results in the absence of triplet, dashed lines in the presence of triplet. (F) *CCF*_*FRET*_ of the simulation results with maximal FRET contrast. The color code is identical to (E).

Next, to show the effect of FRET-induced changes on the *CCF*_*FRET*_ under different conditions, a series of simulations were performed (Figure 6D) assuming a two state system exchanging with a rate constant at 70 µs. If a molecule switches between a low and a high-FRET (LF, HF) state – inducing anticorrelated increase in the red signal and a decrease in the green signal - at a timescale faster than its residence time in the focal volume, the photons coming from LF versus HF are anticorrelated as the molecule can only be in HF or LF, but never be in both at the same time ^6,31,36^. Here, two different FRET-systems were assumed showing either a moderate change in FRET efficiency between the two states (LF<-> HF) or maximal FRET contrast (DOnly <-> VHF) in the absence or presence of triplet blinking and increasing amount of donor crosstalk into the red channels. The simulations were performed using Burbulator ^37^. The diffusion term was modeled as a bimodal distribution with 30 % of fast diffusing molecules at *t*_*D1*_ = 1 ms and the rest of the molecules diffusing slowly with t_*D2*_ = 100 ms. In total, 10^7^ photons were simulated in a 3D Gaussian shaped volume with *w*_*0*_ = 0.5 µm and *z*_*0*_ = 1.5 µm, a box size of 20, and *N*_*FCS*_ = 0.01.

Figure 6E-F shows the simulation results for the FRET-induced cross-correlation *CCF*_*FRET*_ for moderate (Figure 6E) and maximal FRET contrast (Figure 6F) in the absence (solid lines) and presence of triplet blinking (dashed lines). The FRET-induced anti-correlation can easily be seen in Figure 6F and also the “dampening” effect upon addition of an additional triplet state reducing the correlation amplitude is clearly seen (Figure 6E-F) ^38,39^.

However, in the simulation case most similar to the experimental conditions (α = 10 %, 15 % triplet blinking and moderate FRET contrast, dashed yellow line in Figure 6E) the anticorrelation term is nearly diminished. Figure 7 shows the result of analyzing this simulated data using the information encoded in the photon arrival time histograms (i.e. the fluorescence lifetime) by means of Fluorescence Lifetime Correlation Spectroscopy (FLCS) ^17,19^ or species-filtered FCS (fFCS) ^18^. Here, the fluorescence lifetime of the known HF and LF species (Figure 7A) is used to generate weights or “filters” (Figure 7B) which are applied during the correlation procedure. In the obtained species-auto- and crosscorrelation curves (Figure 7C-D) the anticorrelation is clearly seen.

**Figure 7.**
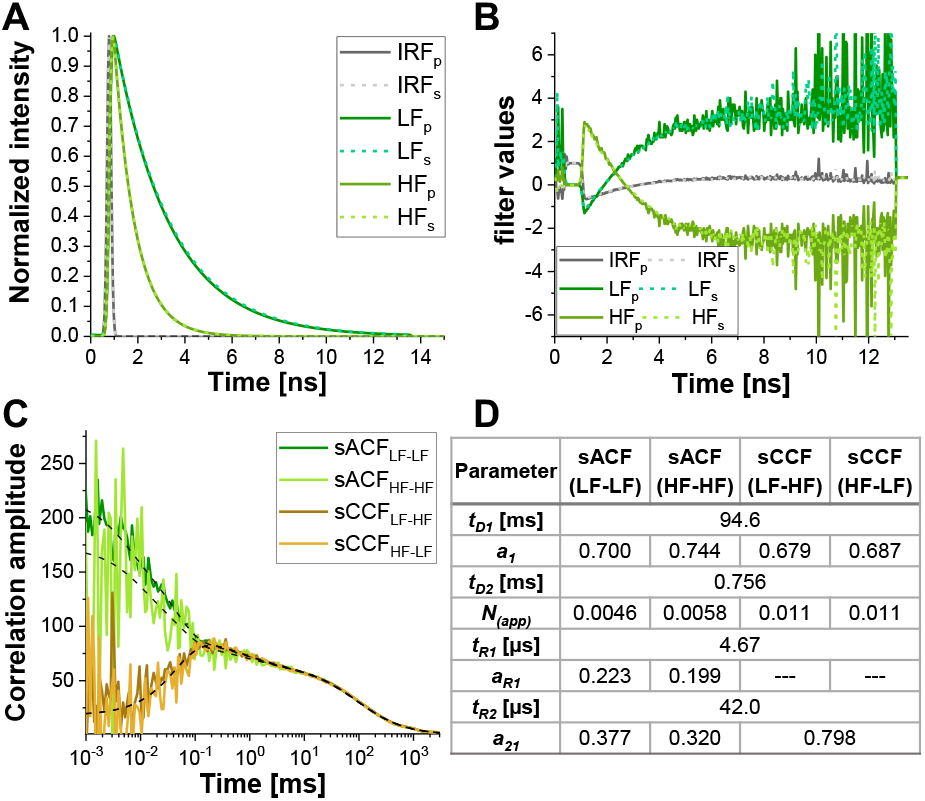
Lifetime-filtered FCS can help to uncover the protein dynamics based fluctuations in FRET efficiency in samples with high crosstalk, significant triplet blinking or other photophysical or experimental properties masking the FRET-induced anticorrelation in the *CCF*_*FRET*_. Here, the approach is shown exemplary for the data shown in Figure 6E for the simulation containing 10 % crosstalk and 5 % triplet blinking. (A) Normalized fluorescence intensity decay patterns for the two FRET-species (light and dark green for high and low FRET, respectively) and the IRF (grey). The pattern for the parallel detection channel is shown in solid lines, dashed lines for the perpendicular detection channel. (B) The weighting function or “filter” were generated based on the patterns shown in (A), color code is identical to (A). Please note that only the signal in the green detection channels, and thus the FRET-induced donor quenching, is considered here. (C) Four different species-selective correlations are obtained: species-autocorrelations of the low FRET state (*sACF*_*LF-LF*_, dark green) and the high FRET state (*sACF*_*HF-HF*_, light green), and the two species-crosscorrelations between the low FRET to the high FRET state (*sCCF*_*LF-HF*_, dark orange) and vice versa (*sCCF*_*HF-LF*_, orange). The sCCF clearly shows the anticorrelation in the µs-range. Dashed black lines show the fits. sACF were fit with **eq. 9** and sCCF with **eq. 11**. Table in panel (D) shows the fit results.

### 4. *CCF*_*PIE*_ amplitude to study Protein-Protein Interaction (PPI)

Finally, a common use case for PIE-based FCS in live cells is to study the interaction between two different proteins. Here, the read-out parameter is the amplitude of the *CCF*_*PIE*_, or more precise the ratio of the autocorrelation amplitudes *ACF*_*gp*_ and *ACF*_*rd*_ to the amplitude of *CCF*_*PIE*_. To exemplify the effect of increasing co-diffusion on *CCF*_*PIE*_, simulations have been performed based on the two single-labeled constructs, β_2_AR-IL3-eGFP and NT-SNAP-β_2_AR, described above (Figure 8A). Figure 8B shows how the amplitude of *CCF*_*PIE*_ increases when the fraction of co-diffusing molecules change from 0 % to 100 %. Please note that the diffusion components were modeled as above and that a 1 % crosstalk of green signal into the red channels in the delay time window was added.

**Figure 8.**
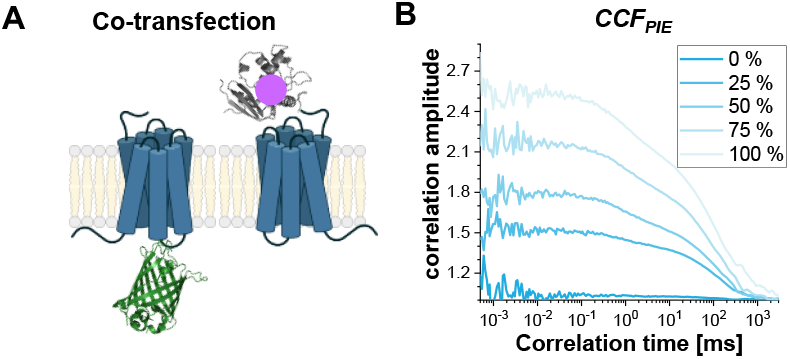
The *CCF*_*PIE*_ can be used to study the interaction of two proteins. (A) Here, a co-transfection study of β_2_AR-IL3-eGFP with NT-SNAP-β_2_AR (carrying a “red” SNAP-label) was simulated. (B) For an increasing amount of co-diffusing molecules (0 % (dark blue) -> 100 % (light blue)) the amplitude G(*t*_*c*_) increases. The diffusion term was again modeled as a bimodal distribution with 30 % of fast diffusing molecules at *t*_*D1*_ = 1 ms and the rest of the molecules diffusing slowly with *t*_*D2*_ = 100 ms. Additionally, 1 % crosstalk of green signal into the red delay time window was added.

## Discussion

The use of FCS techniques in GPCRs allows the mobility and interactions of receptors inside the cell to be assessed ^40^. The advantage of FRET-FCS technique is that, along with mobility we can investigate the conformational dynamics of GPCRs. However, performing FRET-FCS in live cells is challenging and requires good transfected (not overexpressed) cells, a well-calibrated setup and a good pipeline to analyze the data globally. Here, first the critical points in the sample preparation and experimental procedure are discussed concerning all, the biological, spectroscopic and technical point of view.

Critical experimental steps include the minimization of background and autofluorescence (by using extensively cleaned coverslips and phenol-red free media), the optimization of transfection conditions (e.g. amount of plasmid DNA and time after transfection) to achieve low expression levels and efficient labeling. Of course, it is also vital to assure that the function of the labeled protein is not hampered. Thus, in live cell experiments, the decision for the labeling strategy and label position is often made in favor of fluorescent proteins or SNAP/CLIP tag attached to the flexible N- or C-terminus ^41,42^. Alternative labeling strategies like the insertion of an unnatural amino acid with a reactive side chain for labeling with an organic fluorophore are emerging in the last years ^43^.

For dual-color PIE-FCS, where solely the interaction of two molecules of interest are to be investigated, the fluorophores can be selected from quite a pool of available ones. Here, spectroscopy-wise the goal should be select a pair such that little crosstalk and direct acceptor excitation occur. Additionally, the fluorophores should be photostable and show little or no bleaching under the chosen experimental conditions. It is recommend selecting rather fluorophores in the red spectral range as (1) the autofluorescence background from the cell is reduced and (2) the excitation light is of longer wavelength, thus less phototoxic^14^. Photobleaching can be minimized by conducting a so-called “powerseries” first, in which the laser power is increased stepwise and the molecular brightness is observed. The optimal excitation intensity range lies in the linear range of the results^44^.

If the two labels are supposed to report also on protein conformational dynamics, either in a protein assembly or by being placed inside the same protein, by FRET the choice of available fluorophore is more restricted. Here, the possible minimal / maximal distance between the two fluorophores should be estimated beforehand e.g. based on available structures or molecular size, and fluorophore pair selected with a reasonable Förster radius *R*_*0*_ such that FRET can actually occur ^20^.

Here, eGFP and a SNAP tag were chosen for labeling and the SNAP tag was labeled with either an intracellular or a membrane-impermeable surface substrate. The spectra are similar to the ones shown in Figure 2C-D. This combination shows high crosstalk and direct acceptor excitation, and results in significant “false” signal in the red channels in the prompt time window. Ideally, both values should not exceed 5 % ^5,6,38^. However, with a Förster radius of 57 Å it is ideally suited to probe the distance between the labels in the β_2_AR-IL3-eGFP-CT-SNAP construct as can be evaluated from the quenched eGFP lifetime (Supplementary Note 5).

Technically, - of course – as for any fluorescence spectroscopy experiment, the device should be well aligned and possess suitable excitation sources, emission filter and sensitive detectors. To avoid artifacts from detector afterpulsing on the µs timescale, at least two detectors of each color should be present, which can be cross-correlated. In modern counting electronics, dead time of the detection card in the ns time range hardly plays a role due to the independent routing channels, however, it might be checked ^16^ and for high time resolutions each detection channel should be doubled, i.e. four detector per color should be used, to also bypass detector dead times ^2,15,29,45^. If the fluorescence lifetime and distance between the fluorophores are to be analyzed, the emission has to be collected polarization-dependent as is the case here (Supplementary Note 1). Finally, in PIE experiments, the distance between the prompt and delay pulse is critical and should be chosen such that the fluorescence intensity of the fluorophores has been largely decayed (Figure 1B). A common rule is to place the two pulses 5x the fluorescence lifetime apart, i.e. for eGFP with a fluorescence lifetime of 2.5 ns the distance should be 12.5 ns at minimum ^22^.

After having detailed all consideration for the experimental procedure, the data and its analysis is discussed in more detail. As already mentioned in the protocol section, the alignment of the setup must be checked daily, including the analysis of the calibration measurements. The data shown in Figure 2A-C e.g. shows an additional relaxation component in 8-40 µs. Typical triplet blinking of the green calibration fluorophore is known to occur in the 2-10 µs range ^13,15,46^. The slow relaxation component required in all curves of the DNA sample (Figure3C) – too slow for actual triplet blinking – might stem from interactions of the DNA with the fluorophores ^39^. However, this component would not be expected in *CCF*_*PIE*_, and most likely stems from residual crosstalk. Thus, it is highly advisable to perform the analysis of the calibration samples directly prior to proceeding to the cell experiments to judge the quality of the day’s alignment.

The proper calibration of the confocal overlap volume requires a sample with 100 % co-diffusion of the green and red label. Here, commonly DNA double-strands are used. Single DNA strands with fluorophores inserted can be bought tailored to the specific fluorescence properties required and annealed with high yield to the DNA double strands with a specific fluorophore spacing. However, Good Laboratory Practice advises to check the integrity and labeling degree of the DNA strands by agarose gel electrophoresis and measuring the absorption spectrum. Also, the yield of the double-strand assembly should be checked as this DNA calibration measurement critically relies on the fact that the assumption of a 100 % co-diffusion of the green with the red label is valid, else correction factors might have to be applied ^16,22^. In the calibration measurements shown in Figure 2 and Figure 3, a detection volume of 1.4 fL and 1.9 fL in the green and red channel was obtained. This size difference is expected for a setup with nearly diffraction-limited excitation volumes (Supplementary Note 2), where the size of the excitation volume scales with the excitation wavelength. This in turn explains the different correlation amplitudes observed in Figure 3B. The derived correction factors *r*_*GR*_ = 0.56 and *r*_*RG*_ = 0.72 correct for this size discrepancy and potential non-perfect overlap of the two excitation volumes ^3,4^.

Figure 4-7 showcase the exemplary workflow of a PIE-F(C)CS based study aimed toward understanding the conformation protein dynamics. First, the two single-labeled constructs β_2_AR-IL3-eGFP and NT-SNAP-β_2_AR serve as controls to characterize the fluorophore properties in cells in the absence of the respective other fluorophore (Figure 4). Next, the double-labeled construct NT-SNAP-β_2_AR-IL3-eGFP carries a SNAPtag facing the cell outside and an eGFP on the cytoplasmic side. It serves as a “100 % co-diffusion” control (Figure 5).

The last construct, β_2_AR-IL3-eGFP-CT-SNAP, carries both fluorophores on the cytoplasmic side and close enough together to undergo FRET. Here, again a 100 % co-diffusion would be expected next to anti-correlated intensity fluctuation in the green and red channels signal in the prompt time window, i.e. after donor excitation, due to protein dynamics influencing the FRET efficiency ^31-33^ (Figure 6-7).

All GPCR β_2_AR constructs show bimodal diffusion on the cell membrane (Figure 4A). Whereas the β_2_AR-IL3-eGFP shows only the expected triplet blinking (Figure 5B) ^13,15^, NT-SNAP-β_2_AR shows an additional slow relaxation time (Figure 5C-D). It is likely that *t*_*R2*_ might stem from unbound SNAP substrate. This could be elucidated by further experiments, e.g. by also measuring the diffusion and photophysical properties of the used SNAP substrate free in solution. Of note, a straightforward experiment to differentiate between diffusion and relaxation times is to change the pinhole of the confocal setup, i.e. increasing the effective volume: While the diffusion times increase with increasing effective volumes, relaxations term are unaltered ^13^. When determining the concentration of fluorescent protein (FP) based on the fit results, be aware that FPs in general undergo a maturation process, in which finally the chromophore is formed ^12^. This maturation time may differ from FP to FP in addition to photophysics that depends on the local chemical environment ^13,15^. Thus, the actual protein concentration present in the sample reported by FCS is usually underestimated, which can be corrected, if the fraction of non-fluorescent FPs can be determined in the experiment. Finally, it is advisable to check the fluorophore spectra in live cells – if possible - as most fluorophores react sensitive to their environment ^13,15,46^, and to correct the values for α and δ, if required. Here, the background to subtract is determined by the signal collected in non-transfected cells. Additionally, the autocorrelation of the respective other color channel and the *CCF*_*PIE*_ should be checked to be able to identify false signals (Supplementary Note 4 – Figure 30).

The two measurements from the NT-SNAP-β_2_AR-IL3-eGFP (Figure 5D), where the fluorophores are located on different sides of the membrane, were acquired on different days and show very different results. One has a high degree of labeling and due to averaging of measurements relatively lower noise (Figure 5B), while from the other cell, only two measurements could be collected and it shows quite high noise (Figure 5A). This emphasizes the fact to collect sufficient amount of data – from a single cell and from different cells on multiple independent sample preparations, to critically evaluate the results timely and to optimize the labeling strategy, especially when working with labeling substrates. Be aware that here no FRET can occur due to the large distance between the fluorophores. This can also be cross-checked by evaluating the unaltered eGFP fluorescence lifetime (Supplementary Note 5). In the β_2_AR-IL3-eGFP-CT-SNAP construct (Figure 6A), FRET occurs as can be evaluated from the quenched eGFP lifetime (Supplementary Note 5). However, no anticorrelation term is visible (Figure 6B). Up to three additional relaxation term are required in *ACF*_*gp*_, *ACF*_*rp*_, *ACF*_*rd*_ and *CCF*_*FRET*_ (Figure 6C). The slow component in *ACF*_*rp*_, *ACF*_*rd*_ and *CCF*_*FRET*_ might be due to acceptor bleaching and of course influences the obtained value of the slow diffusion found in these curves (∼350 ms compared to 117 ms in *ACF*_*gp*_). *t*_*D*_ in red is supposed to be slightly larger than in green due to the differently sized confocal volumes (Figure 2) – but only by a factor comparable to the size difference. The very fast relaxation time of 3 µs reflects the triplet blinking of the fluorophores ^13,15,46^, where the slower one of 37 µs might be due to FRET: Similarly as FRET induces an anticorrelation in the *CCF*_*FRET*_, positive correlations are expected in the autocorrelations ^31-33^. The presence of this term as “positive” in *CCF*_*FRET*_ and at all in the *ACF*_*rd*_ might be explained with the high crosstalk and should be further elucidated. Note that the *CCF*_*PIE*_ is flat as expected.

On other terms it should be noted that the occurrence of FRET in a system of interest leads to non-linear effects on the correlation curves ^6^. The molecular brightness e.g. of a molecule scales into the correlation amplitude squared and each FRET-state (and the always present molecules without an active receptor) shows different molecular brightness. Indeed, FRET decreases the apparent concentration of green molecules detected (i.e. increases *ACF*_*gp*_ amplitude) and the number of red molecules (determined from red-prompt) is overestimated ^5^. Both effects influence the amount of interaction derived from both *CCF*_*FRET*_ and *CCF*_*PIE*_. However, global analysis shown e.g. for the intramolecular dynamics of Calmodulin ^31,32^ or Syntaxin ^33^ can reveal the protein dynamics. When carefully calibrated, the average FRET efficiency may be extracted from the relative *CCF*_*PIE*_ and ACF amplitudes ^22^, whereas the limiting states might be determined from the analysis of the donor fluorescence lifetime distribution ^33^.

Considering the fact that in live cell experiment with large fluorophores like eGFP the FRET contrast is likely to be even lower and that the direct excitation of the acceptor was not added in the simulation, might explain why the identification of the anticorrelation in live cell experiments provides a challenge. A promising analysis alternative relies on harvesting the information encoded in the photon arrival time histograms (Figure 1B) accessible due to the time-correlated single photon counting data collection ^29,30^. If the fluorescence lifetime (∼patterns) of the two (or more) (FRET) species inside the sample are known (Figure 7A), “filter” or weights can be constructed which are applied during the correlation process (Figure 7B) ^17-19^. Obtained are the correlation curves, which no longer represent the correlation of detection channels but rather the auto- or crosscorrelations between two different (FRET) species, thus renamed to species-ACF (sACF) or species-CCF (sCCF). Applying this approach to the simulated data with moderate FRET contrast, high crosstalk and triplet blinking recovers the anticorrelation term (Figure 7C-D). However, it should be noted that relaxation times can be obtained but the relationship to amplitude are lost ^18^. This approach has been applied previously in live cell experiment e.g. to study the interaction of EGFR with its antagonist ^47^ or to separate the fluorescence from proteins attached to a short and long lived eGFP variant ^48^.

While PIE-based FRET measurements in purified proteins are largely used to study protein dynamics ^36 22^, in live cells it focuses on understanding protein-protein interactions. This approach has been applied e.g. to study the regulation MAP kinase activity in yeast ^49^ or to resolve the interaction of membrane proteins with their cytosolic binding partner as summarized in this recent article ^50^. Here, complications may arise e.g. when significant crosstalk of green fluorophores is still present in the delay time window of the red channels or red signal in the green channels in the prompt time window. The former might be caused by an insufficient delay of the red pulse with respect to the green pulse while both effects stem from too strongly overlapping excitation and emission spectra of the chosen fluorophores. It is recommended to check the respective single labeled constructs carefully and correct for false-positive *CCF*_*PIE*_ amplitudes, especially in cells where (short-lived) autofluorescence might be another complicating factor ^22^.

To conclude, the FRET-FCS approach described here has great potential to understand the interaction between proteins and protein dynamics in live cells under near physiological concentrations. In this protocol, the focus was laid on the required calibration measurements and the necessary quantitative analysis to be performed during live cell measurements. To this end, different live cell measurements were shown complemented with simulations, in which parameter were varied systematically, and the composition of tailored fit models to the specific mobility and photophysical properties of the respective data set was presented. The analysis was performed with open-source software tools with an extensive step-by-step protocol and easy-to-adapt templates. Finally, the technical advancements, and thus the availability of ready-to-buy stable PIE-FCS systems together with the spread of open-source software for data analysis will make this technique easily accessible. Thus, live cell PIE-FCS shows great promise in unraveling live cell protein interaction and dynamics.

## Acknowledgements

This project was supported by the Deutsche Forschungsgemeinschaft (SFB/TR 240, project number 374031971, Project INF) to J. B. and K.G.H.

We thank the Rudolf Virchow Center for financial support and Core Unit Fluorescence Imaging for technical support.

## Disclosures

The authors have no conflicts to declare.

## Table of Material

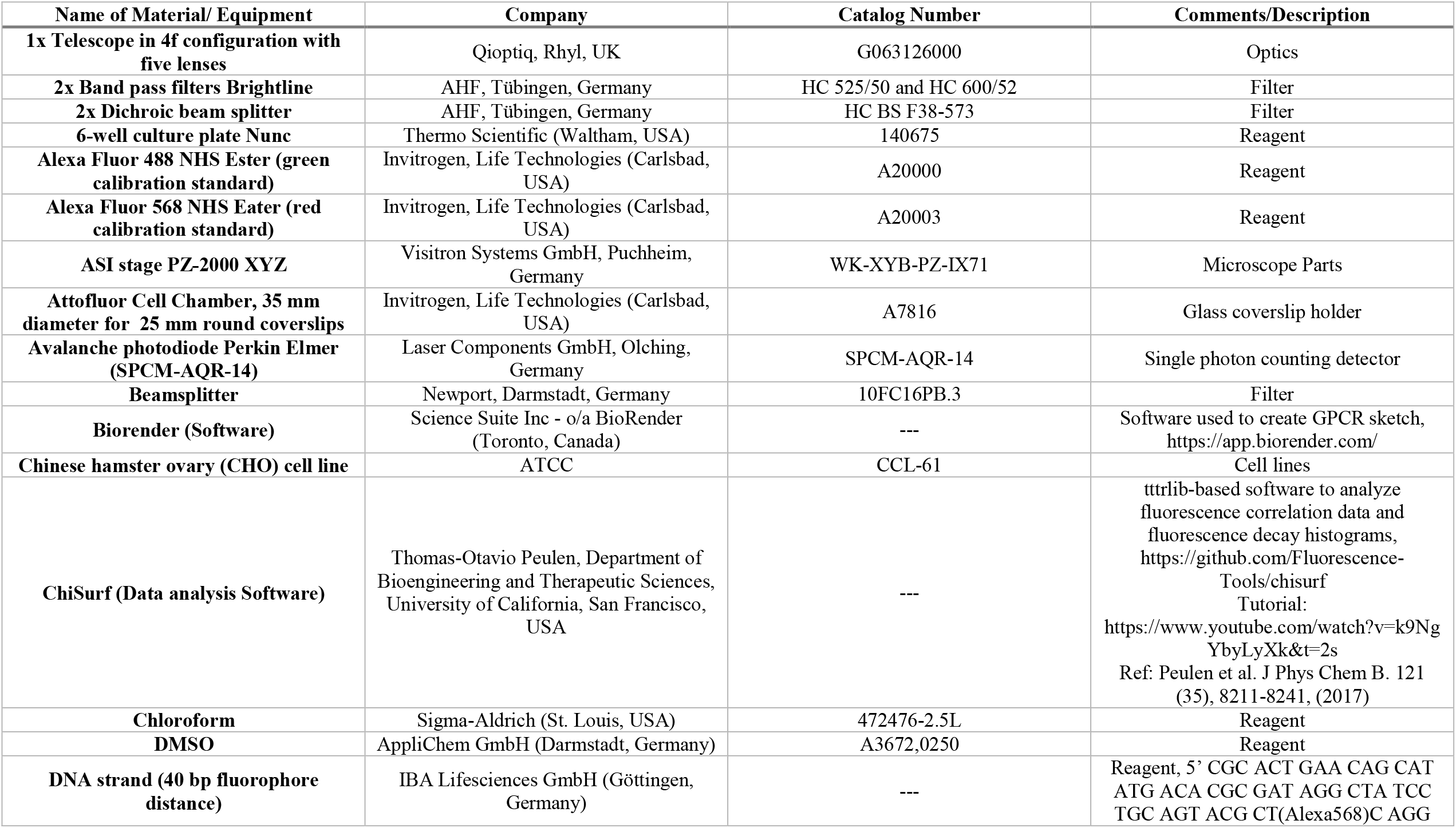
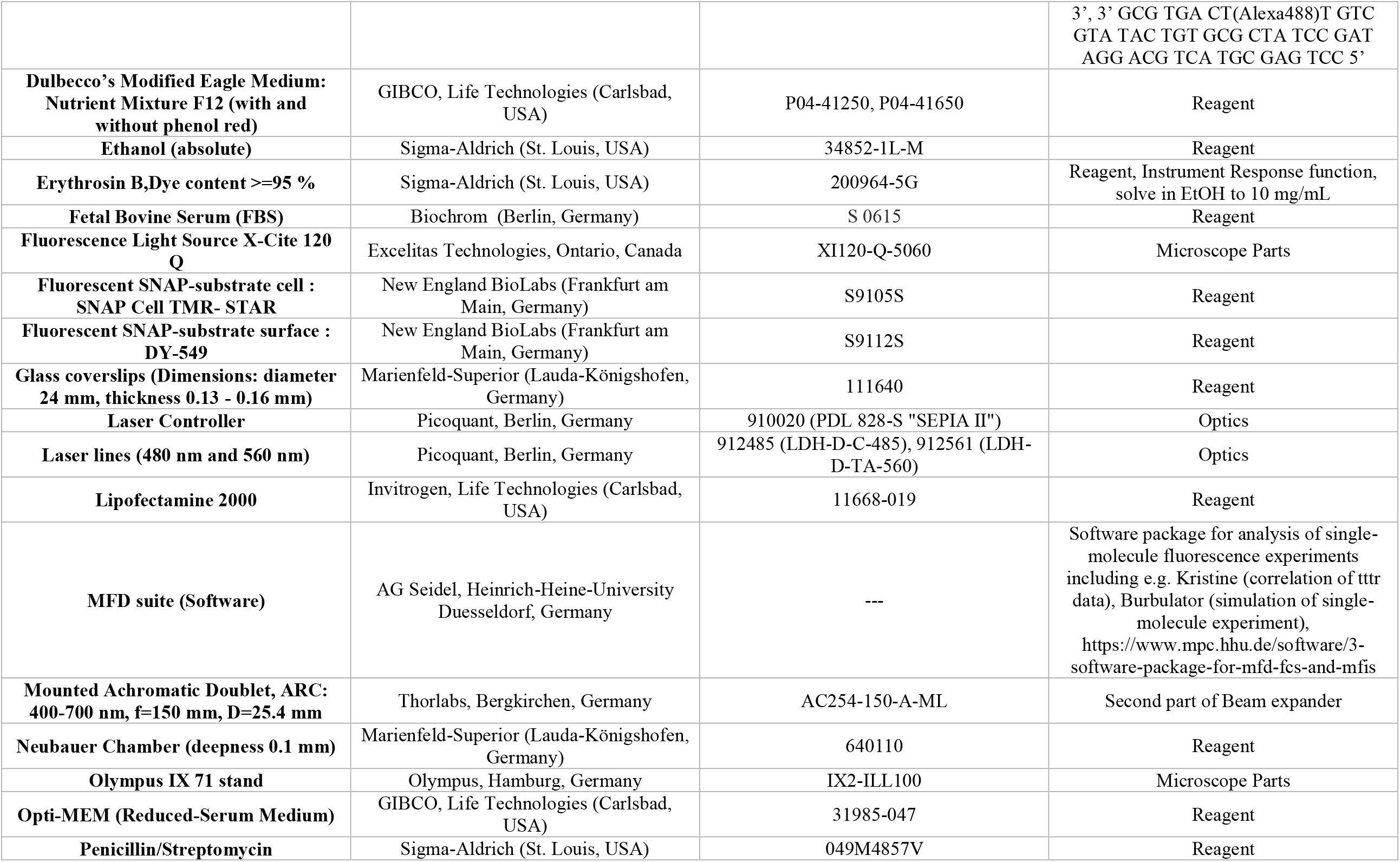
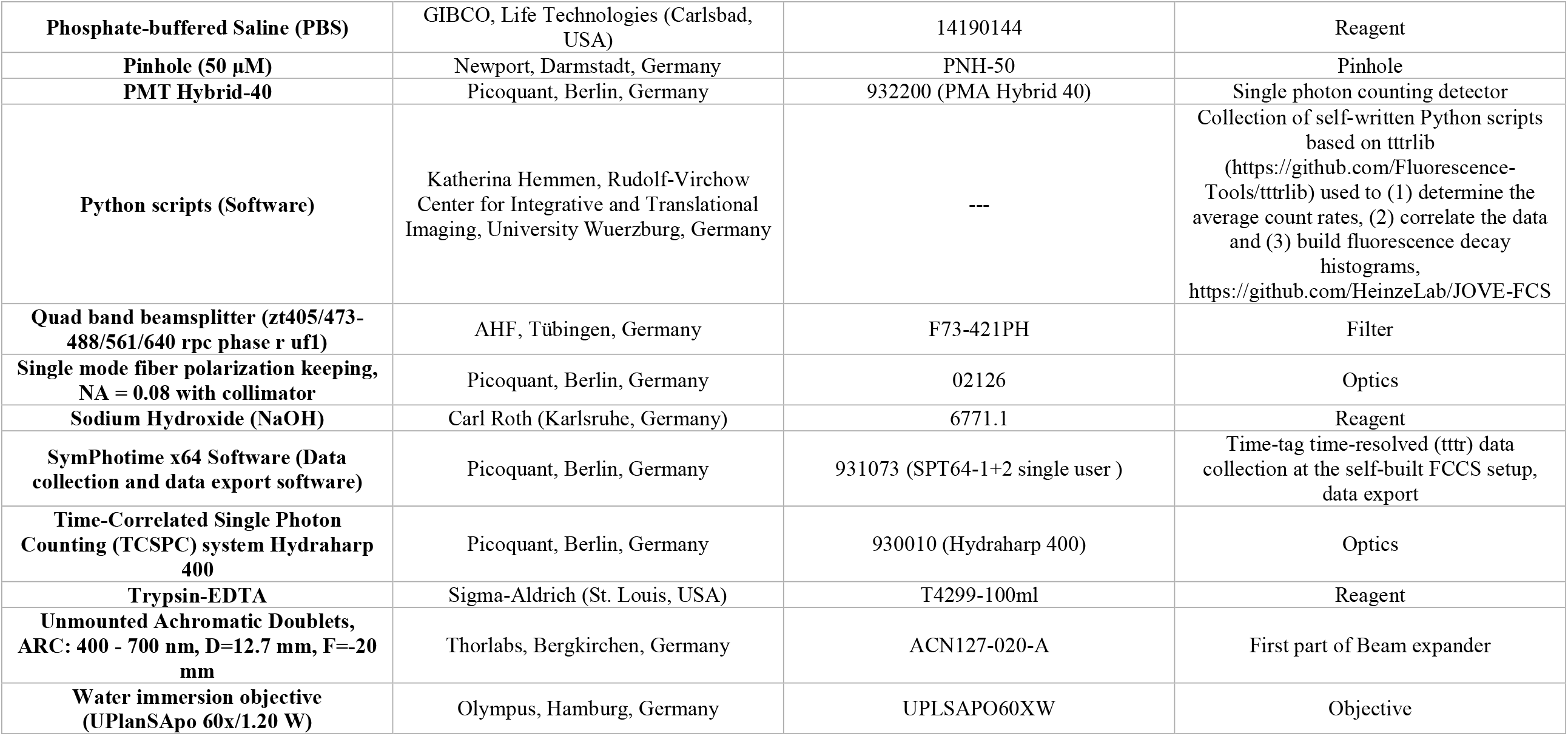

## Notes

https://github.com/HeinzeLab/JOVE-FCS

### Competing Interest Statement

The authors have declared no competing interest.

