## Supplementary File 1-7 for "Dual-color Fluorescence Cross-Correlation Spectroscopy to study Protein-Protein Interaction and Protein Dynamics in Live Cells": SuppNote1_Coverslip cleaning.docx

### 1 Coverslip cleaning for live cell FCS experiments

#### 1.1 Goal

Cleaning of glass coverslips to minimize background signal

#### 1.2 Comments

- Cleaning procedure must be performed under the hood
- Use safety gloves when handling chloroform
- Reuse chloroform and NaOH solution
- Prepare 5 M NaOH solution on ice
- Materials: ultrasonic bath, 2 glass containers, metal rack, small beaker, funnel, 25 ml glass pipet, glass petri dish
- Discard used chemicals according to the local regulations.

#### 1.3 Cleaning Protocol

- Arrange single coverslips in metal rack

| 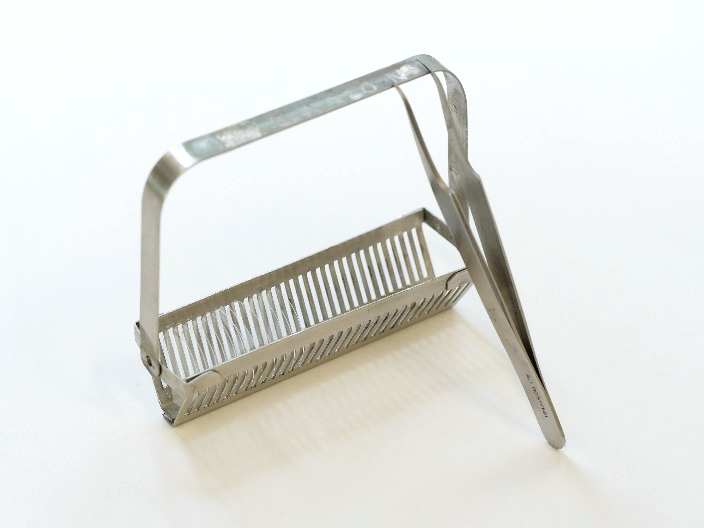 | 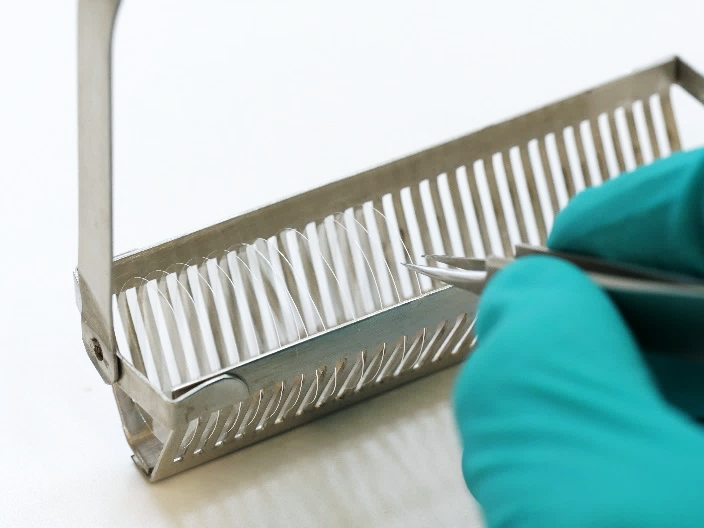 |
| --- | --- |

- Place glass container in ultrasonic bath, fill bath with deionized water
- Hang the rack into the glass container and fix with the loops

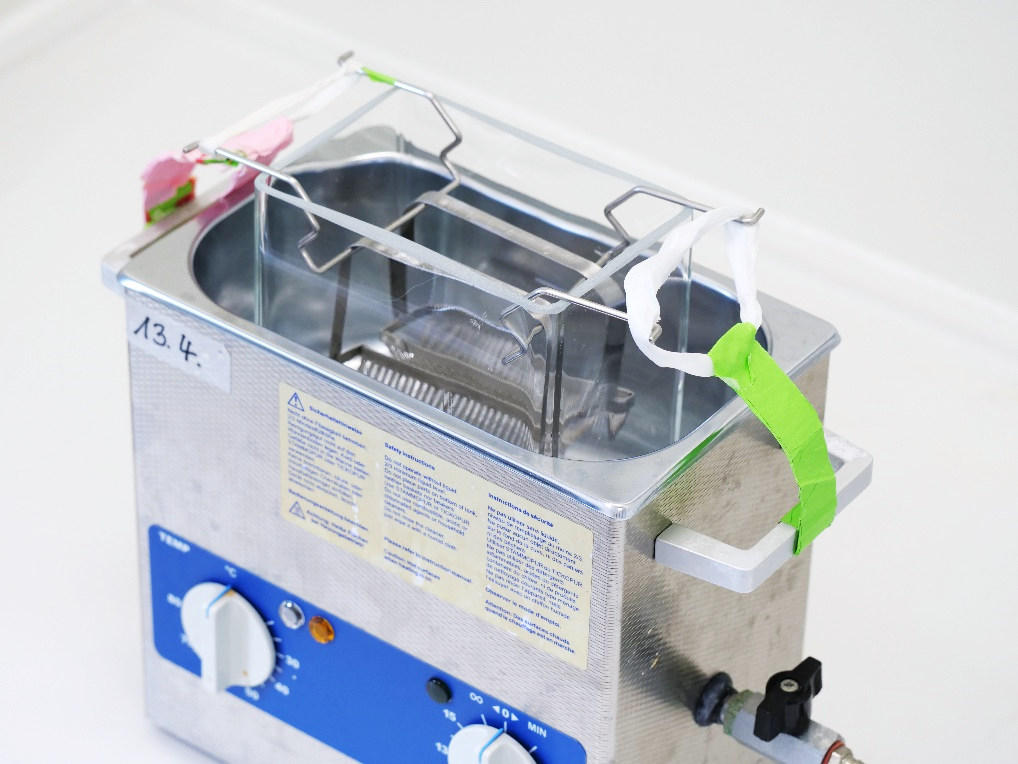

- **Fill the glass container with chloroform and sonicate for 1 hour**
- **Place the rack in an empty glass container and let it dry**
- Put on **safety gloves**!
- Pour the chloroform back into the bottle via a funnel by first scooping it with a beaker, then pipette the rest with a 25 ml glass pipette. Do **NOT** pour it directly out of the glass container! It will spill badly!
- Reinstall the rack in the glass container in the ultrasonic bath.
- **Fill the glass container with 5 M NaOH solution and sonicate for 1 hour**
- **Wash three times in ddH_2_O in a second glass container**
- **Dry glass coverslips**
- **Store in 100 % ethanol in a glass petri dish**

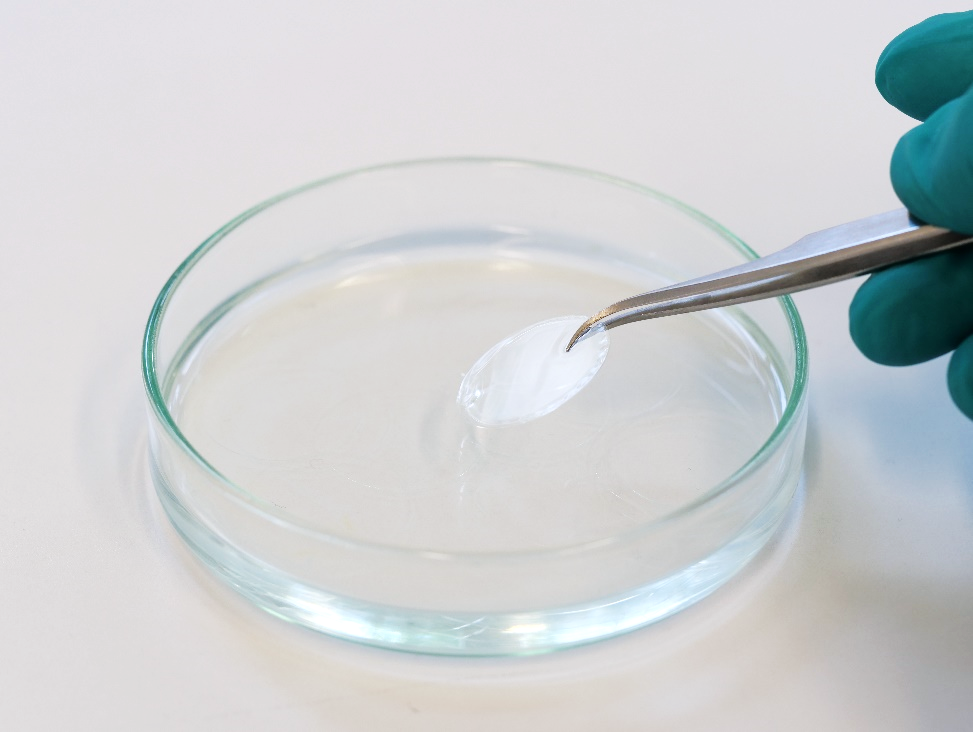

- Pour the NaOH back into the bottle as described before. Clean pipette by pipetting some ddH_2_O.
