## Supplementary File 1-7 for "Dual-color Fluorescence Cross-Correlation Spectroscopy to study Protein-Protein Interaction and Protein Dynamics in Live Cells": SuppNote2_Confocal Setup.docx

### **Confocal FCCS setup**

A custom-built confocal setup based on an Olympus IX 71 stand (Olympus, Hamburg, Germany) equipped with a Time-Correlated Single Photon Counting (TCSPC) system (Hydraharp 400, Picoquant, Berlin, Germany) are used to perform FCS and fluorescence lifetime measurements. Two lasers (488 nm and 560 nm, PicoQuant, Berlin, Germany) are fiber coupled through a single mode fiber (PicoQuant, Berlin, Germany) and expanded via a telescope to fill the back aperture of the objective (60X, water immersion, NA 1.20. Olympus, Hamburg, Germany). A quad band beamsplitter (zt405/473-488/561/640 rpc phase r uf1, AHF, Tübingen, Germany) in the excitation path allows both 488 and 560 nm lasers through the objective to the sample. In the detection path a 50 µm pinhole (PNH-50, Newport, Darmstadt, Germany) rejects out of focus light before being projected on photon counting detectors (2x PMA Hybrid-40, Picoquant, Berlin, Germany and 2x APDs Perkin Elmer SPCM-AQR-14) by a telescope in a 4f configuration (focal length of lenses: 60 mm, G063126000, Qioptiq, Rhyl, UK). The beam was split via a polarizing beamsplitter (10FC16PB.3, Newport, Darmstadt, Germany) into parallel (detector 0 and 1) and perpendicular emission (detector 2 and 3) after the first lens of the telescope. A dichroic beam splitter (HC BS F38-573 Beamsplitter, AHF, Tübingen, Germany) in both parallel and perpendicular path allows the higher wavelength into the APDs (Red detection channels, detector 1 and 3) and reflects the lower wavelength emission into PMTs (green detection channels, detector 0 and 2). Band pass filters Brightline HC 525/50 and Brightline HC 600/52 (AHF, Tübingen, Germany) are used to reject unspecific light in each detection path.
