## Supplementary File 1-7 for "Dual-color Fluorescence Cross-Correlation Spectroscopy to study Protein-Protein Interaction and Protein Dynamics in Live Cells": SuppNote3_Data export.docx

### Data export for PIE-based data analysis

#### Prerequisite for data analysis

In pulsed-interleaved excitation (ref PIE) experiments, the sample is excited by two subsequent laser pulses by two different laser. Commonly the first laser is lower wavelength (“prompt”) and excites the green or donor fluorophore, while the second laser pulse (“delay”) has a longer wavelength and excites the red or acceptor fluorophore. The timing between the two laser pulses is chosen such that the fluorescence lifetime of the fluorophore decays within their specific excitation window.

For a PIE-based data analysis, it is important to know the (time) distance between the two laser pulses and the time resolution, which with the data is saved, as in all data analysis software you will have to specify, which time range should be used in the correlation.

Up to five different correlation can be constructed:

- *ACF_gp_*: Autocorrelation of the green channels in the “prompt” time window
- *ACF_rp_*: Autocorrelation of the red channels in the “prompt” time window
- *ACF_rd_*: Autocorrelation of the red channels in the “delay” time window
- *CCF_FRET_*: Crosscorrelation of the green channels with the red channels in the “prompt” time window
- *CCF_PIE_*: Crosscorrelation of the green channels in the “prompt” window with the red channels in the “delay” time window

Not all correlations are required for each sample.

#### Short list of available software

This list is not complete and only reflects the software we know of.

- Kristine (<https://www.mpc.hhu.de/software/3-software-package-for-mfd-fcs-and-mfis>)
- PAM (<https://gitlab.com/PAM-PIE/PAM>)
- Globals for FCS (<https://www.lfd.uci.edu/globals/>)
- Our self-written scripts for batch export (<https://github.com/HeinzeLab/JOVE-FCS>)
- Proprietary software from the manufacturer of the microscope, e.g. Symphotime (Picoquant).

#### Example data export workflow

##### Symphotime

- Open the project file in Symphotime
  - Caution: The folder needs to contain the ending “.sptw” and must contain a file named “WSLogfile.sptl” to recognized as a project file.

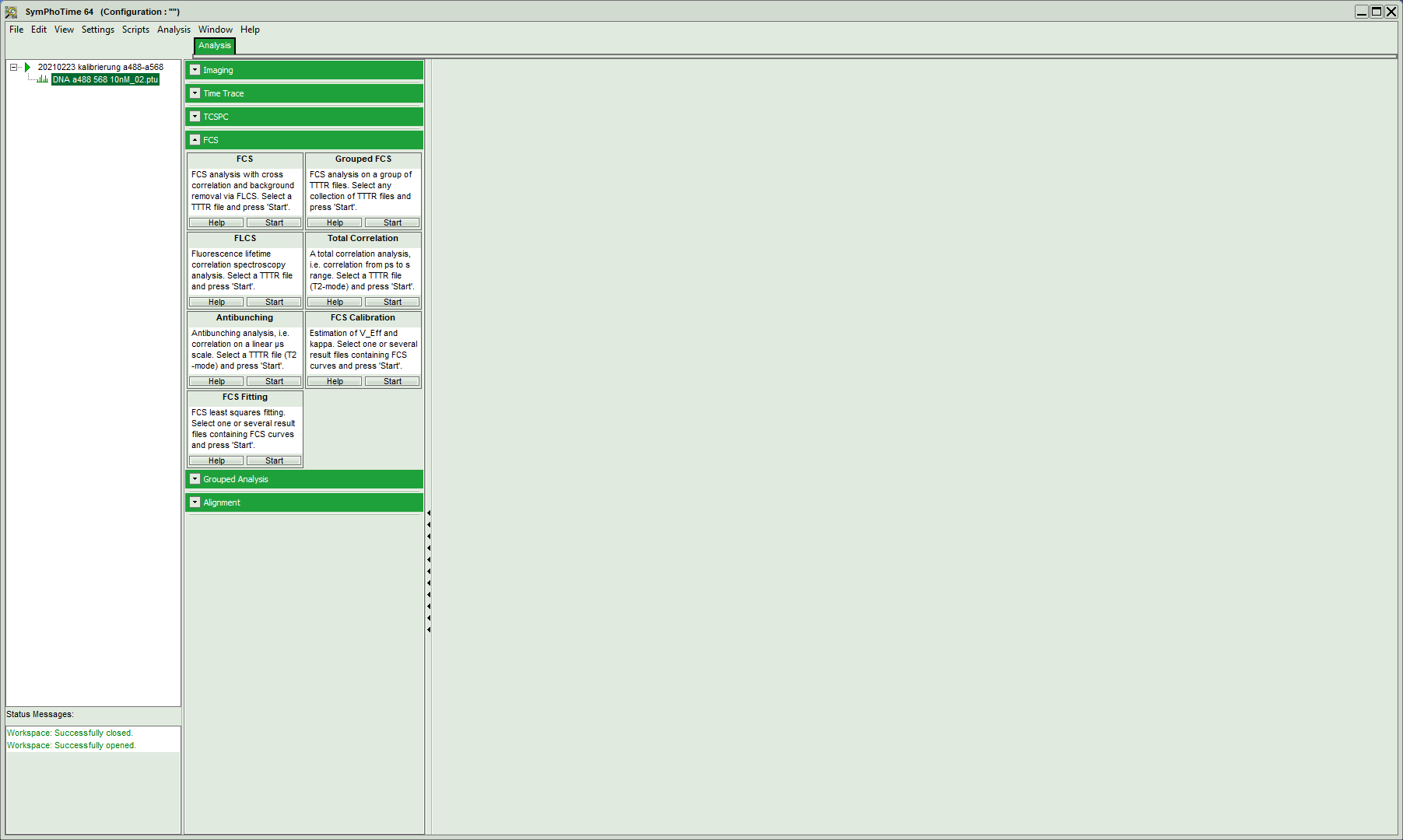

- Start a new “FCS´” and wait until it has opened the data:

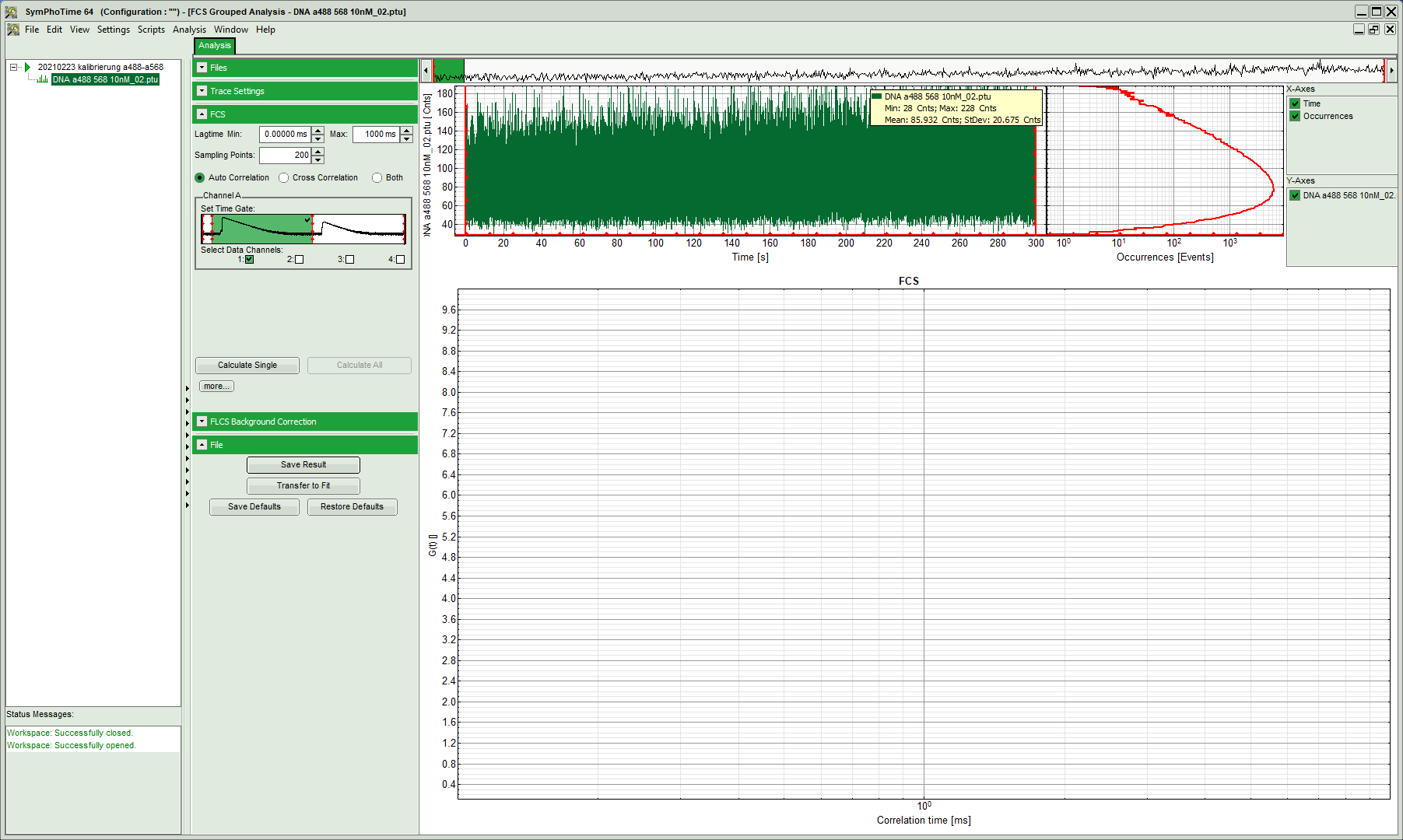

- In the top left corner the correlation setting can be determined:
  - Lagtime “min” and “max”: a minimum of zero means it starts from earliest time point possible, i.e. the distance between two laser pulses.
  - Sampling points: the number of correlation point to be determined
  - Time window and channels selection:
    - Caution! Symphotime start counting the channels with 1 instead of 0!
    - The green channels are 1 & 3 here and the red channels are 2 & 4.
    - For the time windows, Symphotime determines them “automatically” and tends to remove some of the beginning, either do not move the red lines or take care to always move them to the same values/positions for all your data sets.
- For the autocorrelation of the signal in the green channels in the prompt time window, select “crosscorrelation” and tick the boxes of channel 1 and 3:
-
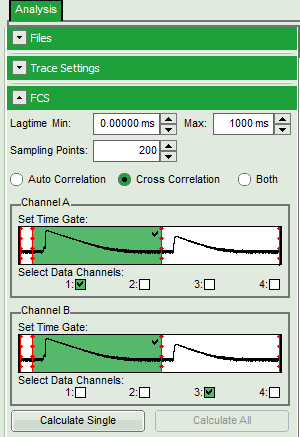

- Press “calculate single” and wait until the result is shown:

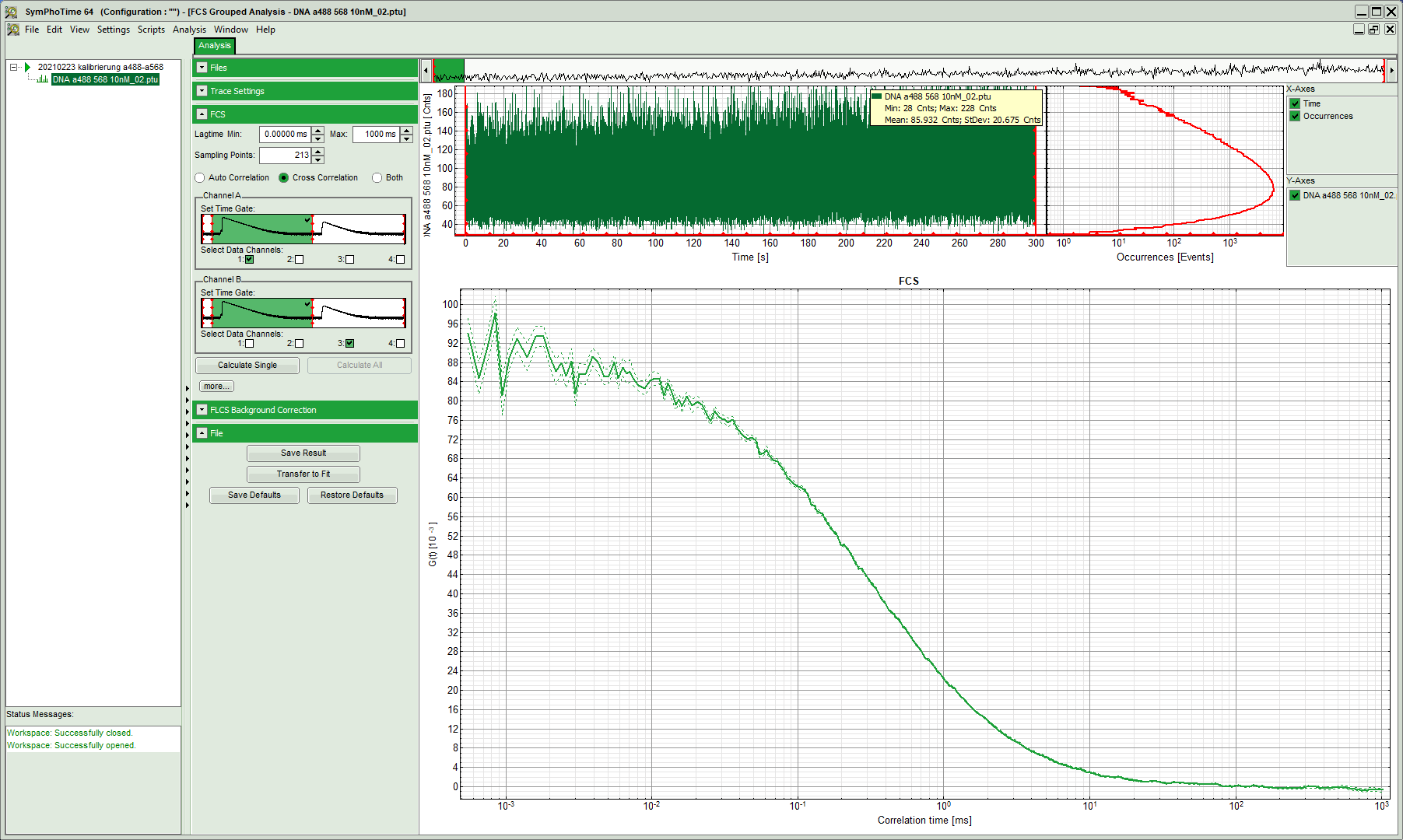

- Caution! Check the labeling on the y-axis, the correlation amplitude. Symphotime tends to report the data scaled by 10^3^. You will need to divide the data manually by 10^-3^ to properly analyse the data in any other data analysis software than Symphotime.
- Export the data by right-click on the curve and select either of the ASCII export option (selected, visible, all cells)

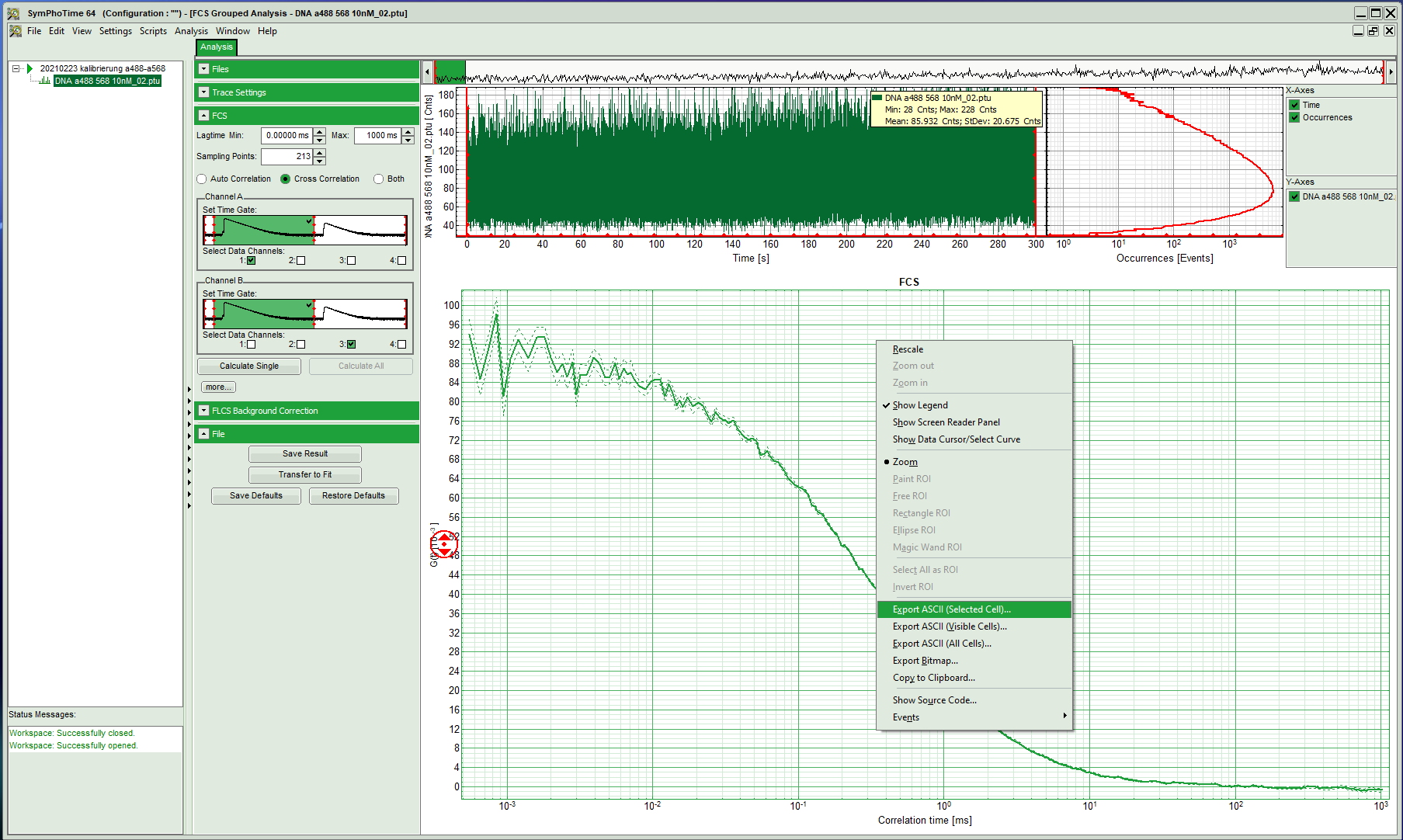

- Proceed to calculate the other two autocorrelation functions:

| Red signal in the prompt time window | red signal in the delay time window |
| --- | --- |
| 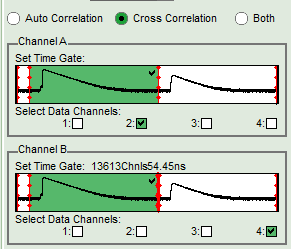 | 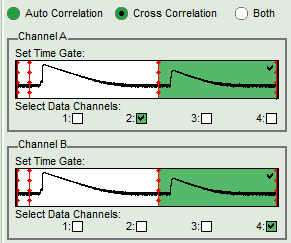 |

- Next, determine the two required crosscorrelations:

| Green signal in prompt time window with red signal in prompt time window | Green signal in prompt time window with red signal in delay time window |
| --- | --- |
| 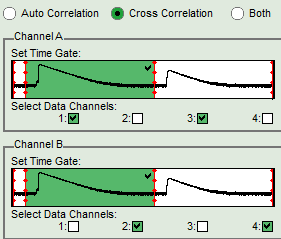 | 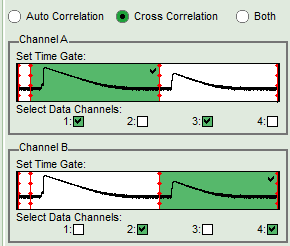 |

- Export all generated curves
  - For the data export, consider which software you would like to use for the fitting. Some software (Kristine, ChiSurf) might require a single
- Finally, determine the count rates for the different channels and time windows
  - The “Header” file of your data set displays the average count rate for each channel, however, not separation between “prompt” and delay time window is made.
- Select “Intensity Time Trace” and press start.

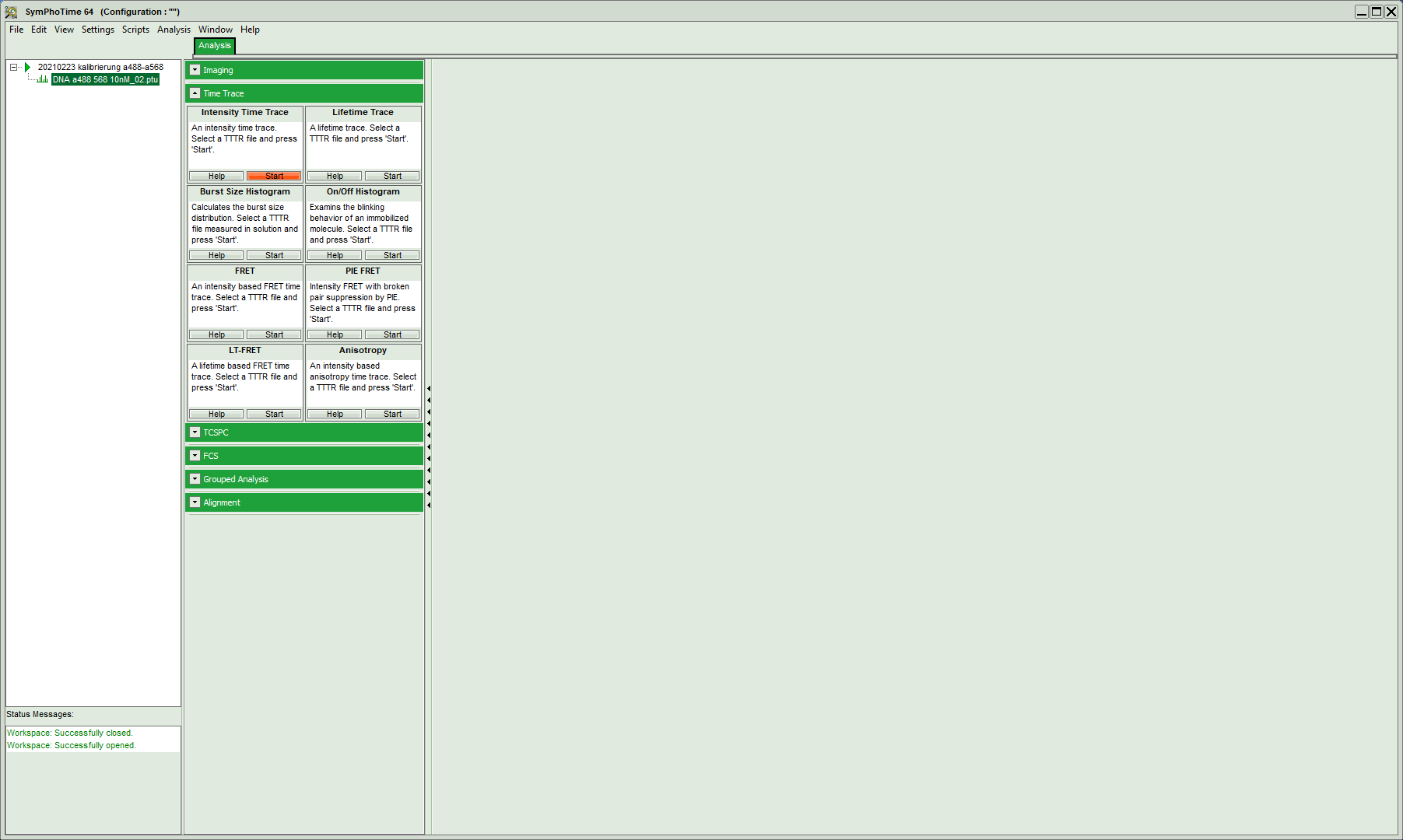

- Select the “prompt” time window and leave all for channels selected and press calculate:

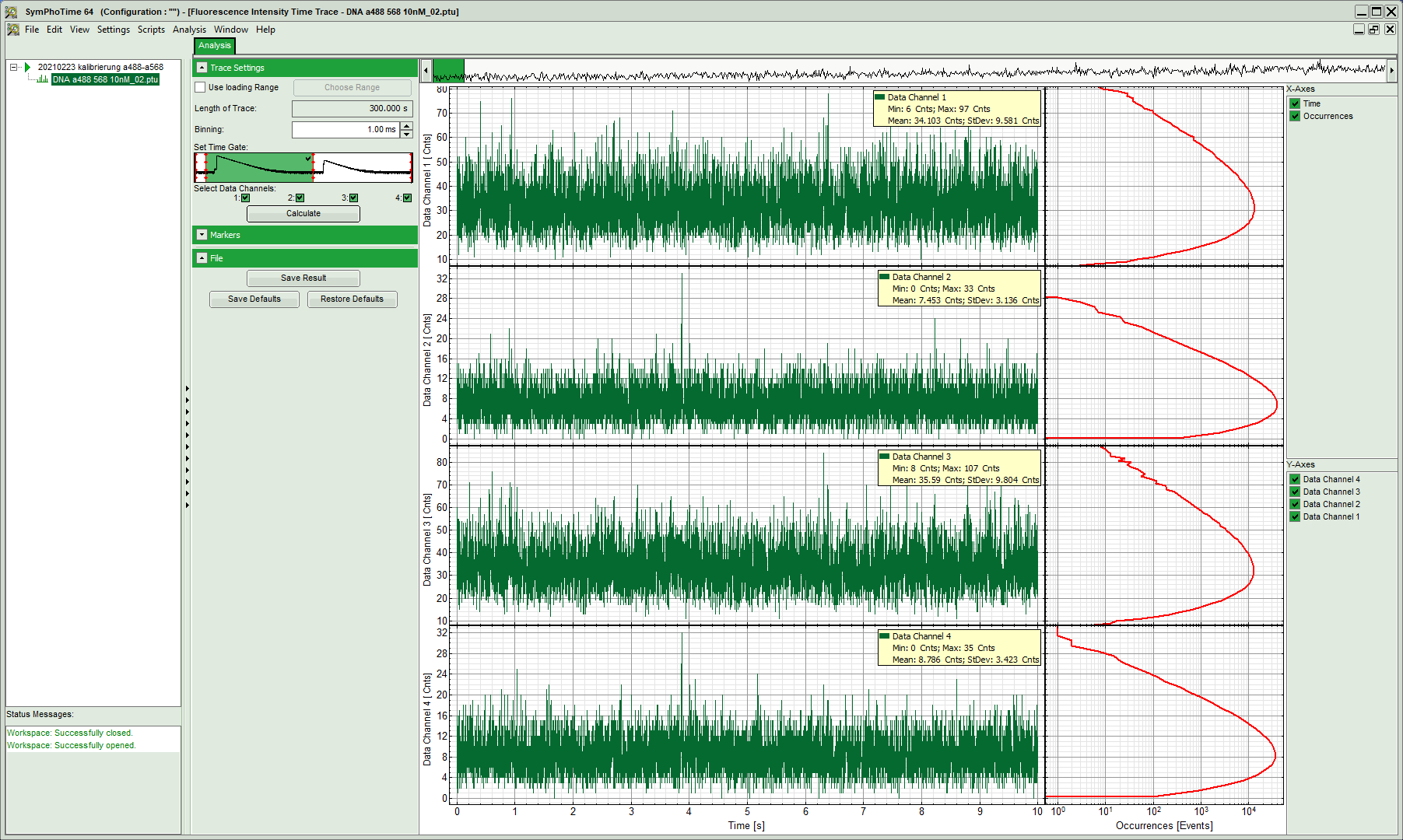

- Note down the mean values (and standard deviations, if required) in your protocol.
- Next, switch to the “delay” time window and repeat the calculation:

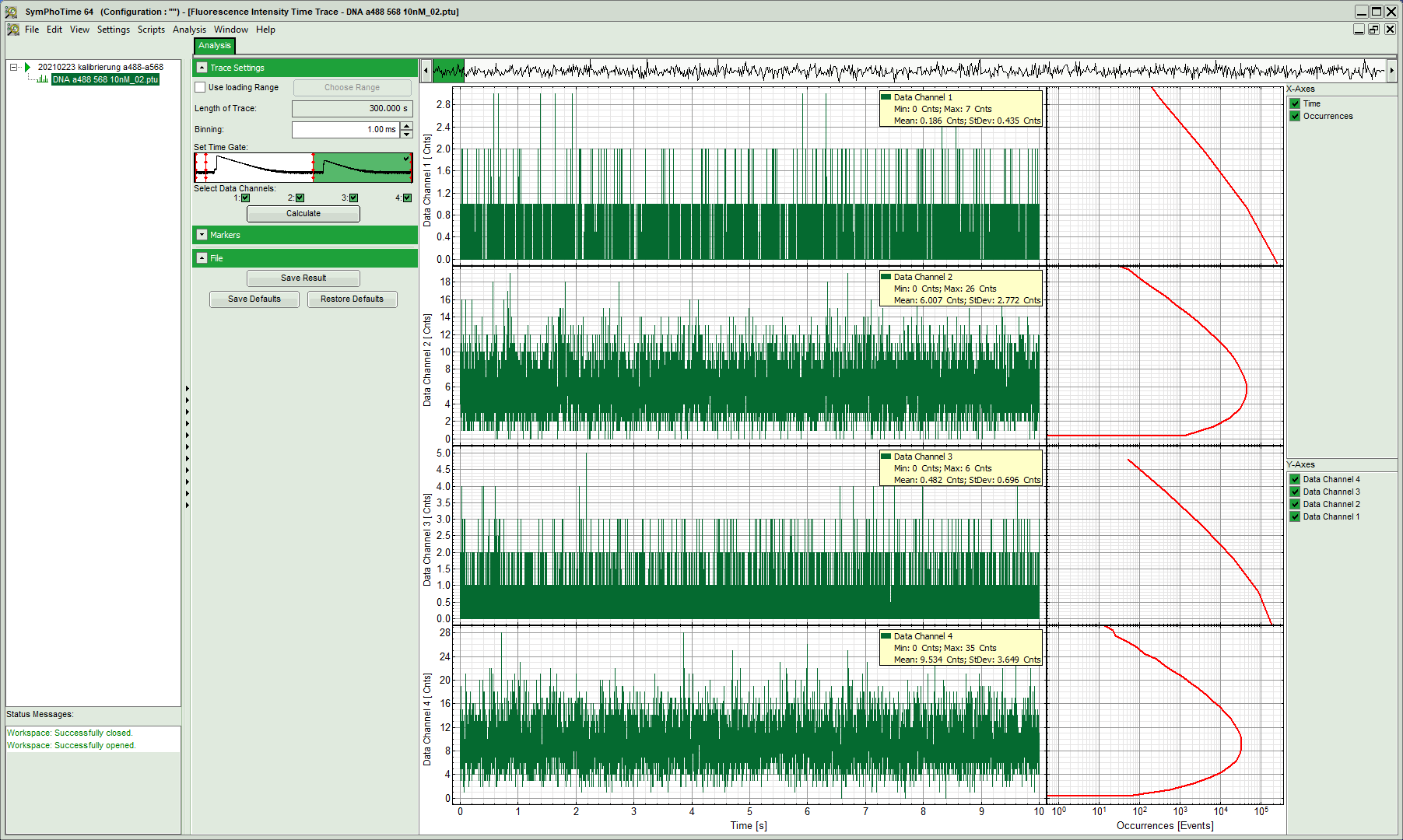

###### Advanced analysis: correcting for background

In cell measurements quite high background count rates might be reached compared to the relatively dim fluorescence of the fluorescent proteins. The Symphotime software offers an option to remove this background from your correlation.

- Switch back to the “FCS” window and unfold the “FLCS background correction” option:

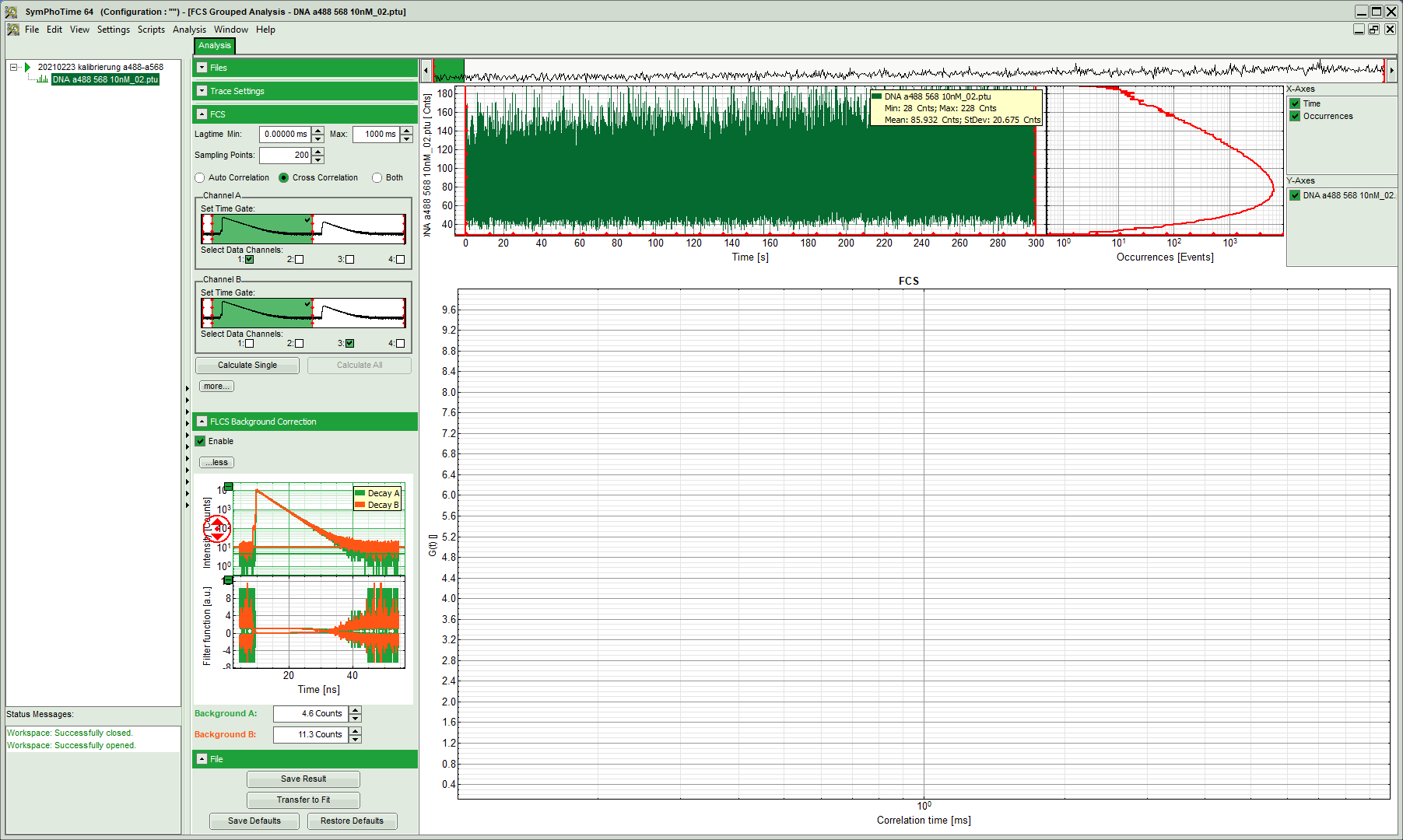

- In the bottom right, now two plots appear:
  - Top one shows the fluorescence intensity histogram on the nanosecond time scale
  - Bottom one shows the weighting factor calculated based on these fluorescence intensity histograms. These weighting factors will be applied in the next step during the correlation.
- Adjust the background or offset of the fluorescence intensity histograms such that the horizontal green and orange lines run through the offset of the histogram:

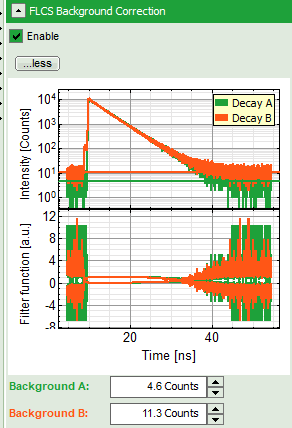

- Run the calculation by pressing “calculate single” and compare the results. For the shown DNA measurement, no big difference is visible. However, in cells with high background this analyse might improve the result.

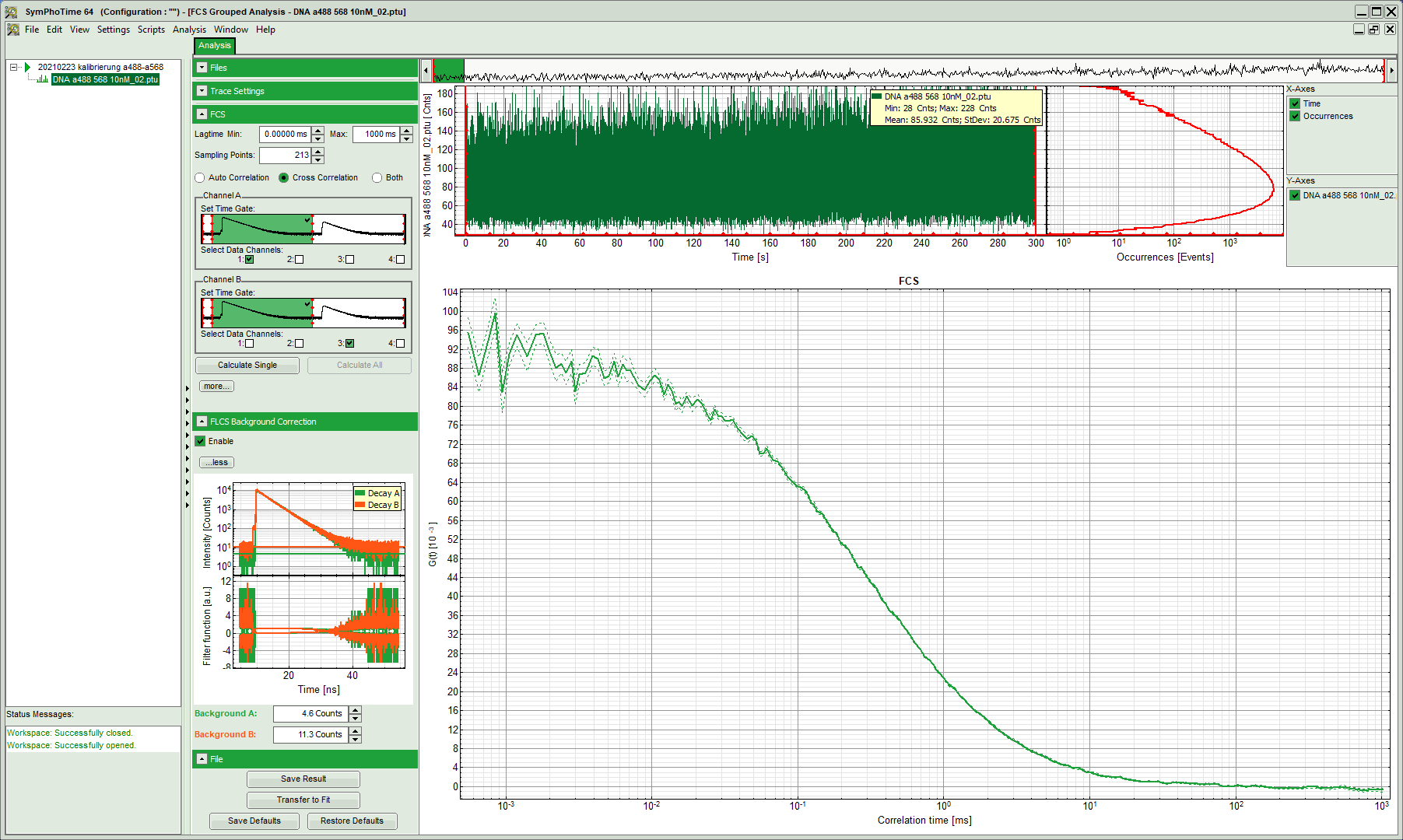

##### Kristine

- Open Kristine_GLOBAL LV2012 x64_sf and select your data format via “Options” -> “Select Setup”

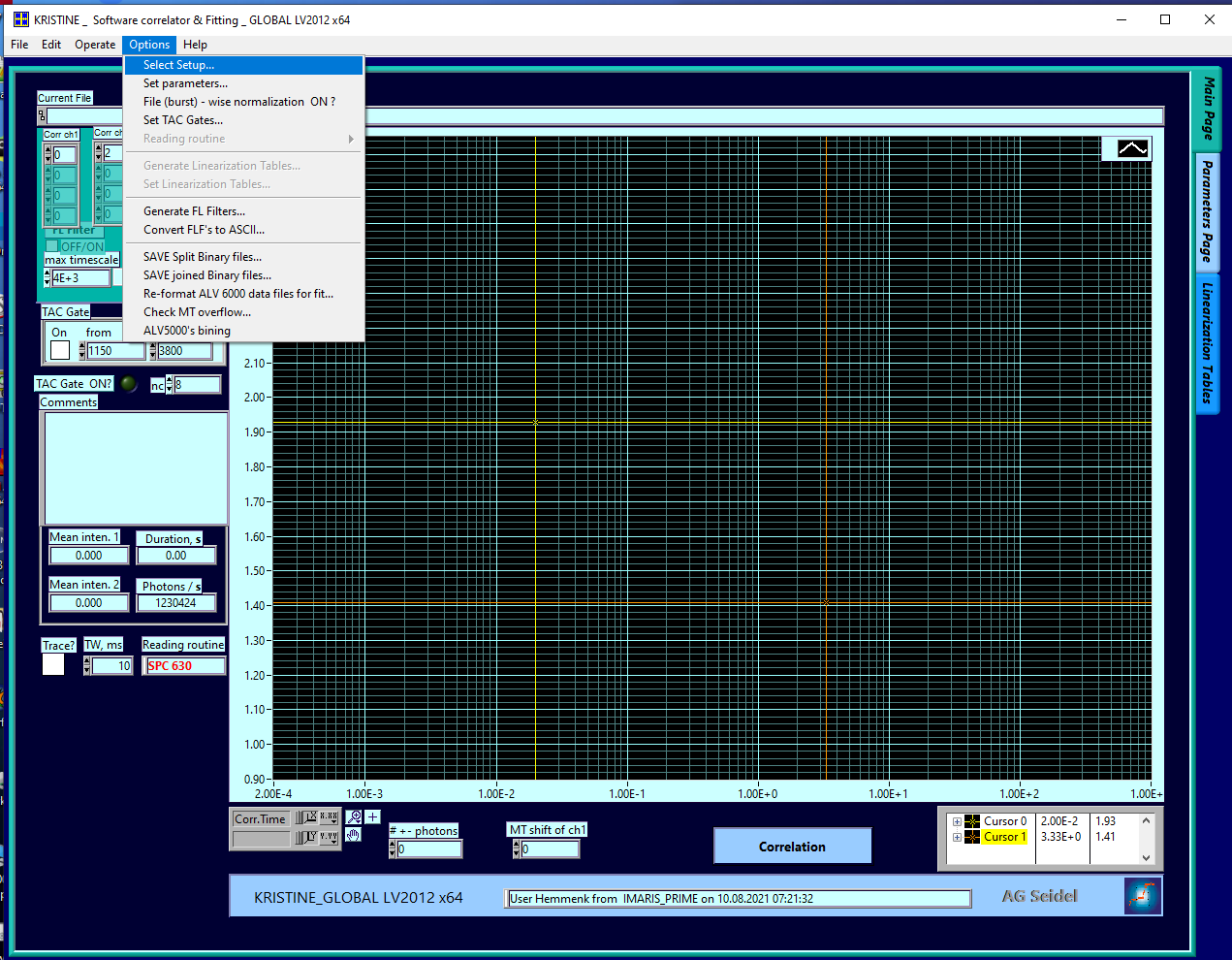

- In the newly opened window, select your data format from the drop-down list at the bottom

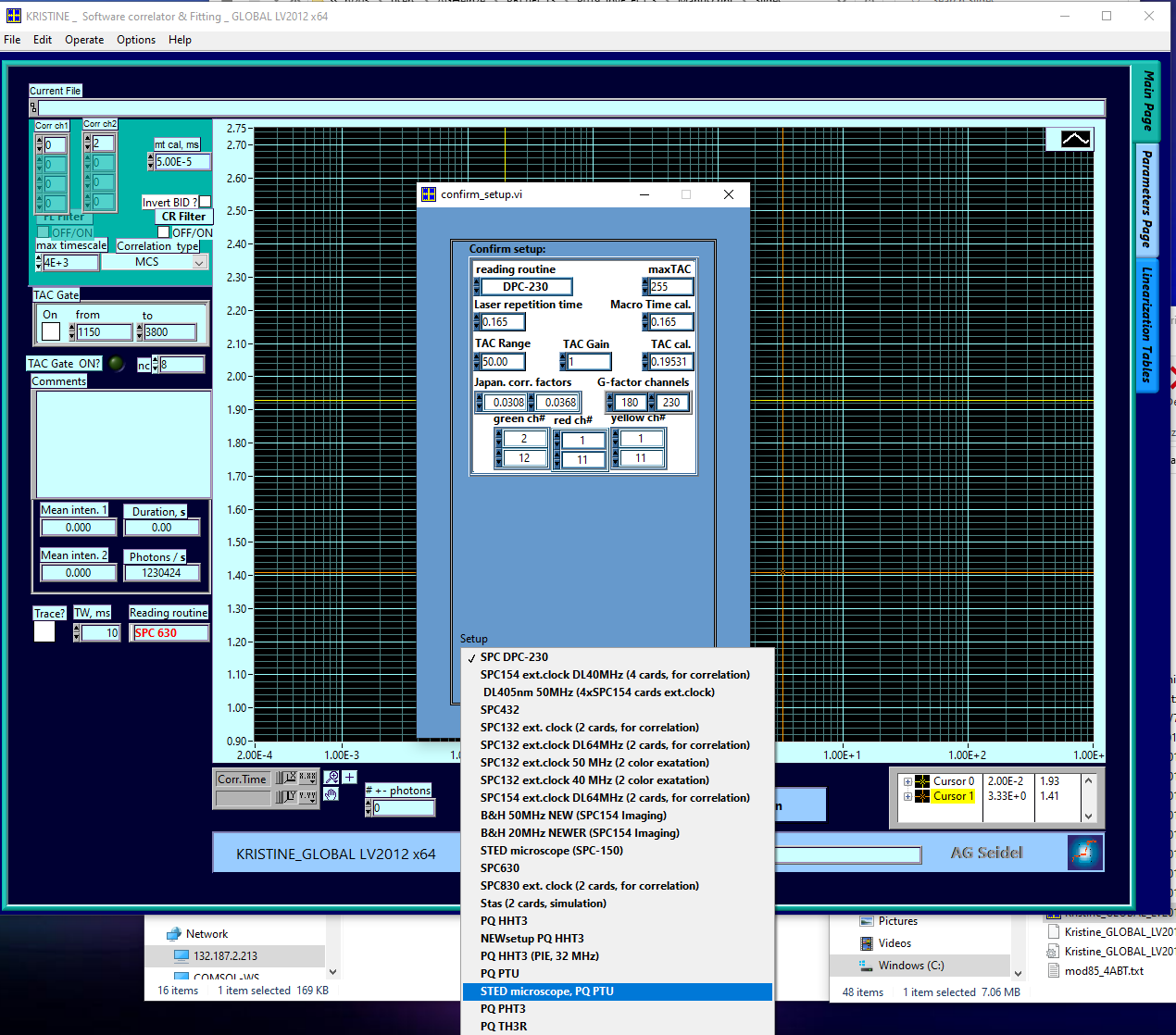

- Here, we select the “STED microscope, PQ PTU”. The name is not important, what matters is the reading routine: “PQ PTU” stands for Picoquant ptu- format.

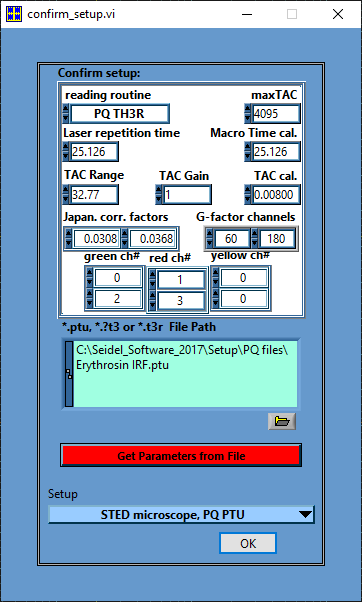

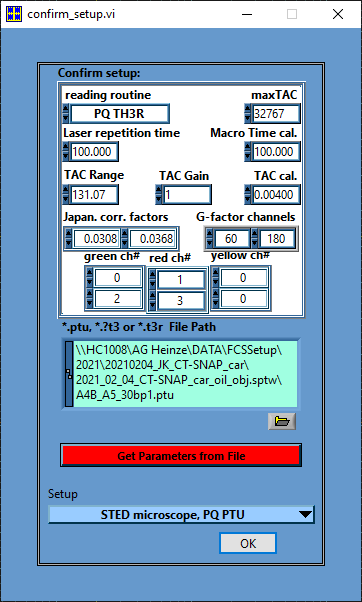

- Next, load an example file from your measurement day. This file s used to collect the header information about the measurement settings. Which channels were used? What was the repetition rate? What was the time resolution?
- Press “Get Parameters from file” and observe how the values change.
- Adjust the “Correction factors” and “G-factor channels”, if your plan to use the anisotropy information, usually not required in correlation analysis
- Adjust the channels numbers: Top row are the perpendicular channel numbers in green, red (and yellow) and bottom row are the parallel channels of the green, red (and yellow) detectors.
- Here, the numbers are correct.
- Press “OK” to save the settings and to return to the main window.
- Set your time gates for PIE correlation:

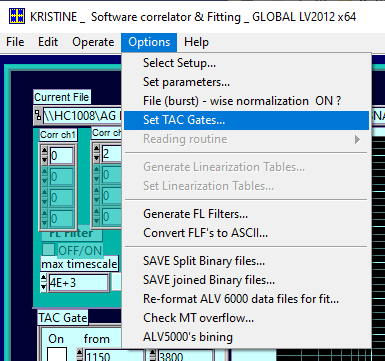

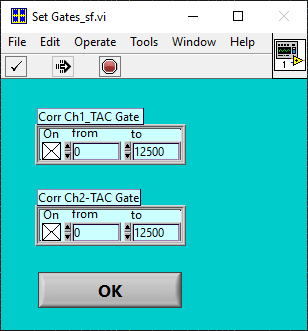

- In the shown example, the time window (TAC window) between two laser pulses of the same color is 100 ns, the base resolution is 4 ps, thus the data is distributed into in total 100 ns / 4 ps = 25’000 time channels. The first 12’500 channel represent the “prompt” time window, the second half the “delay” time window.
- Set the TAC gates for both correlation channels to 0 -12500, select the tick box and press ok.
- In the main window, set your correlation channels – we leave them at 0 and 2 for now, this will give us the autocorrelation of the green channels in the prompt time window.
- Additionally, you could increase the maximum correlation time, now 4000 ms, e.g. for slow diffusion molecules.
- Note: The bright green spot next to “TAC gate ON?” indicates that you will be using TAC gating during your correlation.

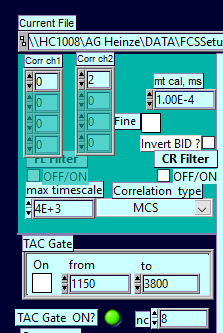

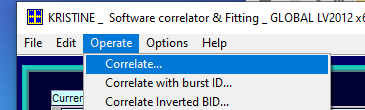

- Run the correlation via “Operate -> Correlate”
- Select your data to correlate in the file selection window:

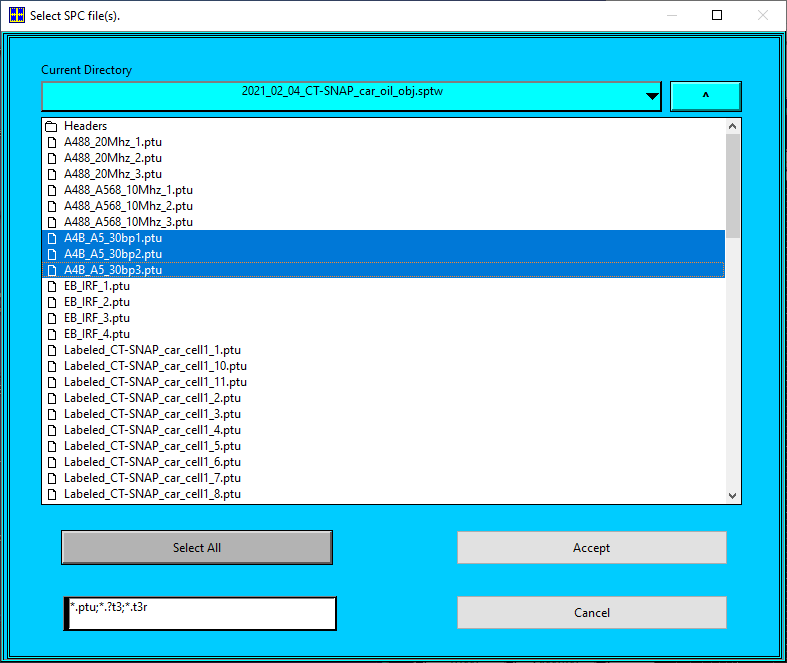

- You can select one or multiple files
  - Caution: multiple files will be correlated subsequently and averaged, thus they must belong to the sample.
  - For cell samples, it is recommended to first correlate each data set separately, then decide which correlations are “good ones”, repeat the process with the selected measurements and obtain the average values.
- The average count rate per channel, the total measurement duration and read photons per second are summarized on the left-mid:

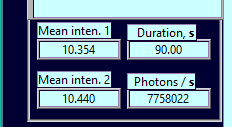

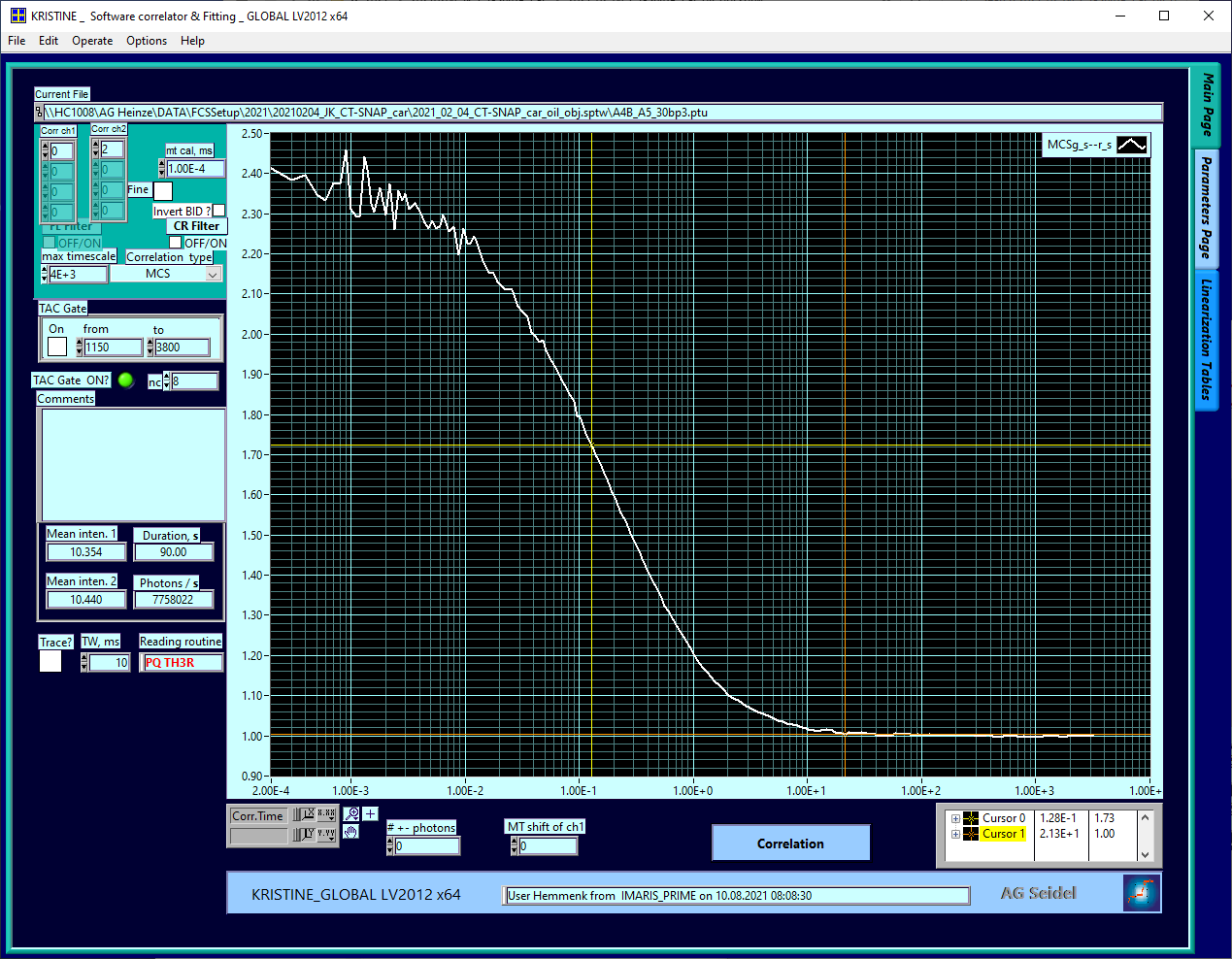

- Save the data by pressing “control + S” or via “File” -> “Save plots for fit”
  - Note: Kristine supports batch saving, thus it is most convenient to perform all correlations of sample first and finally save the data.
- In total, up to five correlations are required, the top row show the TAC gating setting, the bottom row the selected correlation channels.

| ACF “green-prompt” | ACF “red-prompt” | ACF “red-delay” | CCF “FRET” | CCF “PIE” |
| --- | --- | --- | --- | --- |
| 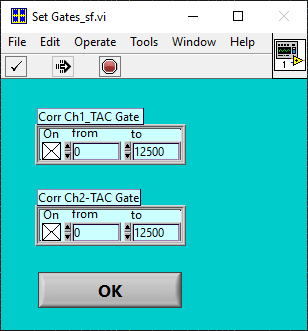 | 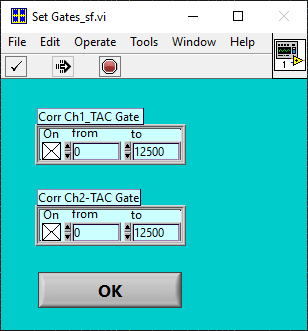 | 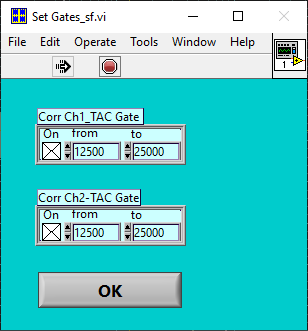 | 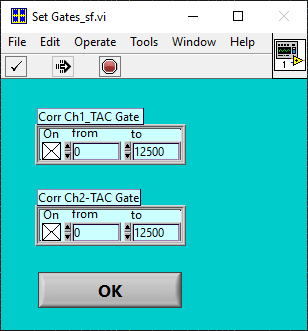 | 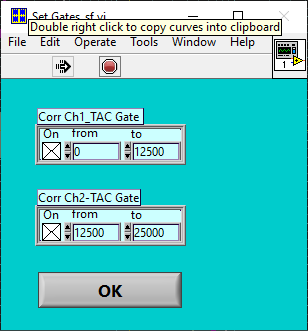 |
| 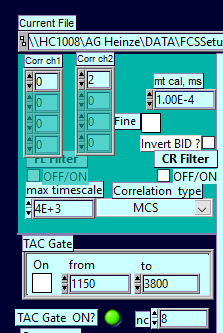 | 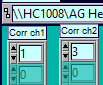 | 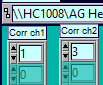 | 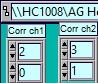 | 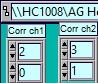 |

- All five required correlations of a FRET sample:

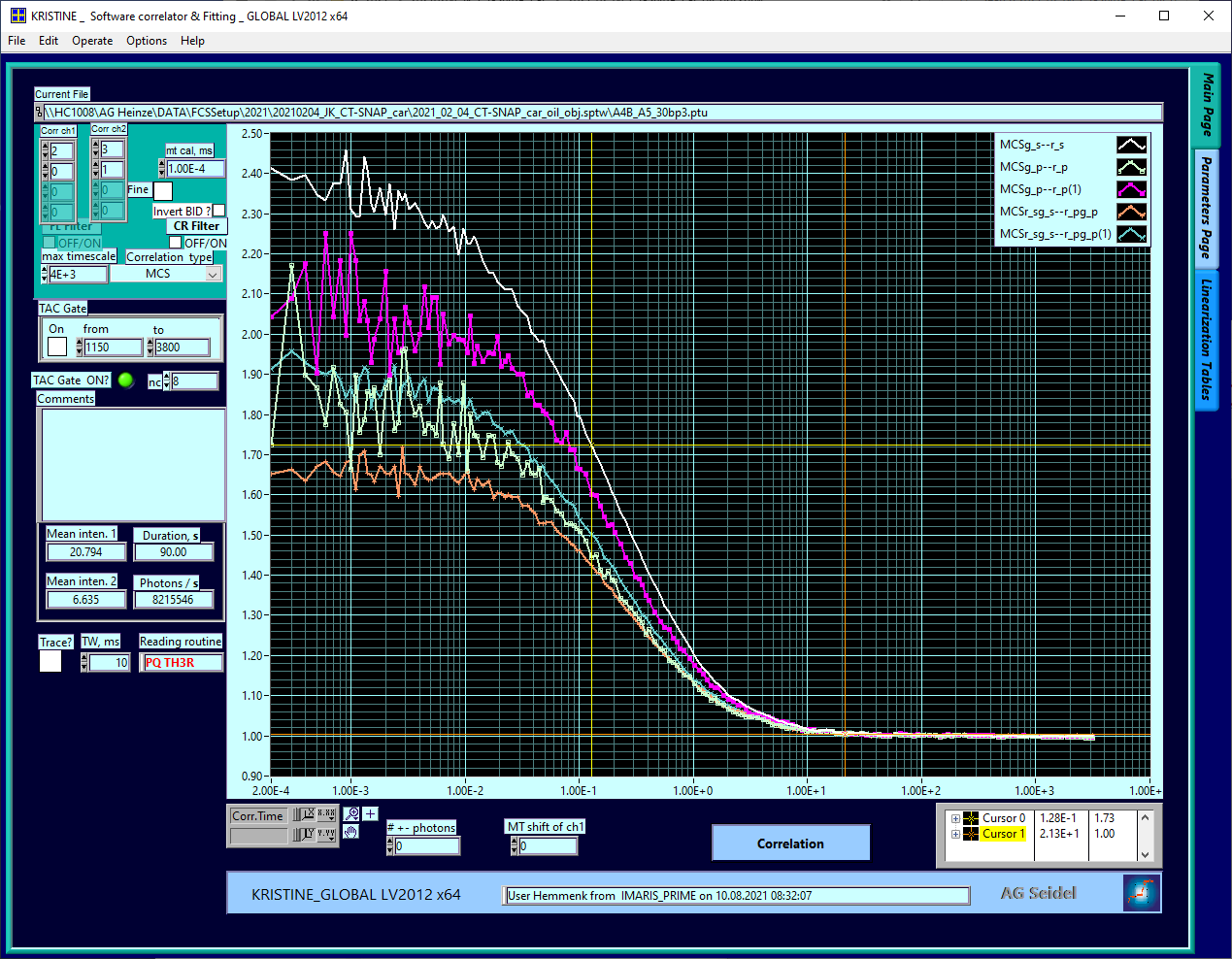

- Save all curves and rename them if required.
- The exported .cor-files and .inf-files can be opened with common text editors, e.g. Notepad or Notepad++.
- The files contain four columns: (1) Time in milliseconds, (2) correlation value, (3) measurement time and average count rate and (4) standard deviation, if available, else this column is filled with zeros.
  - Note: In the absence of a standard deviation, the time and average count rate is used to estimate the weighting factors during the fitting

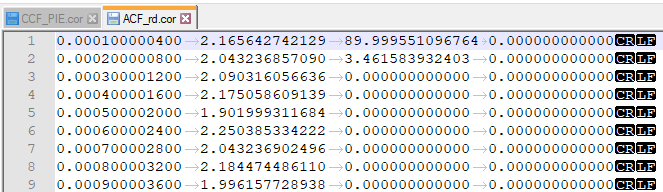

##### Python-scripts

###### Install Miniconda (Python3.7, usually 64-bit)

If you haven’t done so, please install Miniconda first.

<https://docs.conda.io/en/latest/miniconda.html>

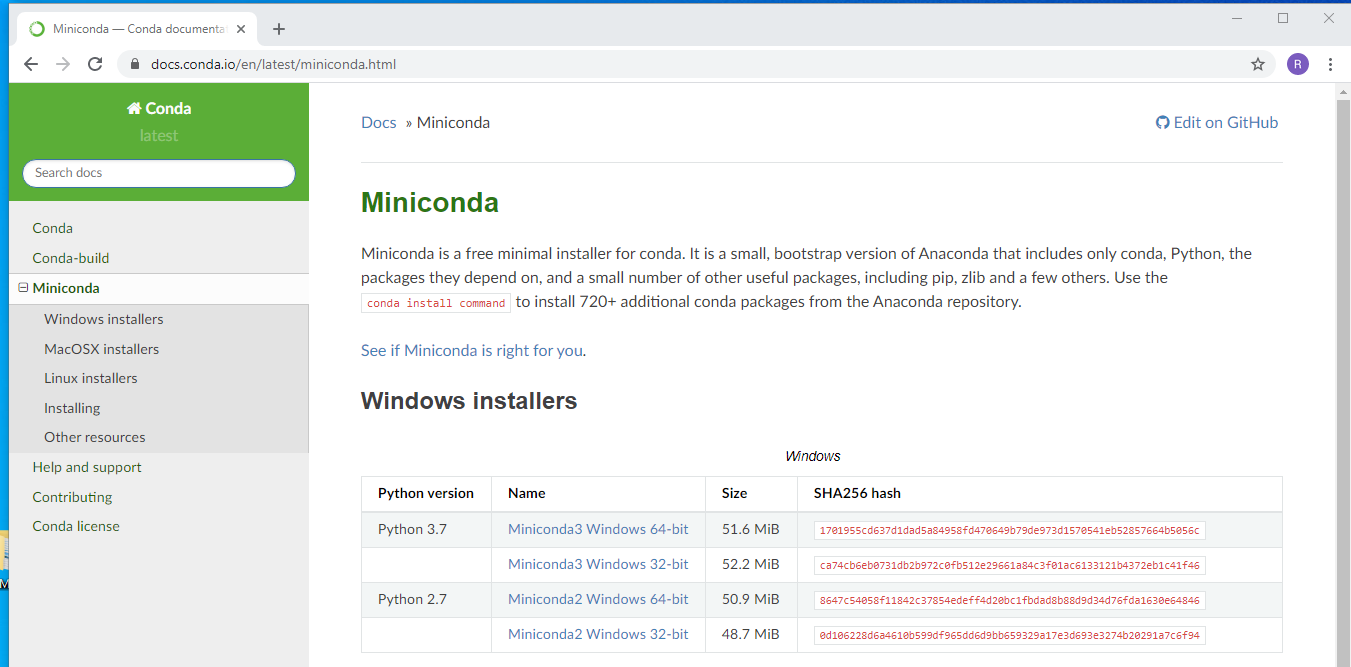

Install for all users:

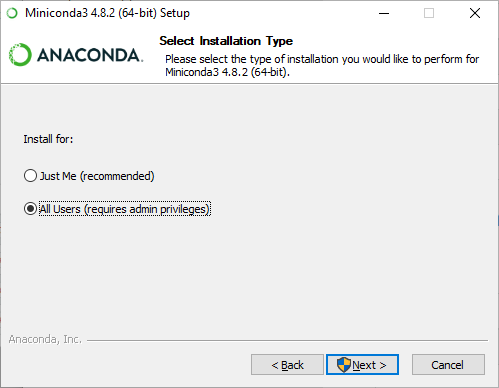

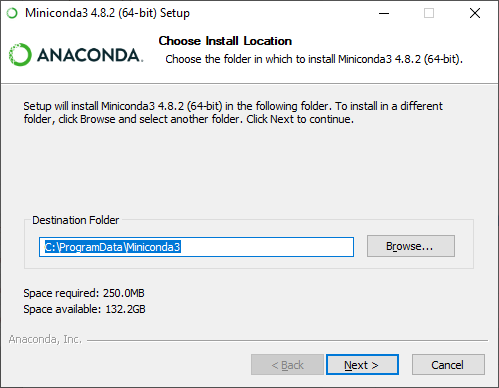

###### Getting “scripts” from Github

Go to the following website and download the newest version:

<https://github.com/HeinzeLab/JOVE-FCS>

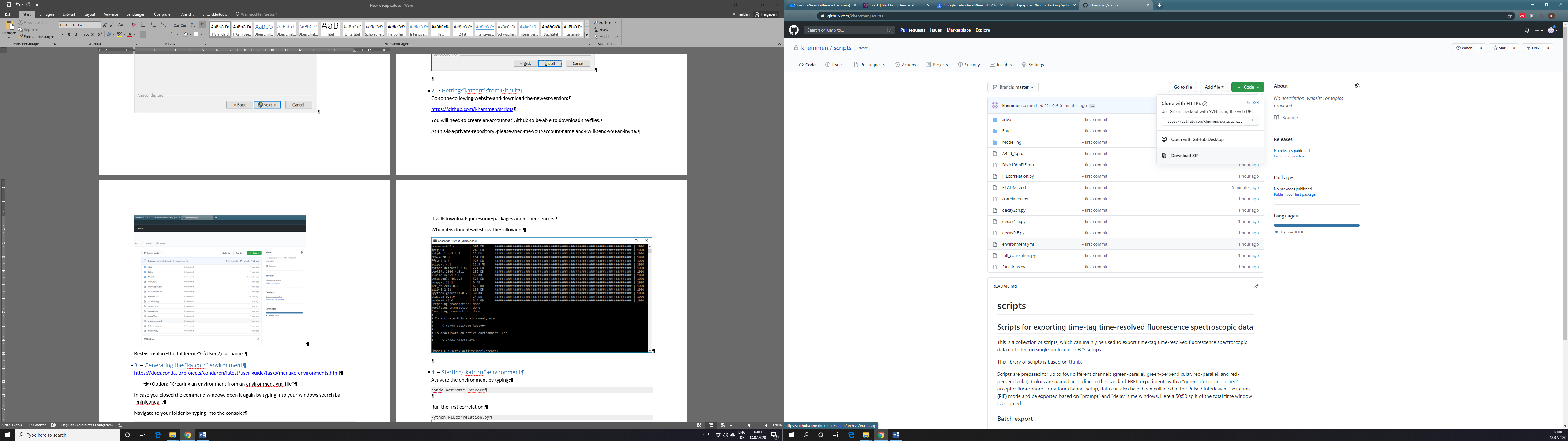

Best is to place the folder on “C:\Users\username”

###### Generating the “scripts” environment

<https://docs.conda.io/projects/conda/en/latest/user-guide/tasks/manage-environments.html>

Option: “Creating an environment from an environment.yml file”

In case you closed the command window, open it again by typing into your windows search bar “miniconda”.

Navigate to your folder by typing into the console:

cd C:\Users\username\scripts

Generate the new environment named „scripts”:

conda env create -f environment.yml

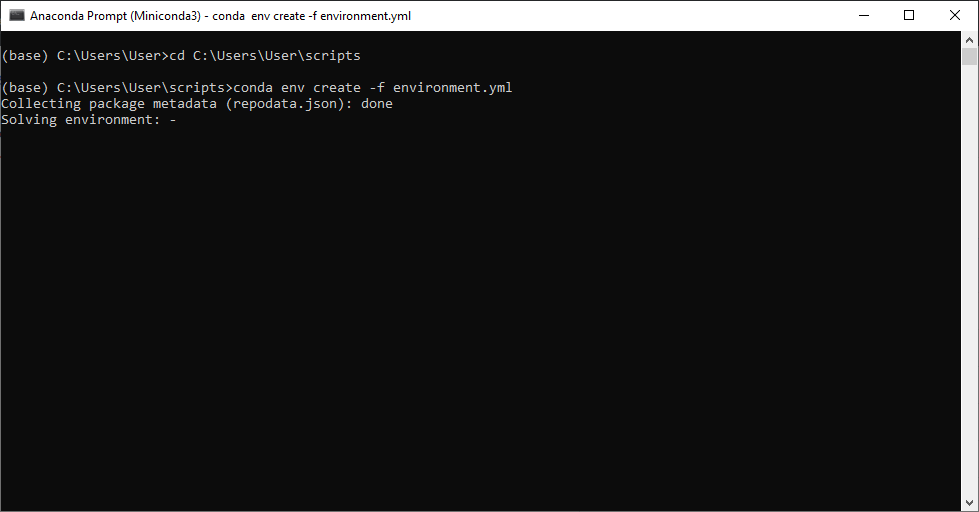

It will download quite some packages and dependencies.

When it is done, it will show the following:

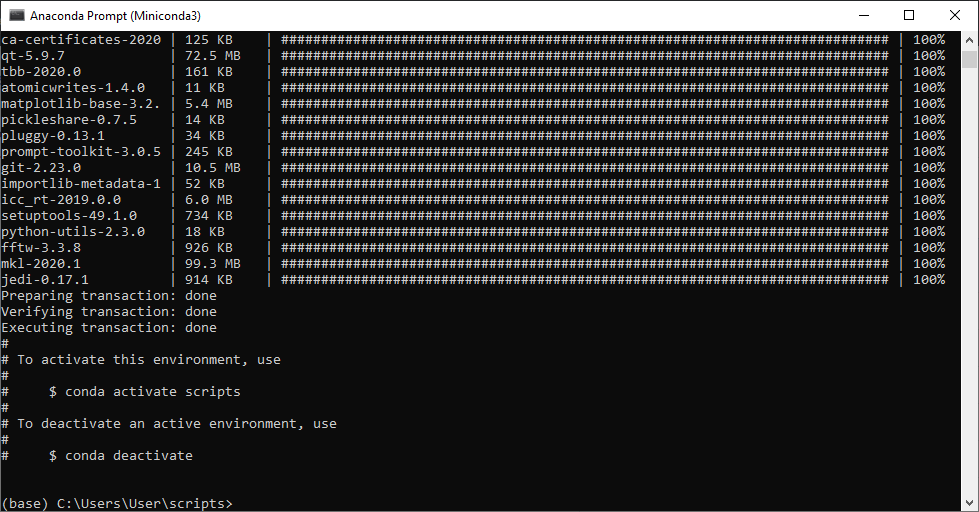

###### Starting “scripts” environment

Activate the environment by typing:

conda activate scripts

When you want to use “scripts” next time, you can directly start from section 4 using the environment activation.

Please make sure you navigate to the scripts-folder before trying to run any command.

(base) C:\Users\User > conda activate scripts

(scripts) C:\Users\User > cd C:\Users\username\scripts

(scripts) C:\Users\username\scripts >

###### Running your first command

e.g. correlate a single data file, which had been obtained by a PIE measurement:

🡪 2 channels from two colours each (“donor”, green, and “acceptor”, red), the two channels of a colour are arranged in two different polarizations “s” and “p”, horizontally and vertically to the excitation laser beam.

The example data is called: DNA10bpPIE.ptu

Start the correlation by typing “python PIEcorelation.py”

(scripts) C:\Users\user\scripts > python PIEcorrelation.py

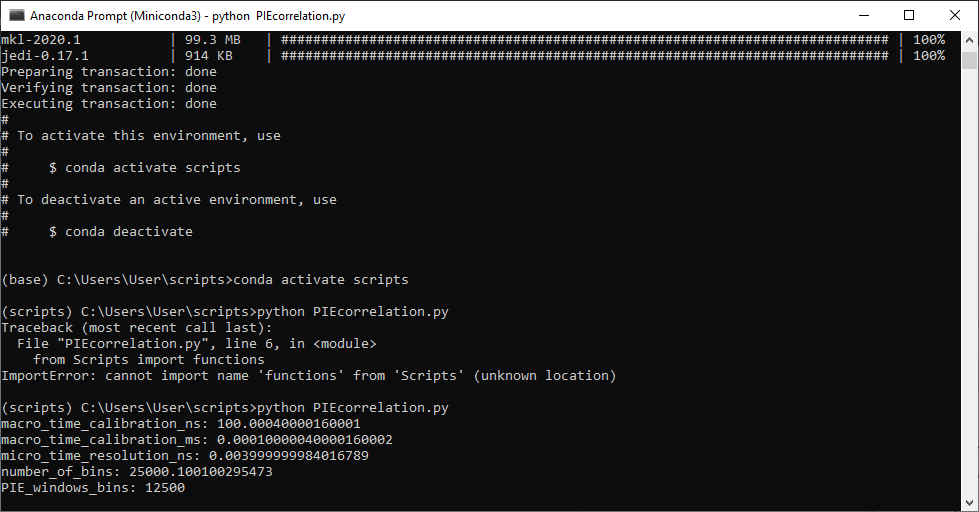

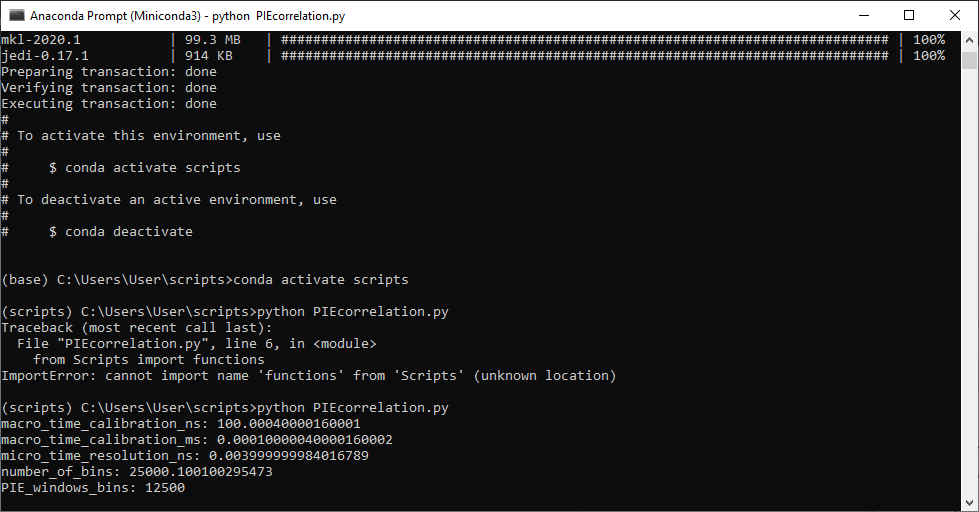

You might want to check the script in Notepad++ (or any editor) to check it and to modify your filename (line 12):

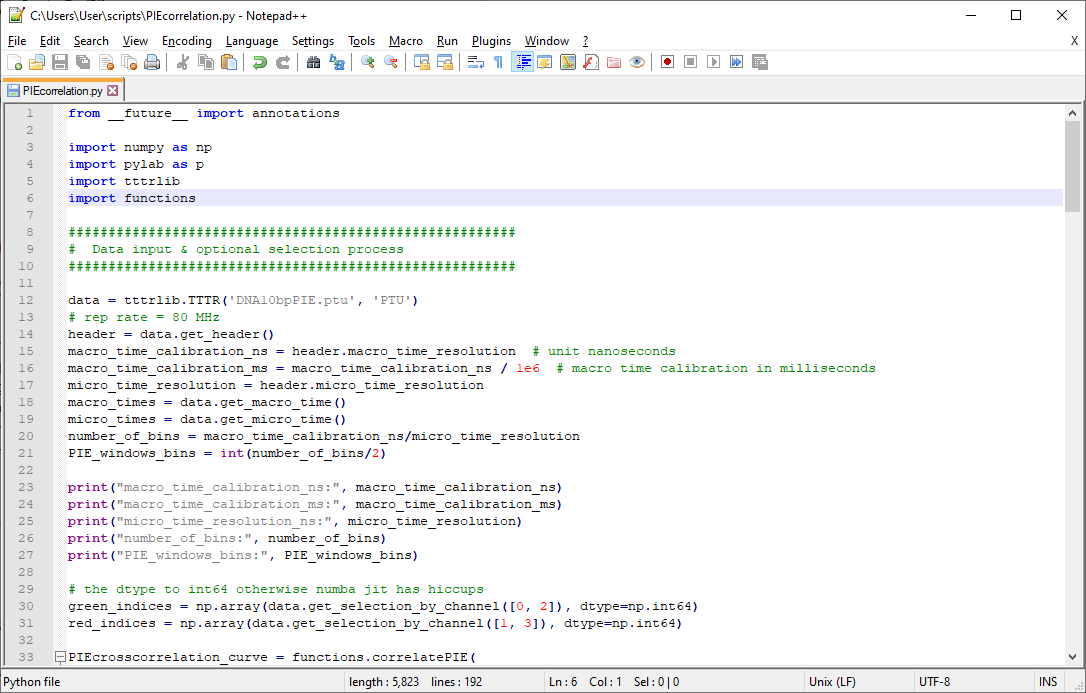

After the correlation is done, a new window containing the data plot is opened:

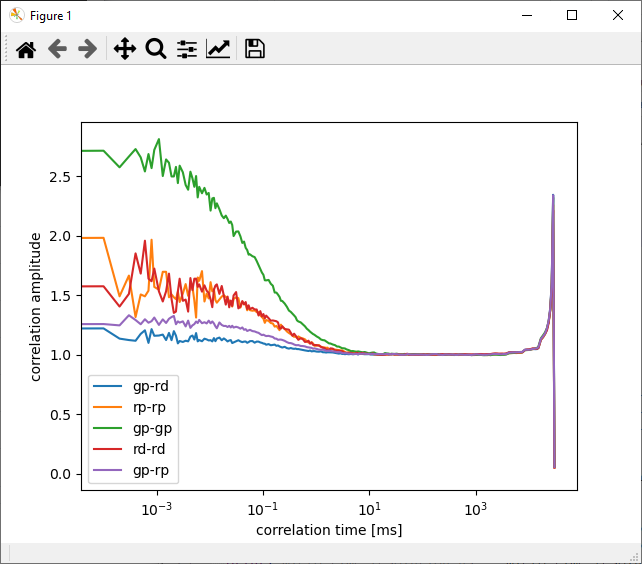

**CLOSE this window to be able to proceed!**

Additionally, the data is saved to your input folder:

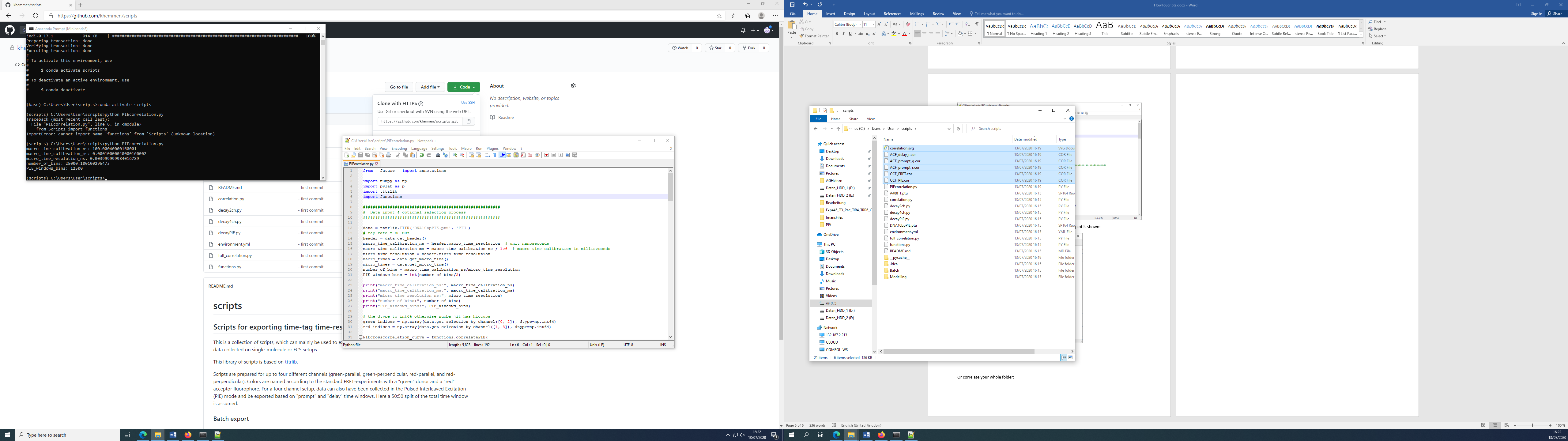

To process a different data set change the filename in line 12 of the scripts.

As it is unpractical to copy all your data to your “C:\Users\User\scripts” folder, you can replace the filename by a data path:

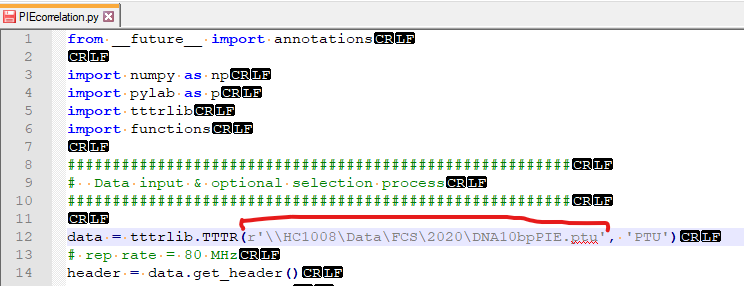

###### Batch correlation of whole folders

If you have many data files, which you all would to process the same way, you can use the scripts in the “batch” folder. These are made for batch processing.

Navigate to the “batch folder” in the console:

(scripts) C:\Users\user\scripts > cd C:\Users\user\scripts\batch

(scripts) C:\Users\user\scripts\batch

This scripts consist of two parts:

- your python script, which does the job and
- your “settings” file, which is a so-called YAML-file. This can also be edited in Notepad.

For correlating a folder of PIE data sets open the “PIEcorrelation_settings.yaml” file in Notepad and adjust your settings as required:

i.e.

- make sure the channel definitions are correct

- adjust the naming suffix for autosaving

Don’t forget to press save after made your changes!

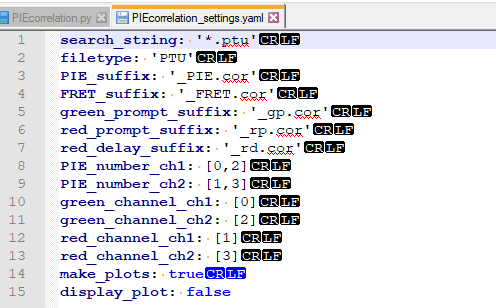

Now switch back to the console and type:

(scripts) C:\Users\user\scripts\batch > python batch_correlation.py --settings PIEcorrelation_settings.yaml --path “\\HC1008\Data\FCS\2020”

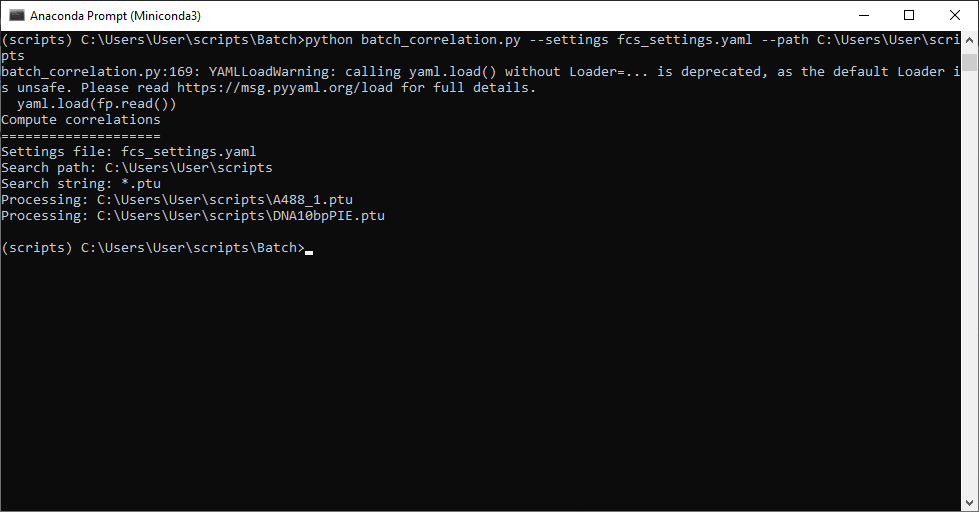

This scripts needs two input parameter:

--settings: your modified settings file

--path: the path to your data, please enclose the path in “ “

**CAUTION!!!**

- **No dots (.sptw!),**
- **no special characters (µm!),**
- **to change drive add path in “E:\path-to-data”**

Here, all folders within your specified path will be search recursive for the key phrase “.ptu” and all ptu-files will be correlated (ch0 and ch2), the auto- and crosscorrelation and a figure for visual inspection is saved:

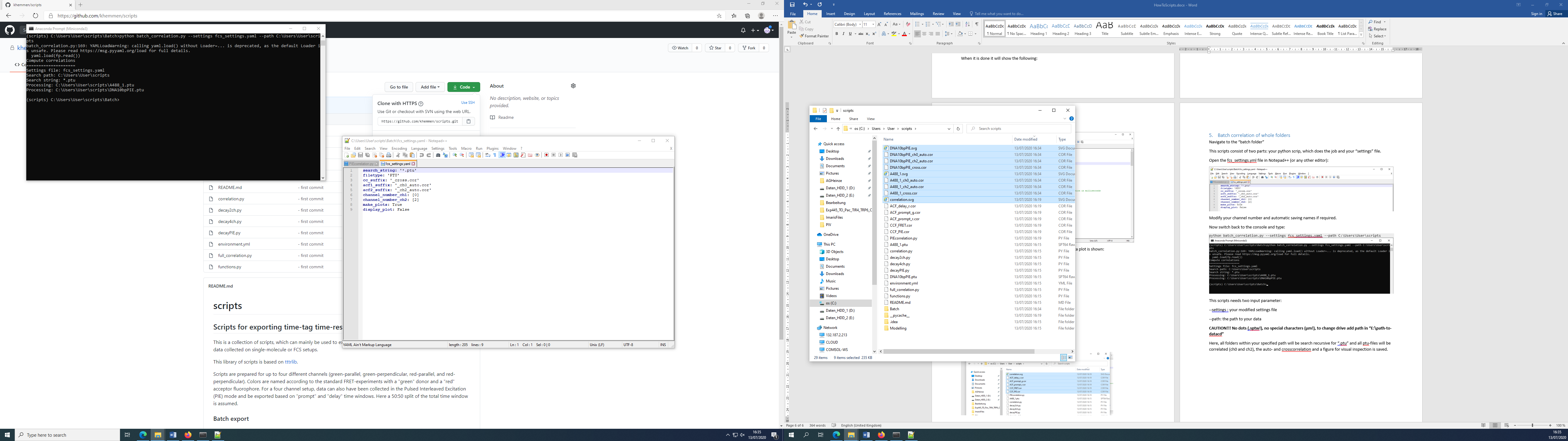

In case you want to correlate e.g. only your cell data, you can change the search phrase to ‘cell*.ptu’, if you data is saved as cell1.ptu, cell2.ptu etc. Calibration measurements named e.g. A488.ptu will not be correlated.
