## Supplementary File 1-7 for "Dual-color Fluorescence Cross-Correlation Spectroscopy to study Protein-Protein Interaction and Protein Dynamics in Live Cells": SuppNote4_FCCS analysis using ChiSurf.docx

### Analysis of FCCS experiments using ChiSurf

#### Analysis of the Fluorescence Correlation curves

In this guide, we will first work solely with the Fluorescence Correlation (FCS) curves exported from your calibration data and use the different measurement to characterize the daily performance of your system. It is assumed that you have exported your data already using (i) the proprietary software of your setup (e.g. Symphotime from Picoquant), or (ii) other, maybe self-written software. In case you are still searching for a software, which fits your needs best, you might try

- Kristine (<https://www.mpc.hhu.de/software/3-software-package-for-mfd-fcs-and-mfis>)
- PAM (<https://gitlab.com/PAM-PIE/PAM>)
- Globals for FCS (<https://www.lfd.uci.edu/globals/>)
- Our self-written scripts for batch export (<https://github.com/HeinzeLab/JOVE-FCS>)

Once you have your workflow and templates established the calibration data is easily analyzed.

In the second section of this guide, we will show you how to (i) add your own fit models, and (ii) analyze live cell experiments.

All data analysis are performed using ChiSurf V2016 ^1^, which can be downloaded here:

<https://github.com/Fluorescence-Tools/chisurf/releases/tag/16.05.14>

Of course, all analysis can also be performed with other software, which supports the implementation of the described fit models, using weights in the fit routine and global analysis.

##### Data format

For analysis with ChiSurf, your correlation data has to be exported in text format with either three or four columns:

- Column 1: correlation time
- Column 2: correlation amplitude
- Column 3: first value reflects the measurement time, second value the average count rate, the rest of this column is filled with zeros OR: the whole column is filled with zeros.
- Column 4: standard deviation of the correlation amplitude

The measurement time and the average count rate in column 3 are used to estimate the uncertainties in your correlation amplitudes if the standard deviation is not available, i.e. single measurement was performed or the hardware / software used for correlation does not provide these values.

#### Analysis of Calibration measurements

##### Provided test data - calibration

| Sample | Purpose | Measured |
| --- | --- | --- |
| ddH2O / buffer | background / dark counts | 2x |
| IRF (EB quenched with KI) | Instrument response function | 2x |
| A488 | green calibration dye | 5x |
| A568 | red calibration dye | 5x |
| DA-labeled DNA | green-red overlap calibration | 5x |

**Settings:**

- 2-color excitation (PIE)
- Excitation power: 485 nm: 6 µW, 568 nm: 1.4 µW
  - measured at objective
- 10 MHz rep rate for each laser line (485 nm prompt, 568 nm delay)
- TAC window: 100 ns (50 ns delay between pulses)
  - TCSPC bin size: 4 ps
- Calibration samples were measured as drop on objective slides

##### Provided test data – live cell experiments

| Sample | Purpose | Description |
| --- | --- | --- |
| eGFP | Donor-only sample (DOnly) | eGFP inserted into the intracellular loop 3 of the β_2_ adrenergic receptor (β_2_AR) |
| SNAP | Acceptor-only sample (AOnly) | SNAP-tag attached to the C-terminus of the β_2_AR, labeled with SNAP Surface DY-549 |
| NT-SNAP | Double-labeled sample (DA) which does not show FRET | β_2_AR construct with both eGFP inserted into the intracellular loop 3 and SNAP-tag attached to the N-terminus, labeled with SNAP Surface DY-549 |

**Settings:**

- Experimental settings are identical to the calibration measurements.

##### Provided test data – simulation

| Sample | Purpose | Description |
| --- | --- | --- |
| CT-SNAP | Double-labeled sample (DA) which shows FRET | β_2_AR construct with both eGFP inserted into the intracellular loop 3 and SNAP-tag attached to the C-terminus, labeled with SNAP Cell TMR |

**Settings / Assumptions:**

- Bimodal diffusion of molecules with 30 % of a fast diffusing species (*t_D1_* = 1 ms) and the rest of the time diffusing slowly with *t_D2_* ~ 100 µs
- Dynamic exchange between two equally populated FRET states, high FRET (HF; FRET efficiency *E* = 0.7) and low FRET (LF, with *E* = 0.2), with a time constant of *t_R_* = 70 µs
- Additionally, 16 % of triplet blinking at 5.5 µs was added.

##### Determination of background / dark count rate

Firstly, the average count rate from ddH2O or appropriate buffer (untransfected cells in case of cells) is obtained – under the same excitation condition as for the sample measurements!

| Channel number | Channel name | Count rate [kHz] (prompt) | Count rate [kHz] (delay) |
| --- | --- | --- | --- |
| 0 | g-p (green parallel) | 0.13 |  |
| 1 | r-p (red-parallel) | 0.445 | 0.375 |
| 2 | g-s (green perpendicular) | 0.23 |  |
| 3 | r-p (red perpendicular) | 0.79 | 0.39 |

The average countrate can be determined from your data in different ways:

- If you measured with the Picoquant software, then the average countrate is written in the header section and can be easily obtained by opening the dataset in Symphotime.
- Else you can get the number of photons detected during measurement time from an export of a photon arrival time histogram and divide this number by your measurement time
- Additionally, you can use the “determine_countrates.py” script to export this information for your whole folder of image (see Help file for using scripts).

##### Calibration of detection volume – green excitation

In a first step, we will use the data from the individual diffusing red and green fluorophores and determine (i) the shape factor of the confocal detection volume, and next, based on this result and on the known diffusion coefficients, (ii) the confocal detection volume in femto liter. Additionally, the (iii) molecular brightness of your fluorophores and finally, (iv) the concentration of the samples can be calculated based on the fit results.

###### Fit of a FCS curve

- Open Chisurf2016
  - It might take a while to open – be patient!
- It has got two panels (Figure 1):
  - Left / white: Here, the data is loaded and the analysis method / model is selected. Additionally, you can use the integrated Python-console to run little scripts.
  - Right / grey: Here, each data set opens in its own window.

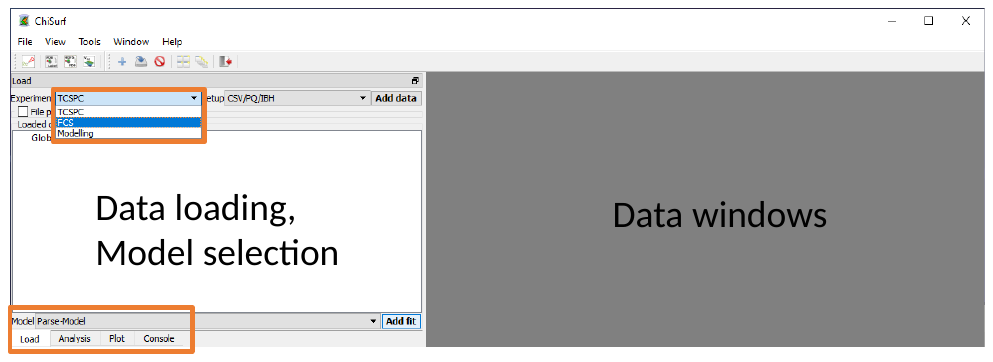

Figure 1 Start window of ChiSurf

- Change to the “FCS” mode und load your all “***A488_ACF_prompt.cor****”* (auto correlation curves of green calibration fluorophore correlated within the prompt time window) dataset(s) by “File”-> “Add dataset” or simply by “drag’n’drop” into the white area (Figure 2).

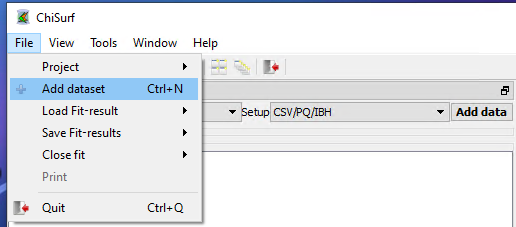

Figure 2 Adding new data sets

- Select your loaded dataset(s) and click on “Add fit” (Figure 3):

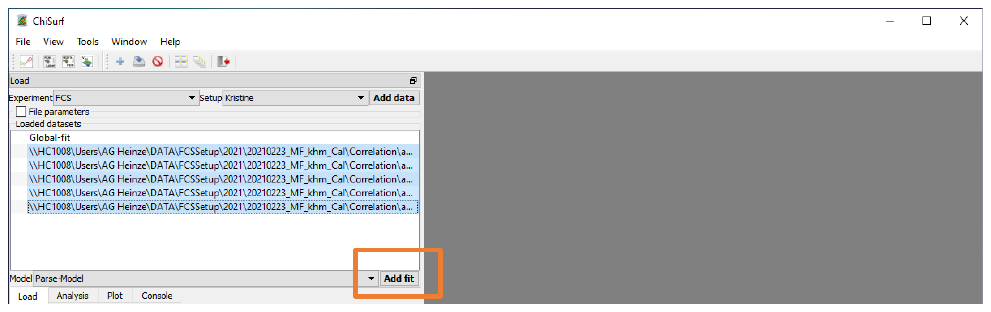

Figure 3 Adding new fits

- Here, five new windows will open in the grey working panel (Figure 4):

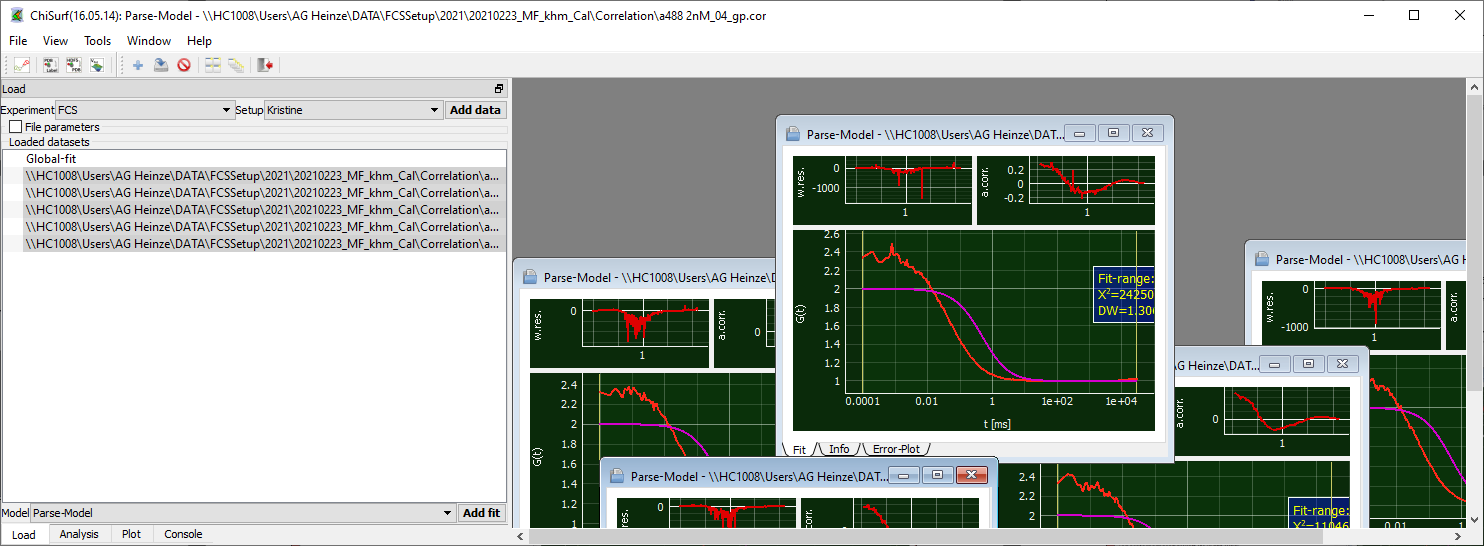

Figure 4 New fit windows open in the grey panel on the right side.

- Caution! The individual windows may lie on top of each other!
- Move the top-visible window aside and arrange the windows as you find it comfortable:
  - You can maximize the windows to see only one curve at a time. Minimize the window to see the other data again.
  - Alternatively, you can display each curve in full-size but in individual tabs (Figure 5). Then you can switch between the data by clicking on the respective tab and / or use the little arrows on the side to change between the tabs.

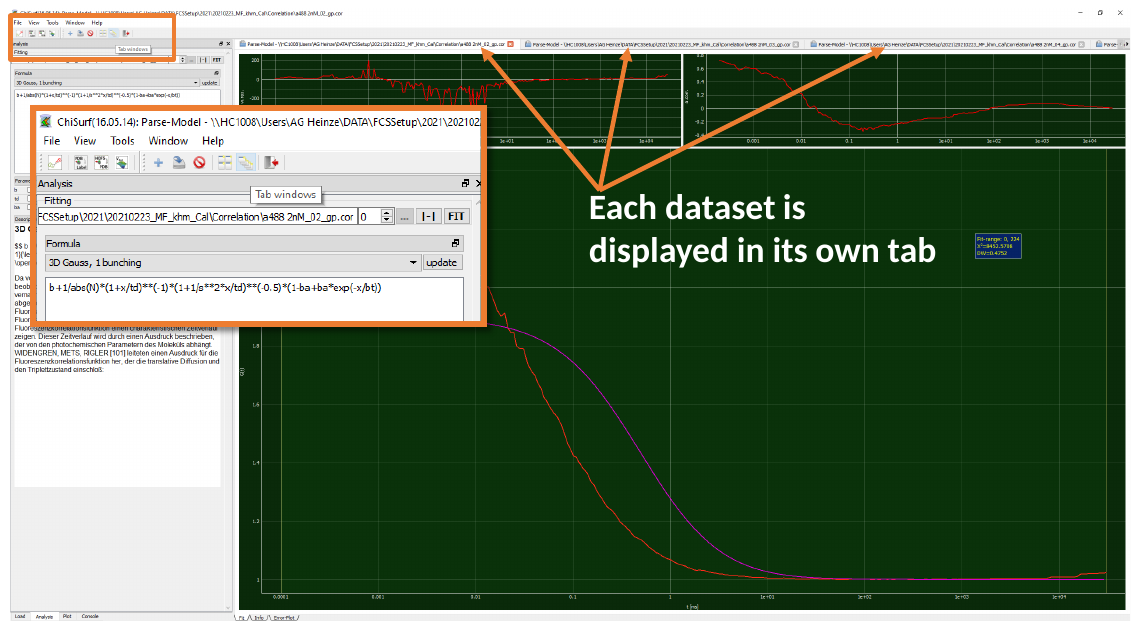

Figure 5 Fit windows of data sets arranged in individual tabs.

- Or you can distribute all windows automatically to evenly cover the full working panel (Figure 6):

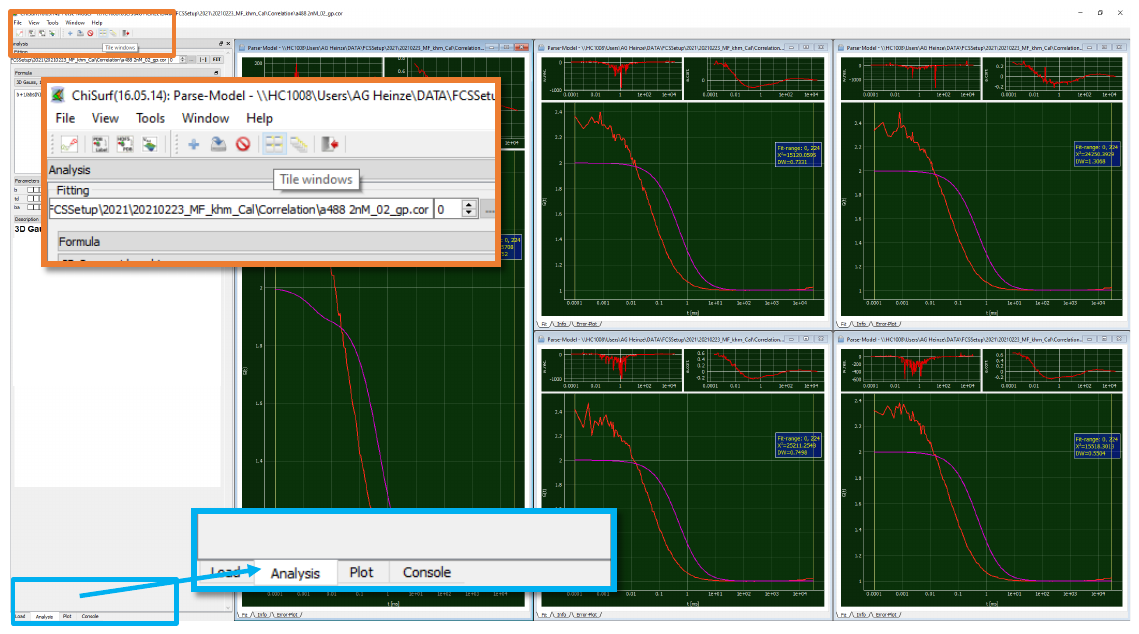

Figure 6 Fit windows of data sets distributed evenly over the data window pane.

- Now change to the analysis tab in the data panel and select the “3D Gauss 1 bunching” model from the drop-down menu for your first curve (Figure 7):

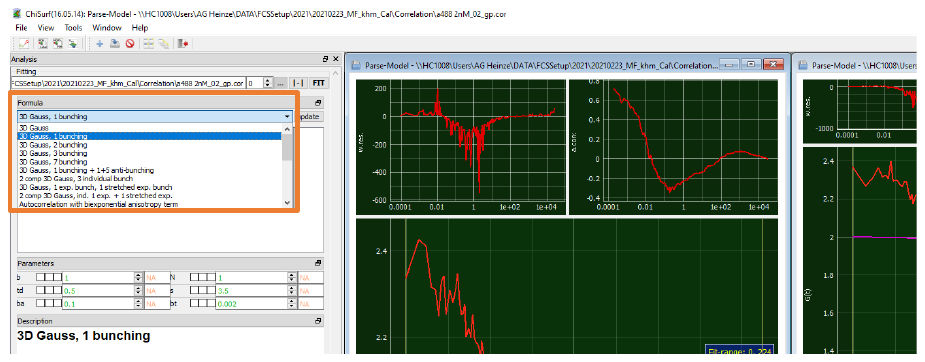

Figure 7 Selection of a fit model.

- This model has six parameters (Figure 8):
- The “1 Bunching” term as described by “ba” and “bt” is used to model the typical photophysical triplet blinking of many fluorophores in the µs time range.
  - Note: ChiSurf comes upon installation with a selection of pre-defined fit models, you can modify these fit models or add your own models easily.
  - This will be shown later.

| b | offset of the curve (either 1 or 0) | 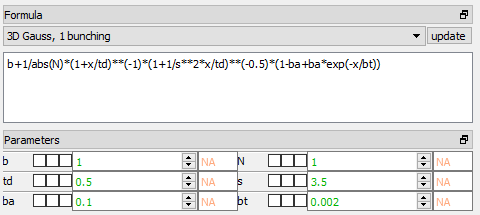 |
| --- | --- | --- |
| N | number of molecules in focus |  |
| td | diffusion time of the molecule [ms] |  |
| s | shape factor of the confocal volume:  s = z_0_/w_0_  (ratio height / width) |  |
| ba | amplitude of relaxation time |  |
| bt | time constant of relaxation time [ms] |  |
| Figure 8 Fit parameter of the 3D diffusion model containing an additional relaxation term. | | |

- Press “Fit” for fitting:

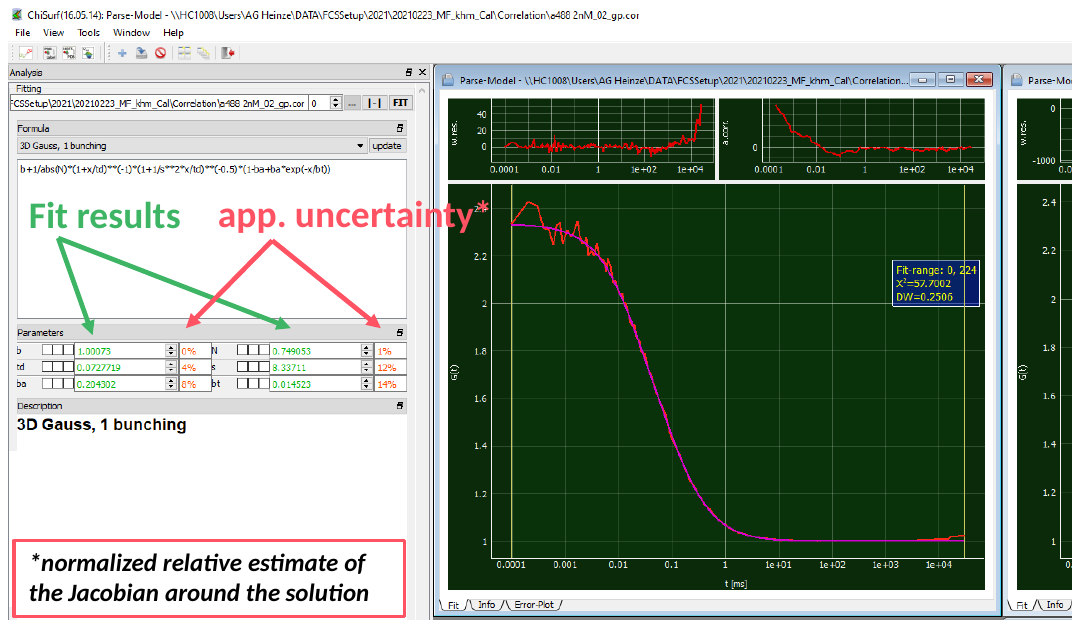

Figure 9 After the fitting, the obtained values for the parameter are shown and the apparent uncertainty is displayed. The small panel on top of the fit show the weighted residuals (left) and the autocorrelation of the weighted residuals (right)..

- We obtain a number of molecules in the focus N = 0.75, a diffusion time td = 0.073 ms (i.e. 73 µs) and triplet blinking time constant of 14.5 µs with an amplitude of 0.20. However, from the weighted residuals and the autocorrelation of the residuals, we can see a mismatch at long correlation times (Figure 9): This is because the absolute measurement time in this measurement was too short to reliable obtain these values.
- Next to the fit results, also the *normalized relative of the Jacobian Matrix around the solution* can be seen. This is **NOT** reflecting the **uncertainty** of the fit result, but the values might give a first hint whether the uncertainty is rather large or small. For more details on this topic, please check out the information provided on the following web page and the references cited herein:

<https://root.cern.ch/doc/master/Minuit2Page.html>

- For a more reliable estimate on the uncertainty of the fit parameter you have two options within ChiSurf to (i) sample the χ²-surface or (ii) to run a Markov-Chain Monte-Carlo simulation, which also allows you to obtain the mutual dependencies between the fit parameter. However, this uncertainty analysis is beyond the scope of this analysis of the calibration samples and will be shown in a different tutorial.
- Here, we take advantage of multiple measurements of the same sample and take these as additional restraints.
- Shorten the fit range to ~ 100 ms by grabbing the right yellow line and move it to the left (Figure 10):

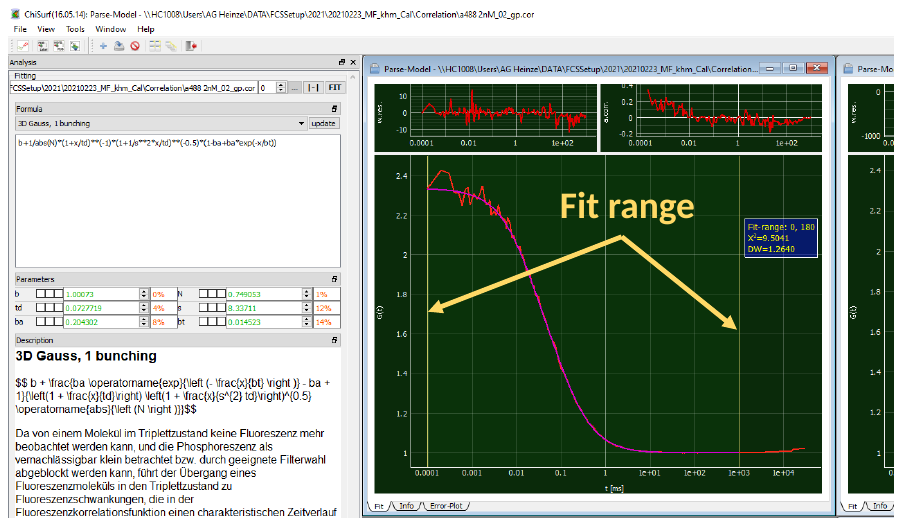

Figure 10 Modifying the fit range.

- *Of note*: To reliably fit your diffusion time, the **baseline** (0 or 1, depends on correlation algorithm) **MUST** **be reached** in your correlation curve (or in the fit range, respectively)
- Press “fit” and observe the changes:

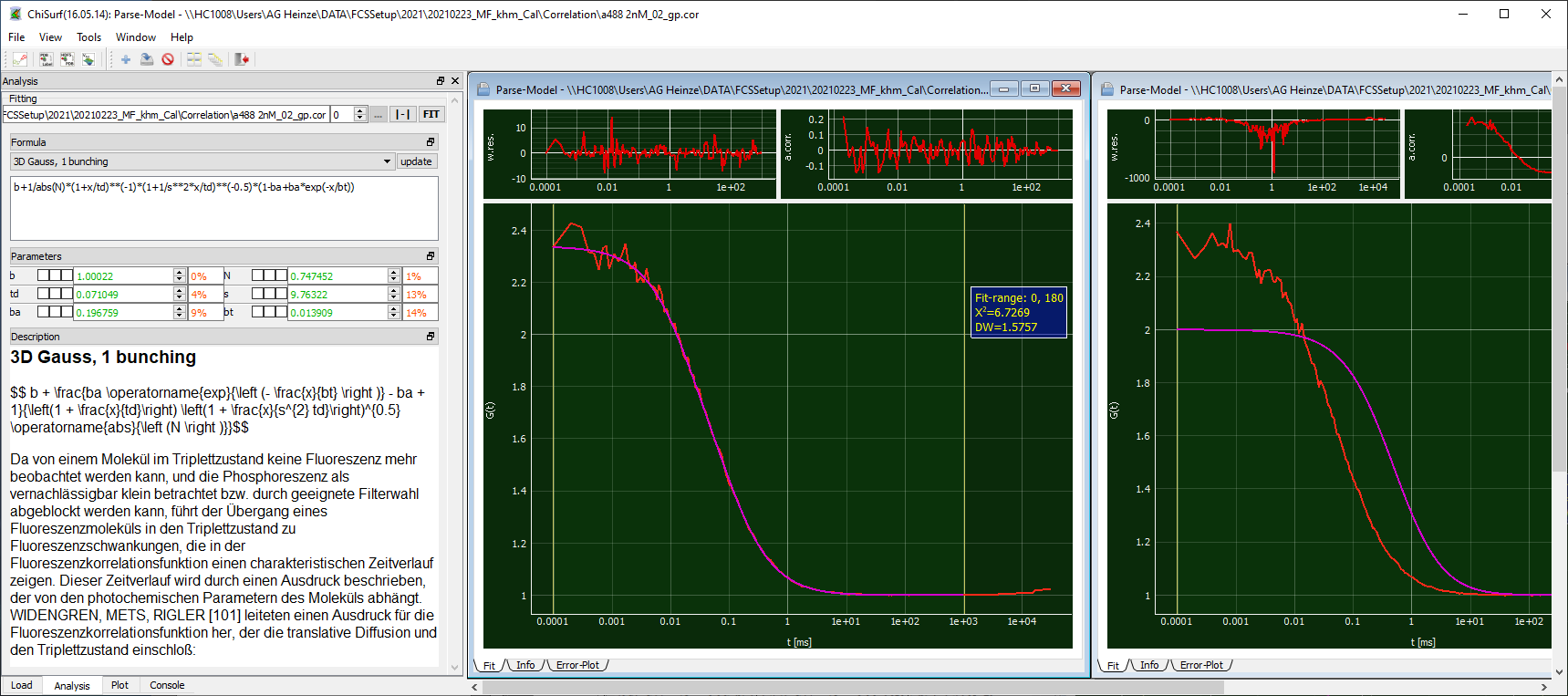

Figure 11 Individual fit of a single data set with the 3D diffusion model including one relaxation term with optimized fit range..

- Now the shape has increased to 9.76, which is very huge and would indicate a misalignment of your system. In ideal case, the shape factor should lie between 3 - 7. The other values have changed only slightly (Figure 11).
- Now let’s add the other measurements into the play and see whether a global fit of all measurements stabilizes this value.
- Go to the other fit windows, change the fit to “3D Gauss, 1 bunching”, adjust the fit range and fit them as done for the first curve (Figure 12):

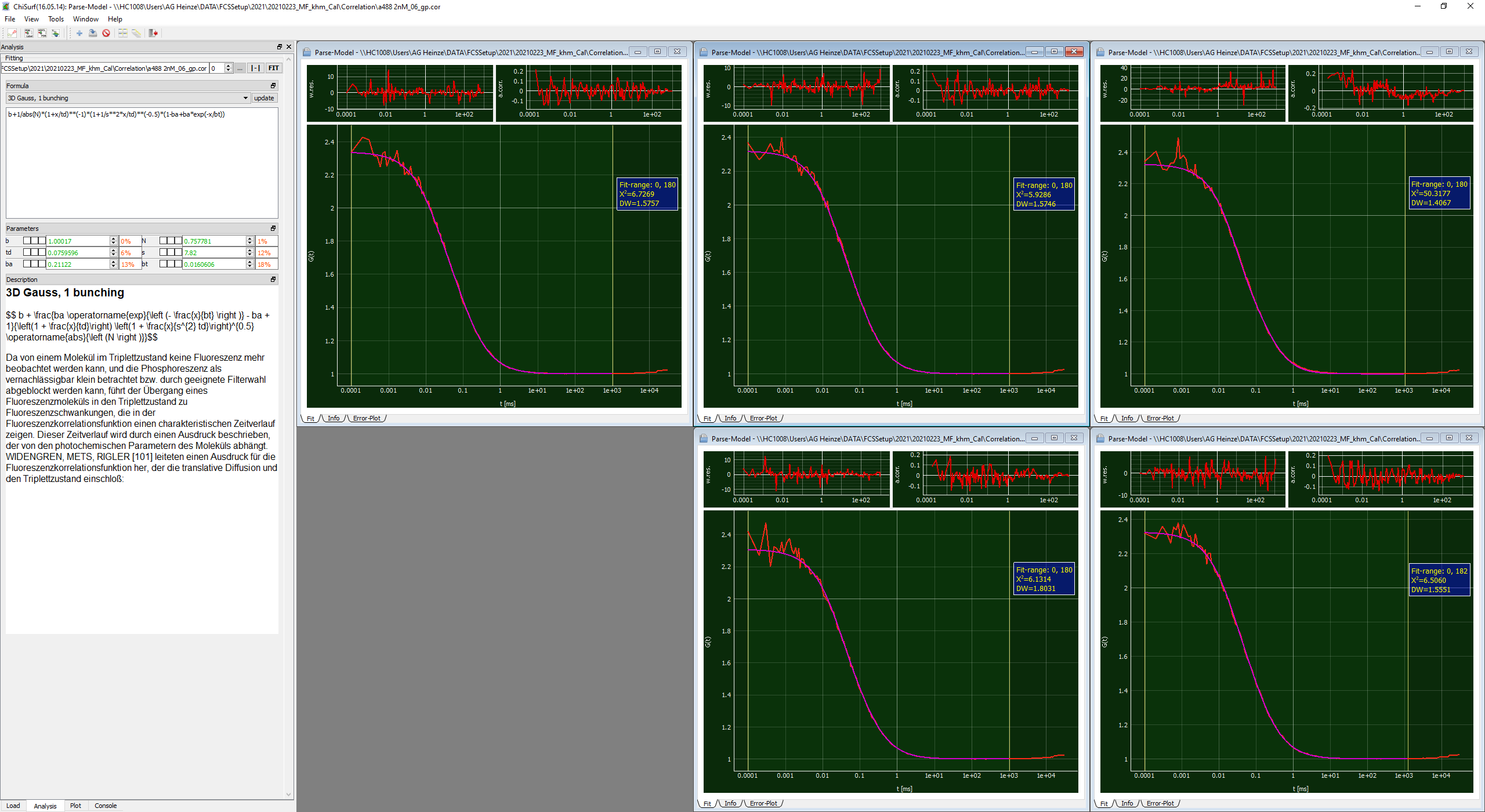

Figure 12 All five data sets of the A488 measurement were fitted individually with the 3D diffusion model including an additional relaxation term..

- Next, we will link the fits together such that the fit parameter are jointly minimized
- For this, first decide for one “parent” dataset, to which all other datasets are pointing. Here, I will simply take curve #1.
- Now, switch to the first of your “child” or “dependent” dataset. We will now work with the three checkboxes located between each variable name and variable value:

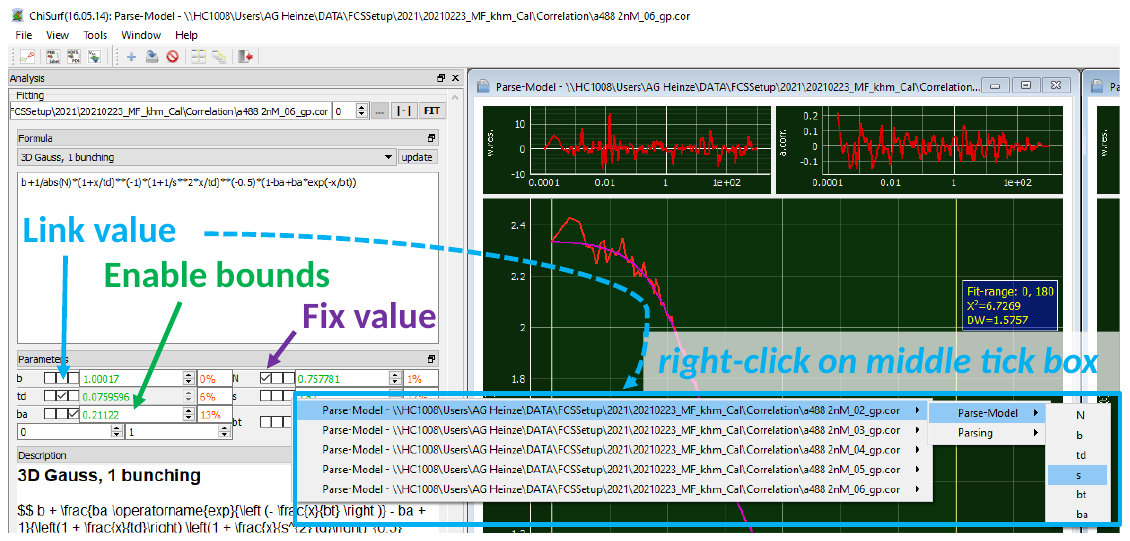

Figure 13 Linking the fit parameter in preparation for a global, joint fit.

- Each tick box has a different function (Figure 13):
  - Left: fixes the value of the parameter to its current value
  - Right: Two new parameter fields open, in these ones the lower (left) and upper (right) boundaries for this parameter during the fitting can be defined. An example is to define the allowed fit range for a correlation amplitude to be positive and lie between 0 -1.
  - Middle: By right-clicking into this tick box a list of all opened fit windows opens. Move your mouse towards the right, as soon as you approach the little arrow, new options appear, move further right on the height of “Parse Model” until the list of fit parameter appears.
- From this list of fit parameter select the appropriate one.
- For our global fit, we will now link (i) the diffusion time td, (ii) the shape parameter s and (iii) the triplet time constant bt to the first data set (Figure 14):

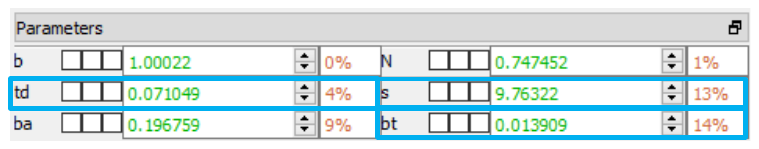

Figure 14 Global fit parameter: td, s and bt.

- Linked parameter will appear greyed out.
- After you are done with the linking, add a new global fit in the “load” tab:

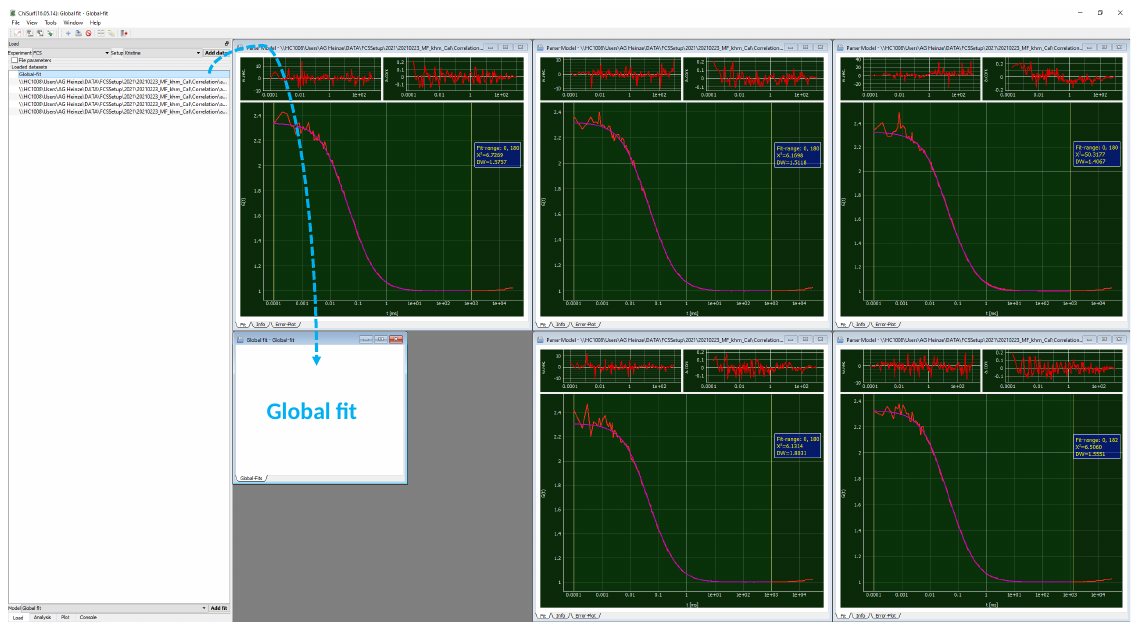

Figure 15 The fit window for the global fit parameter opens as a new, white window.

- It opens as empty / white window (Figure 15).
- Switch back to the analysis tab and add dataset which are to be jointly fitted by clicking firstly on “update” and then on “add”.

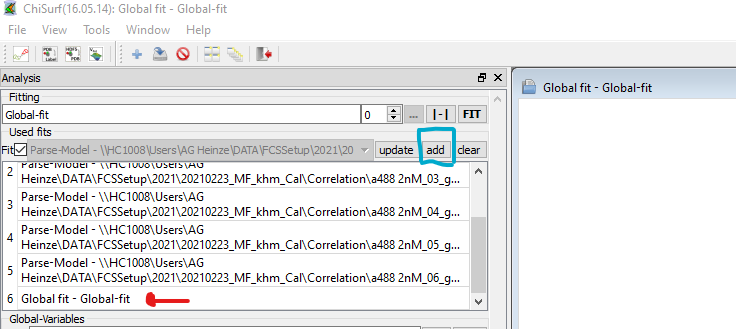

Figure 16 In a first step all open datasets are added to the global fit, next the global fit is removed from the list to avoid “circular reference”.

- Now a list of all available fit windows appears (Figure 16).
- If you want to select which datasets to add, remove the tick behind “fits”. Then you can choose from a drop-drown list which datasets are to be included in the global fit.
- We need to remove the “global fit” from list (“circular reference”). You do so by simply double-clicking on the item in the list.
- Now, press fit and observe what happens.
- If you fit many and / or complicated models, the program might take a few moments and display “not responding”. This is nothing to worry about and wait until it responds.
- Now all datasets have fitted jointly and we obtain the following values (Figure 17):
  - td = 85.7 µs
  - s = 5.84
  - bt = 21.0 µs
  - ba varies between 0.255 – 0.270
  - N varies between 0.748 - 0.766

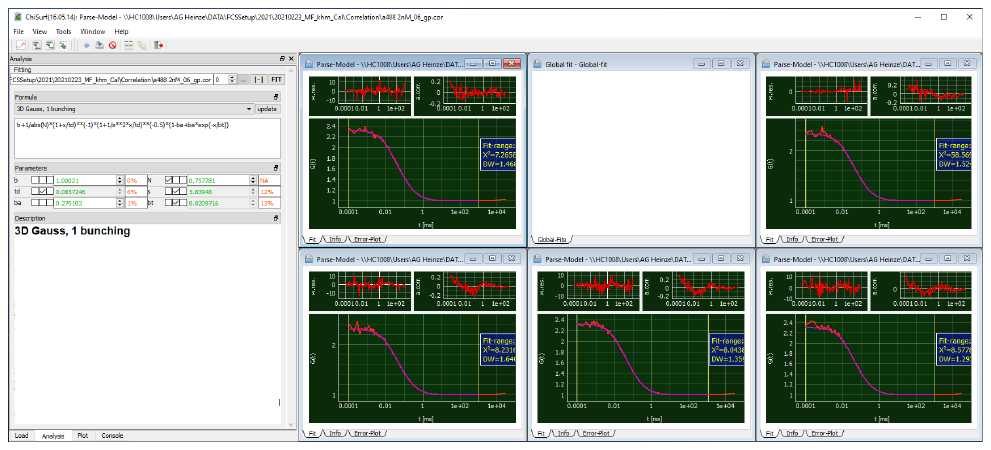

Figure 17 Result of the global fit.

- Finally, let’s save the fits by either selecting “File” -> “Save Fit-results” -> “current fit” or by pressing “Ctrl + S” (Figure 18).
- Caution! Saving all fits may not work, if the filenames are (a) similar and (b) the whole file path is too long. (AutoSaving uses complete path as automatic save name currently).

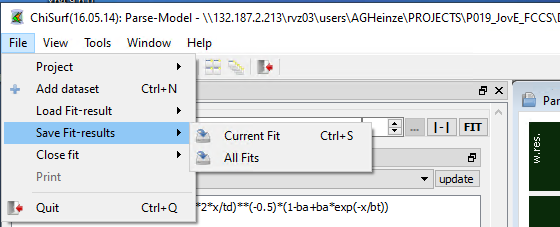

Figure 18 Saving of the fit results.

- Save the results of all measurements, we will need the fit results in the next step.

###### Calculation of confocal volume

Note:

- Use the provided excel sheet “*FCCS_calibration.xlsx*” to retrieve the results semi-automatically (Figure 19).
- Don’t forget to update each time your fit results from the respective measurement day!
- Please note the different units (µm, m etc.) and unit conversions!
- To determine the confocal volume, we need three different values:
  - Diffusion coefficient of the freely diffusing standard dye, here A488,
    - These values can be found in the literature for most common fluorophores.
    - Caution! They are temperature and solvent dependent, i.e. if you measure at low temperature or in a more viscous environment, you have to correct for the viscosity of the solution.
    - **D_A488_ = 414 µm²/s** (@25°C in ddH2O ^2^
  - Diffusion time ***t_D_*** from our fit: **85.7 µs**
  - Shape factor ***s*** from our fit: **5.84**
- Diffusion time and diffusion coefficient are related by the following relationship:

$D= \frac{w_{0}^{2}}{4t_{D}}$	🡪	$w_{0}^{2}=4t_{D}D$

- Inserting our fit results and *D* into this equation, we obtain ***w_0_* = 0.377 µm**
- Based on the value from ***w_0_***, we can determine the height ***z_0_*** of the confocal volume:

$s= \frac{z_{0}}{w_{0}}$	🡪	$z_{0}=s w_{0}$

- For our example, ***z_0_* = 2.2 µm**
- Finally, the confocal volume is assumed to have in good approximation an elliptical shape from which the volume ***V_eff_*** can be obtained using the following formula:

$$V= \pi^{3/2}z_{0}w_{0}^{2}$$

- Thus, our confocal volume in the green excitation range has a size of ***V_eff,green_* = 1.74 fL**.

###### Estimation of the fluorophore concentration

- Using the above determined confocal volume, the concentration of the fluorophore in your measurement solution can be estimated.
- The following parameters are required:
  - Confocal volume ***V_eff,green_* = 1.74 fL**
  - Number of molecules in focus, ***N***, from our fit: **0.76** (in average)
  - Avogadro’s number: ***N_A_*** = **6.022*10^23^ mol^-1^**
- The concentration of the fluorophore is determined from the following relationship:

$$c= \frac{N}{V_{eff,green}N_{A}}$$

- The approximated concentration of green calibration dye ***c_A488_* ~ 0.72 nM.**

| 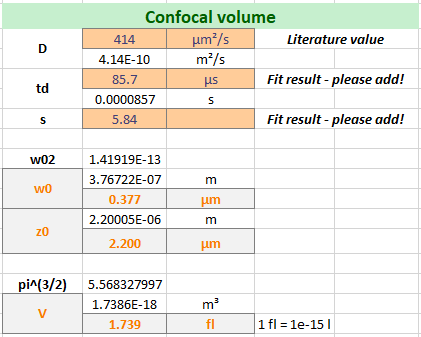 | 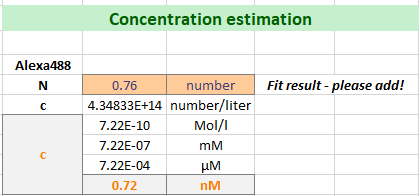 |
| --- | --- |
| Figure 19 Calculation of the size of the green confocal volume and the concentration of the Alexa488 solution. | |

##### Calibration of detection volume – red excitation

- For the red detection volume, we proceed as described for the green detection volume.
- Only the fit results will be provided for a guidance (Figure 20).
- Important: Use the correlated signal of the directly excited red fluorophore within the delay time window of your measurement, e. g. “***A568_ACF_delay.cor***”

###### Fit of FCS curve

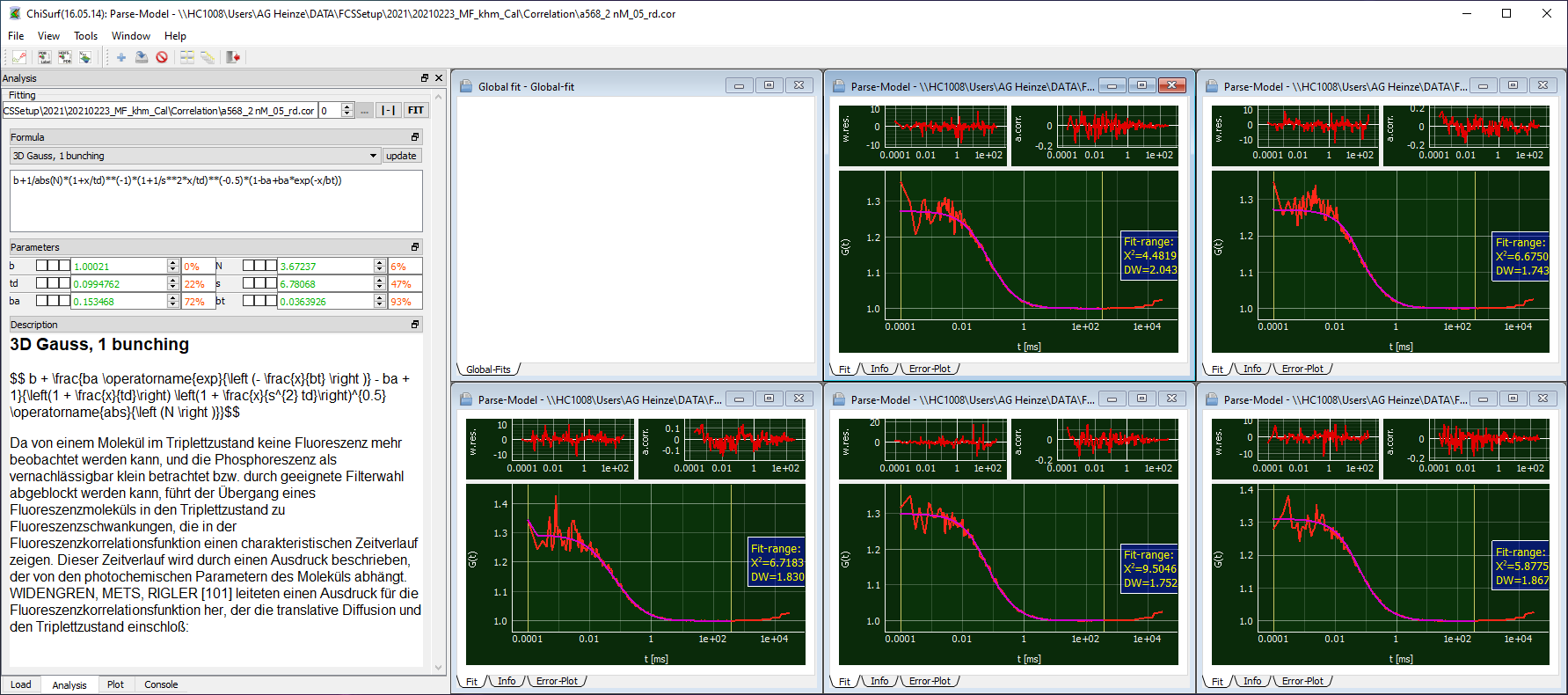

Figure 20 Global fit of the Alexa568 solution.

- Here, also the “3D Gauss, 1 Bunching” model will be sufficient.
- We obtain:
  - td = 99.5 µs
  - s = 6.78
  - N varies between 3.16 – 3.67
- Save the fit results and proceed with the calculations as described for the green detection volume.

###### Calculations

- As A568 is larger than A488, its diffusion coefficient is reduced compared to A488.
- There are different studies reporting a ***D_A568_*** in the range of 330 -360 µm²/s.
- Here we use a value of ***D_A568_* = 363 µm²/s** ^3^.

| 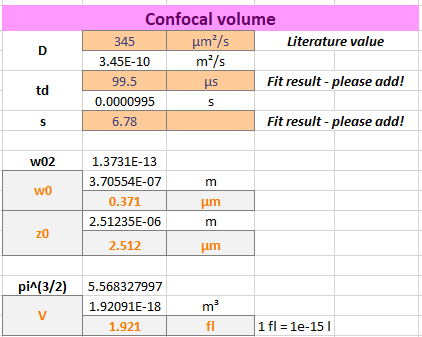 | 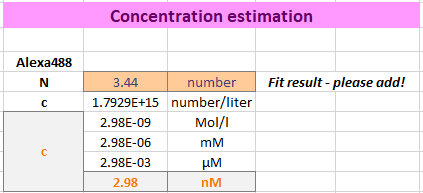 |
| --- | --- |
| Figure 21 Calculation of the size of the red confocal volume and the concentration of the Alexa488 solution. | |

- As expected for a measurement at red-shifted, thus longer wavelengths of excitation and emission, the confocal detection ***V_eff,red_*** volume is increased compared to ***V_eff,green_*** (Figure 21).

##### Correction factors for spectral crosstalk & direct acceptor excitation

- Absorption and emission spectra of fluorophores are usually quite broad and extent over quite a wavelength range, and not seldom the spectra from green and red fluorophores used in the measurement overlap (Figure 22).
- By introducing optical (bandpass) filters into the emission path of your setup, it is possible to minimize these effects.

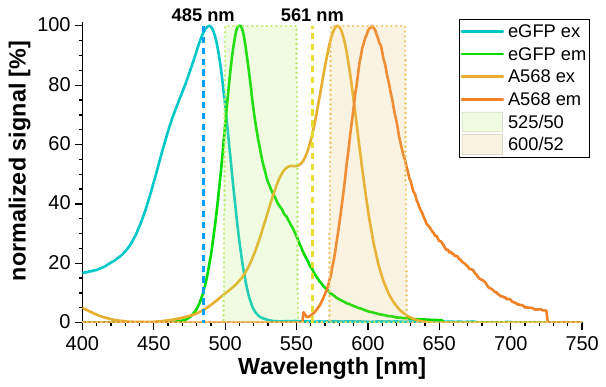

Figure 22 Normalized absorption and emission spectra for the eGFP and Alexa568 fluorophores used in the cell experiments. Excitation laser lines are shown as dashed lines and transparent boxes denote the bandpass filter through which the emitted fluorescence is collected.

- However, even if these contributions to your signal are small, they might be become critical when you have a low count rate / low signal in one channel compared to the other channels.
- For more detailed information on required corrections and the nomenclature of correction factors, we refer to the recently published multi-laboratory benchmark study of Hellenkamp *et al.* ^4^.

###### Spectral crosstalk of green fluorescence into red detection channel

- Required is the measurement from the freely diffusing green fluorophore.
- Collect the count rates in all four detection channels for the ddH2O /buffer measurement and the green dye (A488) measurement:

| Channel number | Count rate [kHz] BG | Count rate [kHz] A488 | Corrected count rate [Hz] |
| --- | --- | --- | --- |
| 0 | 0.13 | 5.30 | 5.18 |
| 1 | 0.36 | 1.23 | 0.87 |
| 2 | 0.23 | 5.38 | 5.15 |
| 3 | 0.69 | 1.53 | 0.85 |

(Note: average values of repeated measurements are reported)

- The background count rate needs to be subtracted from value and then the ratio of count rates in the red channels to the green channel is determined:

$$\alpha= \frac{(\left( {CR}_{1,A488}-{CR}_{1,BG} \right)+\left( {CR}_{3,A488}-{CR}_{3,BG} \right))}{(\left( {CR}_{0,A488}-{CR}_{0,BG} \right)+\left( {CR}_{2,A488}-{CR}_{2,BG} \right))}*100\%$$

- In our example the spectral crosstalk ***α* = 16.6 %.**

###### Determination of direct excitation of red fluorophore by green excitation

- Required is the measurement of the freely diffusing red fluorophore.
- Collect the count rates in the two red detection channels for the ddH2O / buffer measurement and the red dye measurement A568, take care to separate the count rate into the “prompt” and “delay” time window:

| Channel number | Count rate [kHz] BG | Count rate [kHz] A568 | Corrected count rate [Hz] |
| --- | --- | --- | --- |
| 1 delay | 0.36 | 4.24 | 3.88 |
| 1 prompt | 0.36 | 1.90 | 1.54 |
| 3 delay | 0.69 | 4.81 | 4.12 |
| 3 prompt | 0.69 | 2.37 | 1.68 |

(Note: average values of repeated measurements are reported)

- In the “prompt” time window the green excitation pulse might excite a bit of the red fluorophores.
- In the “delay” time window the red fluorophores are directly excited.
- Subtract the background counts from each channel and the respective time window.
- Calculate the ratio of the “prompt” time window to the “delay” time window:

$$\delta= \frac{(\left( {CR}_{1,prompt, A568}-{CR}_{1,prompt,BG} \right)+\left( {CR}_{3,prompt,A568}-{CR}_{3,prompt,BG} \right))}{(\left( {CR}_{1,delay, A568}-{CR}_{1,delay,BG} \right)+\left( {CR}_{3,delay,A568}-{CR}_{3,delay,BG} \right))}*100\%$$

- Caution! If you measure in a mixture of green and red fluorophores, you need to also subtract the green crosstalk into the red channels (α determined above!)
- For the shown example, the direct excitation of acceptor by green laser is ***δ* = 40.3 %.**

##### Determination of molecular brightness

- After having correcting the count rates, we can now determine the molecular brightness of our fluorophores (Figure 23).
- Note: It is advisable to monitor this molecular brightness of your calibration standards carefully as they can give you early hints about a possible misalignment and / or reduction of laser output power.
- Required parameter:
  - Number of molecules in focus as determined from the fits above
    - ***N_green_***: 0.76
    - ***N_red_***: 3.44
  - Corrected count rates:
    - ***CR_corr,green,0_*** = 5.18 kHz
    - ***CR_corr,green,2_*** = 5.15 kHz
    - ***CR_corr,red,1_*** = 3.88 kHz
    - ***CR_corr,red,3_*** = 4.12 kHz
- The molecular brightness is calculated as count per molecule and second:

$$B_{green}= \frac{{CR}_{corr,green,0}+{CR}_{corr,green,2}}{N_{green}}$$

$$B_{red}= \frac{{CR}_{corr,red,1}+{CR}_{corr,red,3}}{N_{red}}$$

- For the data shown here, the following molecular brightness is obtained:

| 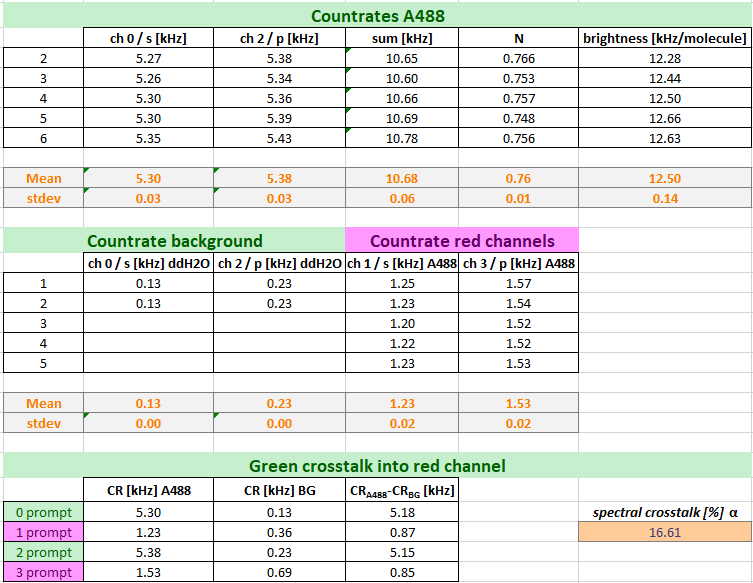 | 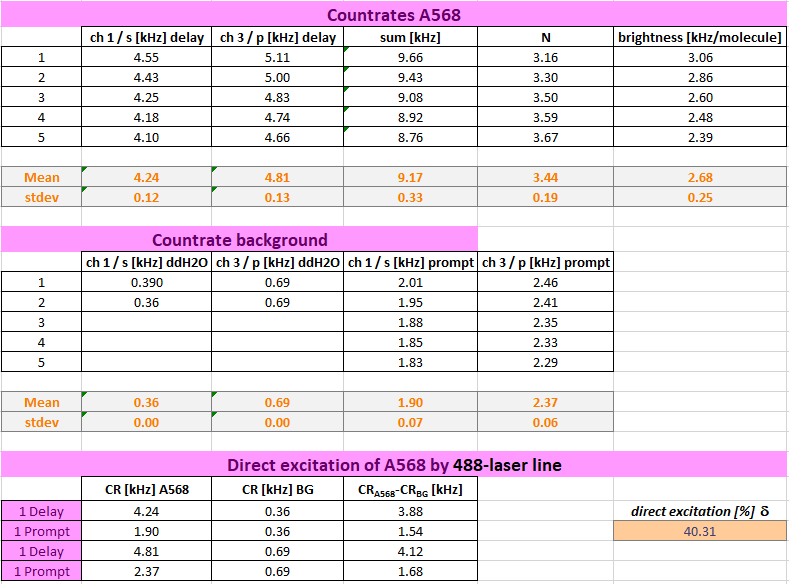 |
| --- | --- |
| Figure 23 Calculation of the molecular brightness of both calibration standards.. | |

##### Determination of overlap of green & red detection volume

- For calibration of the overlap of the green and red excitation volume, DNA strands are used, which carry both the green and red calibration fluorophore. However, the fluorophores are placed usually so far apart that no energy transfer due to FRET can occur.
- For single-molecule measurements, a mixture of DOnly-labeled DNA strands and high- and low-FRET showing DNA strands is additionally used to calibrate the green-to-red detection efficiency ratio.
- For our determination of the confocal volume overlap, we will use the amplitude of the green (ACF_green_) and red autocorrelation function (ACF_red_) as well as from the “Green-prompt”-“Red-delay” cross-correlation function (CCF_PIE_).

***Important note!***

The detection volume increases with the excitation and emission wavelength of the fluorophores. This effect is more prominent in diffraction-limited setup as used here and commonly for live cell FCS.

In setups for single-molecule experiments, with freely diffusing molecules, with larger detection volumes this effect can often be ignored and the diffusion terms in the green and red channels can be fit jointly.

###### Global fit of FCS curves

- Switch back to ChiSurf2016 and load the following correlation curves from your DNA-sample:
  - *DNA_gp.cor*: Autocrrelation of green channels in prompt time window
  - *DNA_rd.cor*: Autocorrelation of red channels in delay time window
  - *DNA_PIE.cor*: Crosscorrelation of green signal in the prompt time window with red signal in the delay time window
- Add a “3D Gauss, 1 bunching” model to your correlation functions
  - Of note: theoretically no bunching term should be required as the photophysics from the two fluorophores is independent from each other, and is thus not resulting in a correlating signal.
  - However, in practice, in case of significant photophysics / triplet blinking and considerable crosstalk as is the case here, we often observed an apparent “photophysics” term, which we model by the relaxation term.
- Fix the shape factor ***s_green_*** and ***s_red_*** to your determined values from the free dye measurements above.
- Remember to adjust your fit range at long lag times if required.
- Note: If you have multiple DNA measurements, fit the respective correlation curve from the same correlation channels jointly, i.e. link ***td***, and ***bt*** and – for the CCF_PIE_ also ***s_PIE_***
- Observe ***s_PIE_*** in your CCF_PIE_, it should get a “reasonable” number, usually between ***s_green_*** and ***s_red_*** (Figure 24).
- Save all fit results, we will need them for our calculations below.
- Here, we obtain the following fit results for our DNA samples:

| Parameter | DNA_gp | DNA_rd | DNA_PIE | |
| --- | --- | --- | --- | --- |
| *s** | 5.7* | 6.78* | 4.63 | |
| *td* [µs] | 438 | 474 | 520 | |
| *N* | 11.6 – 7.54 | 14.4 – 8.73 | 1/G(tc) | 20.4 – 13.4 |
| *bt* [µs] | 88.6 | 101 | 170 | |
| *ba* | 5 - 22 % | 1.3 – 24.5 % | 0 – 15.7 | |

**s_green_ and s_red_ are fixed from the calibration measurements above.*

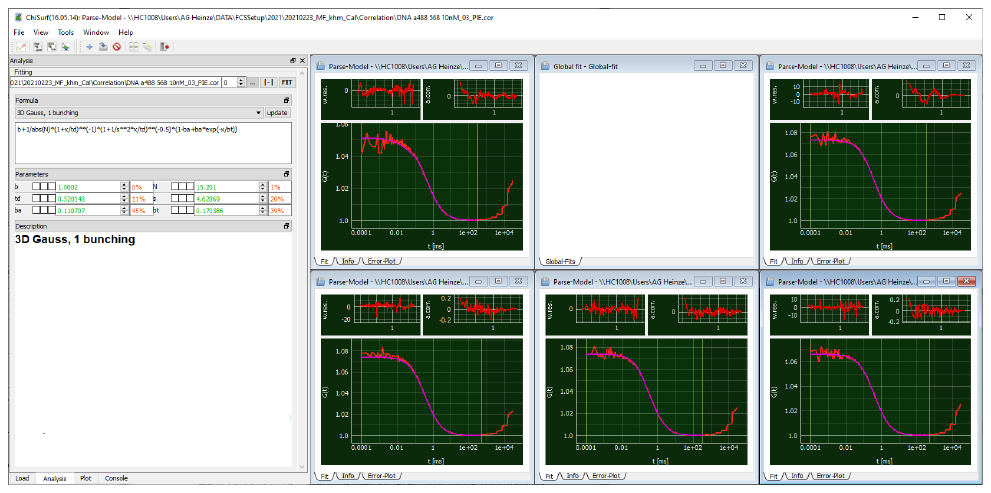

Figure 24 Global fit of the CCF_PIE_ correlations from the DNA sample..

###### Calculations

From our double-labeled DNA measurements, we need to derive two important parameter:

- The size of the overlapping confocal detection volume *V_eff,PIE_*
- The Cross-correlation amplitude, which reflects 100 % co-diffusion

The size of the overlapping confocal detection volume can be determined using the already provided equations used in the sections above for the calculations of the confocal detection volume of the green and red channel.

In a first step, we use the obtained diffusion times from our *DNA_gp* and *DNA_rd* fits and determine the translational diffusion coefficient of our DNA sample:

$D_{DNA,green}= \frac{w_{0}^{2}}{4t_{D,green}}$	and	$D_{DNA,red}= \frac{w_{0}^{2}}{4t_{D,red}}$

Here, we obtain a value of ***D_DNA,green_* = 81.8 µm²/s** and ***D_DNA,red_* = 72.4 µm²/s** using the value of w_o_ from A488 and A568 dye, respectively. Thus, in average ***D_DNA_* = 77.1 µm²/s**.

Based on this value, we can obtain *w_0,PIE_* and *z_0,PIE_*:

$w_{0}^{2}=4t_{D}D$	and	$z_{0}=s w_{0}$

Here, ***w_0,PIE_* = 400 nm** and ***z_0,PIE_* = 1.85 µm**. This results in a ***V_eff,PIE_*** of **1.66 fL**:

$$V= \pi^{3/2}z_{0}w_{0}^{2}$$

Next, we observe the amplitudes of the auto- and cross correlation functions: In an ideal system the amplitudes of the three curves, *DNA_gp*, *DNA_rd* and *DNA_PIE* should be identical. However, as the detection volumes differ with the excitation and emission wavelength, this is rarely the case. In the next-optimal setting, the amplitude of *DNA_PIE* would be identical to the amplitude of the autocorrelation curve with the lower amplitude.

In common experimental settings, the overlap of the green and red confocal detection volumes is suboptimal and the apparent amplitude of a 100 % co-diffusion sample is required for calibration.

The concentration, and thus, later the fraction of co-diffusing particles in your sample, of double-labeled particles can be calculated based on the ratio of the correlation amplitudes:

$c_{RG}=\frac{G_{0,CCF}}{G_{0,ACFgreen}}\cdot c_{green}$	and	$c_{GR}=\frac{G_{0,CCF}}{G_{0,ACFred}}\cdot c_{red}$

where the amplitudes *G_0,ACFgreen_* and *G_0ACF,red_* are the inverse of the respective number of particles, *N_green_* and *N_red_*, in focus.

Here, we obtain amplitude ratios for 100 % co-diffusion of ***ratio_GR_* = 0.57** for the green and of ***ratio_RG_* = 0.68** for the red autocorrelation curves (Figure 25).

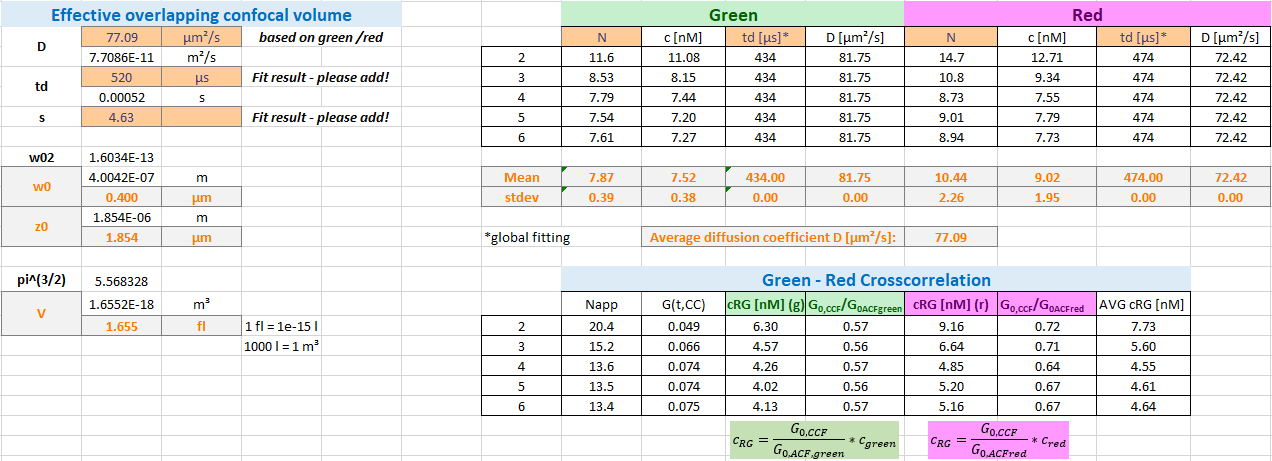

Figure 25 Calculation of the effective overlap volume and the correction factor for the co-diffusion amplitude.

Now we are ready to switch to our real samples measured in live cells.

#### Analysis of live cell experiments

Again, also here, we assume that you have already correlated your data using the software of your system or any of the software mentioned in the beginning.

We will show you first how to add your own fit models to ChiSurf and next analyze first the measurements of the singly labeled constructs (β_2_AR-eGFP-IL3 and NT-SNAP-β_2_AR) before switching to the double-labeled NT-SNAP-β_2_AR-eGFP-IL3 sample.

##### Adding the membrane-diffusion models

Upon installation, ChiSurf comes with a bunch of FCS fit models; however, your required fit model might not be among them. The fit models are defined in a JSON-file (“fcs.model.json”) which can be found in the installation folder of ChiSurf (Figure 26):

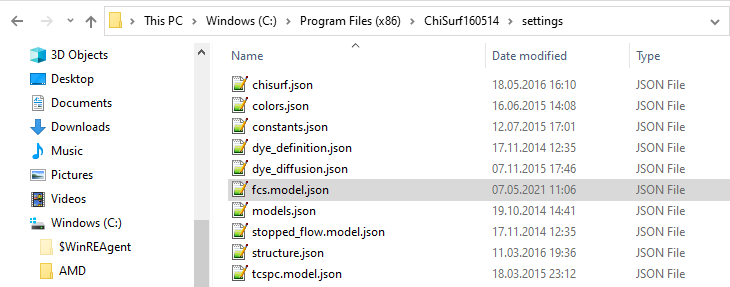

Figure 26 The settings file containing the fit models can be found directly in the installation folder.

Before modifying the file, (i) create a copy on a different place as a backup and (ii) make a second copy to work on as modifying / saving directly in the programs installation folder is usually not allowed.

Open the JSON-file using a text editor, e.g. Notepad++.

Each fit model consists of four sections (Figure 27):

- Model name
- Model equation
- Model parameter definition by initial values
- Model description

*It is vital to keep this notation and take care of proper punctuation and indentation!*

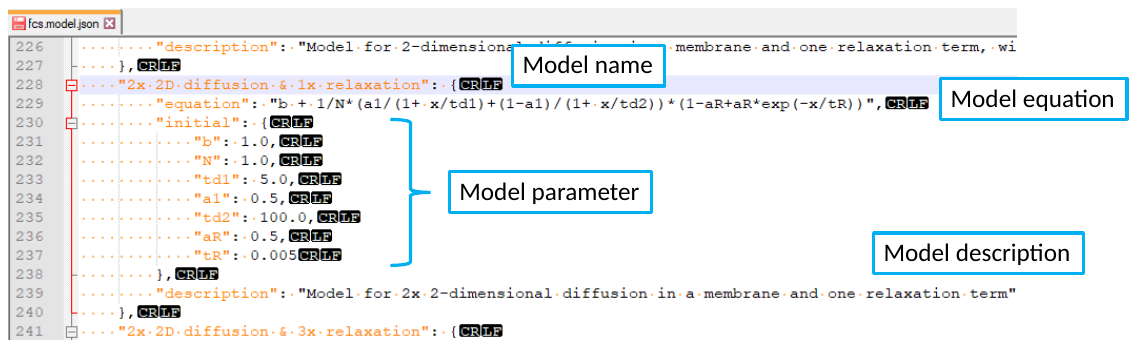

Figure 27 JSON-format of the fit models.

Add the two following fit models for bimodal membrane diffusion with or without and additional relaxation / triplet term to your JSON-file:

- For analysis of autocorrelation curves:

$$G_{ACF,2D}\left( t_{c} \right)=b+\frac{1}{N}\left[ \frac{a_{1}}{1+\frac{t_{c}}{t_{D1}}}+\frac{1-a_{1}}{1+\frac{t_{c}}{t_{D2}}} \right] \left[ 1-a_{R}+a_{R}\cdot\exp\left( -\frac{t_{c}}{t_{R}} \right) \right]$$

where *t_D1_* and *t_D2_* are the two diffusion time and *a_1_* is the fraction of *t_D1_*. *a_R_* and *t_R_* describe the triplet blinking / photophysics.

- For analysis of cross-correlation curves:

$$G_{CCF,2D}\left( t_{c} \right)=b+\frac{1}{N}\left[ \frac{a_{1}}{1+\frac{t_{c}}{t_{d1}}}+\frac{1-a_{1}}{1+\frac{t_{c}}{t_{d2}}} \right]$$

In cross-correlation curves, usually no triplet blinking can be seen.

Don’t forget to define reasonable initial values for each of the model parameter.

Replace the original JSON-file in your programs folder with your modified version and restart ChiSurf.

*Note: For changes to the JSON-file to take effect, ChiSurf must always be restarted!*

##### Individual transfected β_2_AR-eGFP-IL3 and NT-SNAP-β_2_AR

###### Fit of autocorrelation curves

Open the autocorrelation of the green channels from the prompt time window from the β_2_AR-eGFP-IL3 measurements and the autocorrelation of the red channels in the delay time window from the NT-SNAP-β_2_AR measurements in ChiSurf.

Select the bimodal membrane diffusion model for autocorrelation curves, which we have just added to ChiSurf and fit the data (Figure 28):

For average of the β_2_AR-eGFP-IL3 measurements, a slow diffusion time *t_D1_* = 118 ms and a fraction of 54 % is obtained. The faster diffusion lies at *t_D2_* = 1.9 ms. Additionally, 25 % triplet blinking at *t_R_* ~ 9 µs is observed. However, at this short correlation times the data is already quite noisy and care should be taken in the interpretation. The number of molecules in focus is 5.

For the average of the NT-SNAP-β_2_AR measurements, two additional relaxation terms seem to be required (modify your JSON-model file accordingly!) with relaxation times (and fractions) of *t_R1_* ~ 5 µs (12 %) and *t_R2_* ~ 180 µs (11%). The two diffusion components show times of *t_D1_* = 49 ms (49 %) and *t_D2_* = 2.7 ms. The number of molecules in focus lies at 35 and this is much higher compared to β_2_AR-eGFP-IL3. One could speculate whether *t_R2_* ~ 180 µs might not be a photophysics-related term but rather unreacted SNAP substrate diffusing through the confocal volume.

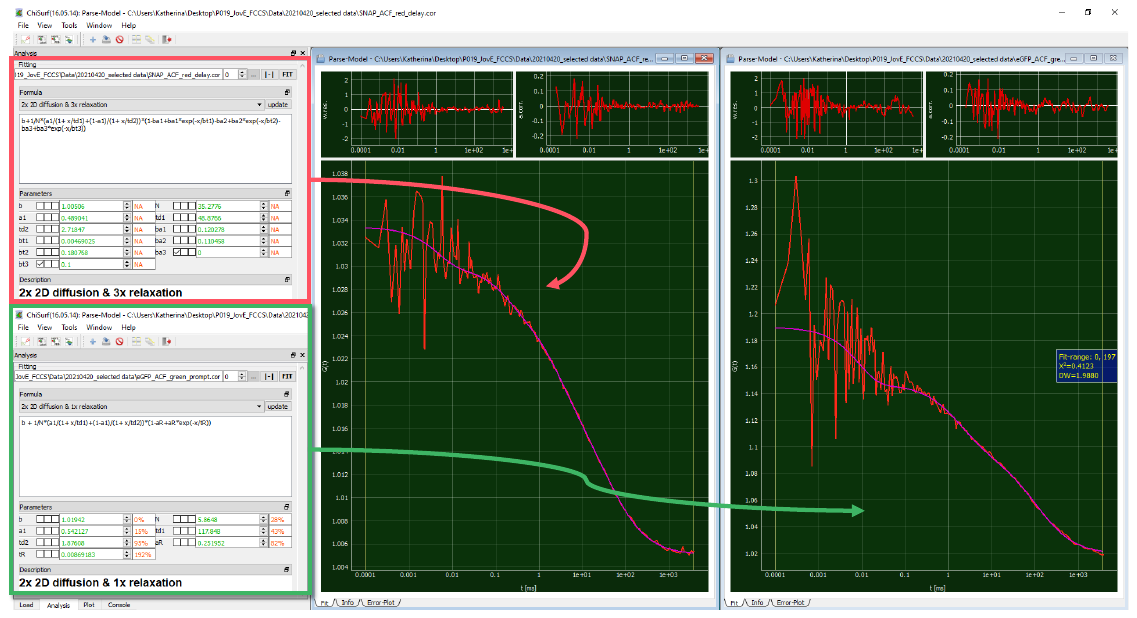

Figure 28 Fitting of the single-labeled constructs (red: NT-SNAP-β_2_AR, green: β_2_AR-eGFP-IL3) with 2x 2D diffusion model and additional relaxation times..

Based on the obtained number of molecules in focus and the known average count rates, we can also determine the molecular brightness of our fluorophores in the live cell settings and estimate the concentration of molecules using the equations explained in the calibration section.

Take care to subtract the background signal e.g. measured on non-transfected cells from the average count rate of your fluorescence samples (Figure 29).

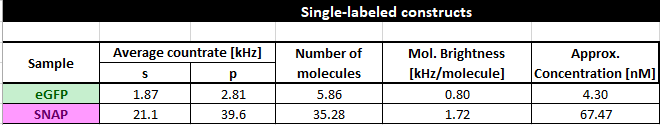

Figure 29 Characterization of the single-labeled constructs

###### Crosstalk-induced correlations

Based on these single-color experiments also one can test how much artificial / unwanted cross-correlation into the respective other color channel is present.

For this, we export and plot the following correlations:

- From the β_2_AR-eGFP-IL3 measurements: red channels in the delay time window and PIE cross-correlation (green channel prompt time window with red channels in the delay time window)
- From the NT-SNAP-β_2_AR measurements: green channels in the prompt time window and PIE-cross-correlation (green channel prompt time window with red channels in the delay time window)

In ideal case, all of these combinations show flat curves or better-said noise distributed around 1. Here, this is the case only for the PIE-cross-correlations (Figure 30). The respective “false-color” autocorrelation functions, red-delay from β_2_AR-eGFP-IL3 and green-prompt from CT-SNAP-β_2_AR, reflect the crosstalk into the red channels and the direct excitation of red fluorophore by the green excitation wavelength, respectively.

| **β_2_AR-eGFP-IL3** | **NT-SNAP-β_2_AR** |
| --- | --- |
| 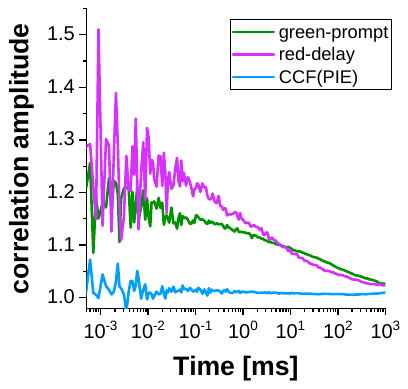 | 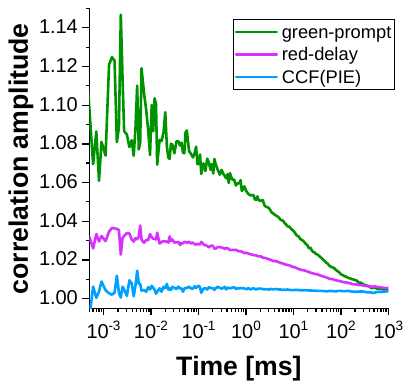 |
| Figure 30 Autocorrelations and the PIE-based crosscorrelation of the single-labeled constructs. | |

##### Double-labeled sample: NT-SNAP-β_2_AR-eGFP-IL3

###### Fit of auto- and cross-correlation curves

Open all three average curves, the green-prompt autocorrelation, red-delay autocorrelation and PIE-cross-correlation curve in ChiSurf. For both autocorrelation curves, we add the fit model with bimodal membrane diffusion and an additional relaxation term. For the PIE-cross-correlation curve, a bimodal membrane diffusion model is sufficient (Figure 31).

For the red autocorrelation and especially the PIE-cross-correlation, the fit ranges have to be adjusted as particularly for short correlation curves the noise is very high and no reliable fit could be acquired here.

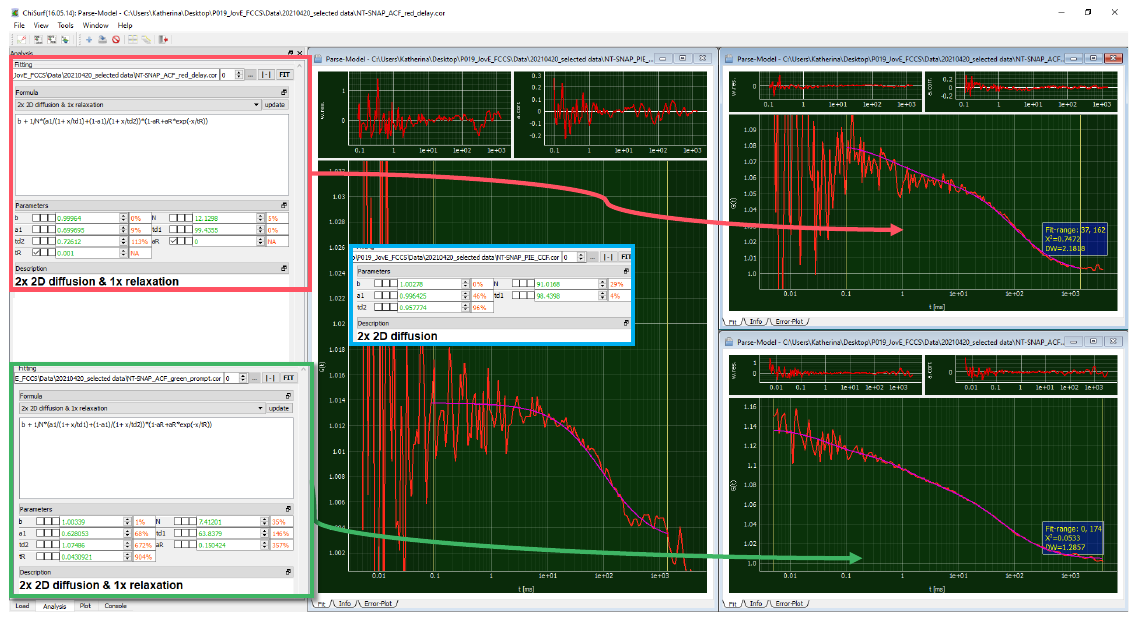

Figure 31 Global fit of both autocorrelation curves (green: eGFP, red: SNAP-substrate) and PIE-based crosscorrelation curve (blue, large window).

###### Calculation of co-diffusing molecules

Based on the (apparent) number of molecules in focus and our determined correction factors for the confocal overlap volume from the DNA measurements, the fraction of double-labeled molecules can be calculated (Figure 32).

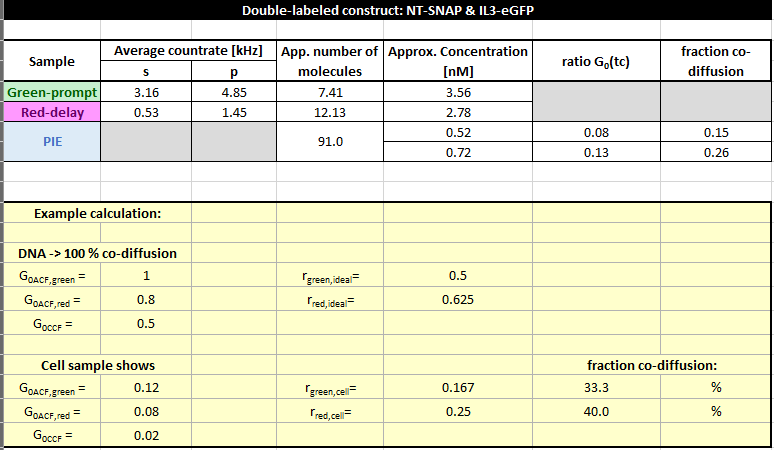

Figure 32 Calculation of the fraction of co-diffusing molecules.

Here, *N_eGFP_* = 7.4, *N_SNAP_* = 12.1 and *N_app,PIE_* = 91. Thus, the amplitudes are zero correlation time G(*t_c_*=0) have the following values: G_eGFP_(0) = 0.135, G_SNAP_(0) = 0.082, and G_PIE_(0) = 0.011.

Using the correction factors from the DNA measurement, here only between 15 -26 % molecules show both labels. However, (i) the data is quite noisy and (ii) the correlation amplitudes are very low. Both factors lead to large errors.

*Conclusion: Search for cells with low expression level, i.e. low fluorescence and take your time to collect a decent amount of photons to correlate!*

#### Analysis of simulated data

##### Simulation details

The simulations of the β_2_AR-eGFP-IL3-CT-SNAP measurements (short: CTSNAP) were performed using Burbulator ^5^ (part of the MFD software package, <https://www.mpc.hhu.de/software/3-software-package-for-mfd-fcs-and-mfis>).

The NTSNAP construct has both fluorophore on the inner side in a membrane and we assume (i) the fluorophore to be close enough to each other to undergo FRET and (ii) the membrane receptor β_2_AR to show dynamics such that the fluorophores exchange between two different levels of FRET.

Burbulator uses the Becker&Hickl spc-file format with 4096 TAC channel with a width of 4.07 ps and a laser period of 13.596 ns. The green-to-red detection efficiency ratio and the g-factor was set 1 and the fundamental anisotropy to 0.38. The fluorescence lifetime of both eGFP and SNAP was set to 3 ns with a molecular brightness of 10 kHz/molecule, fluorescence quantum yield of 0.8 and a rotational correlation time of 100 ns. Additionally the background in the green channel was set to 1 kHz and in the red channel to 0.5 kHz. The green crosstalk into the red channels was set to 0.1. All listed values for the fluorophores were adopted based on our measurements from the NTSNAP construct and the setup-describing values were set to reasonable values or 1.

The mean lifetime of the low FRET (LF) and high FRET (HF) states for dynamic exchange was set to 2.4 ns (*E* = 0.2) and 0.9 ns (*E* = 0.7). The equilibrium fractions of LF and HF were set to 0.5 each with a relaxation rate of 71 µs and – in case of triplet – with 16 % triplet blinking at 5.5 µs (Caution: Burbulator adds triplet blinking to both donor and acceptor molecules at the same time!).

The diffusion term was modeled as a bimodal distribution with 30 % of fast diffusing molecules at *t_D1_* = 1 ms and the rest of the molecules diffusing slowly with t*_D2_* = 100 ms.

In total, 10^7^ photons were simulated in a 3D Gaussian shaped volume with *w_0_* = 0.5 µm and *z_0_* = 1.5 µm, a box size of 20, and *N_FCS_* = 0.01.

*Of note*: As the slower modeled diffusion time is quite long, the number of photons and the box size might have to be increased further to allow good fitting at large correlation times *t_c_*. Here, for the sake of time / simplicity, the fitting was stopped at *t_c_* = 1 sec.

##### Global fit of auto- and cross-correlation curves

Before starting ChiSurf, add the fit model for the cross-correlation curve to your JSON-file as described above:

$$G_{CCF,2D}\left( t_{c} \right)=b+\frac{1}{N}\left[ \frac{a_{1}}{1+\frac{t_{c}}{t_{d1}}}+\frac{1-a_{1}}{1+\frac{t_{c}}{t_{d2}}} \right]\left[ 1-a_{R}\cdot\exp\left( -\frac{t_{c}}{t_{R}} \right) \right]$$

where *a_R_* and *t_R_* describe the amplitude and relaxation time of the anticorrelation.

A more general equation – in case of more than one relaxation term – would have the following form:

$$G_{CCF,2D}\left( t_{c} \right)=b+\frac{1}{N}\left[ \frac{a_{1}}{1+\frac{t_{c}}{t_{d1}}}+\frac{1-a_{1}}{1+\frac{t_{c}}{t_{d2}}} \right]\left[ 1-a_{f} \right]\left[ 1-\sum_{i} a_{Ri}\cdot\exp\left( -\frac{t_{c}}{t_{Ri}} \right) \right]$$

where *a_f_* describes the total amplitude of the anticorrelation (identical to *a_R_* in the single anticorrelation term model above) and *a_Ri_* and *t_Ri_* the respective relaxation times and amplitudes.

Load in total five different correlation curves into ChiSurf:

- Green-prompt (autocorrelation of green signal in prompt time window)
- Red-prompt (autocorrelation of the FRET-induced red signal in the prompt time window)
- Red-delay (autocorrelation of the red signal (direct excitation) in the delay time window)
- FRET-CCF (cross-correlation of green prompt and red-prompt signal)
- PIE-CCF (cross-correlation of green-prompt with red-delay)

The two new curves (red-delay and FRET-CCF), which we have not used to far, both stem from the FRET-induced red signal now present in our data (Figure 33).

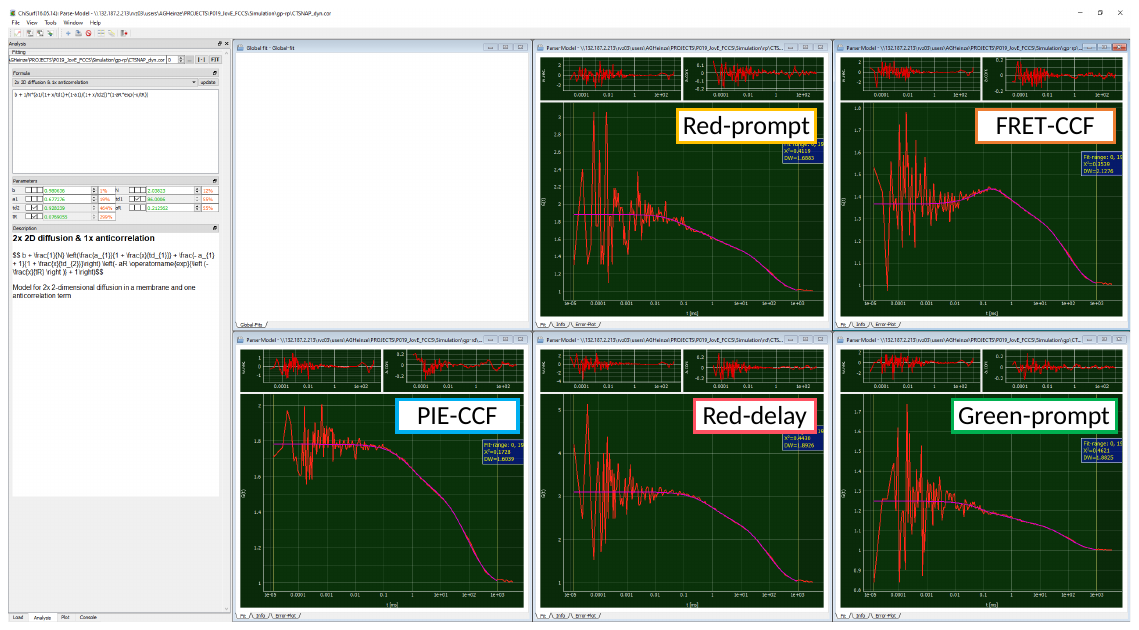

Figure 33 Global fit of all correlation curves of simulated data of a CT-SNAP construct. No triplet blinking was added.

Due to the FRET-induced anticorrelated behavior of green and red signal in the prompt time window. The FRET-CCF shows a “dip” at short correlation time, coinciding with a rise in both autocorrelation curves from the prompt time window.

All five loaded curves are fit jointly with linked *t_D1_*, *t_D2_* and *t_R_*. The fit results are summarized in the table below. Please note that here the diffusion times can be fit jointly as the simulation software does not support the modelling of differently sized confocal detection volumes. For experimental results, this joint fitting of *t_D_* might not be possible, however *t_R_* should be linked.

| Parameter | green-prompt | red-prompt | red-delay | FRET-CCF | PIE-CCF |
| --- | --- | --- | --- | --- | --- |
| *N(app)* | 3.90 | 1.09 | 0.455 | 2.04 | 1.23 |
| *t_D1_* [ms] | 86.0 | | | | |
| *a_1_* | 0.710 | 0.687 | 0.694 | 0.677 | 0.687 |
| *t_D2_* [ms] | 0.928 | | | | |
| *a_R_* | 0.203 | 0.144 | 0 | 0.213 | 0 |
| *t_R_* [µs] | 76.9 | | | | |

###### Influence of FRET efficiency

The extent of the anticorrelation can be used also as a marker for the extent of change in FRET efficiency *E*: If we change the LF state to *E* = 0 and the HF state to *E* = 0.95, the induced dip in the FRET-CCF is much more pronounced than for the first example (Figure 34).

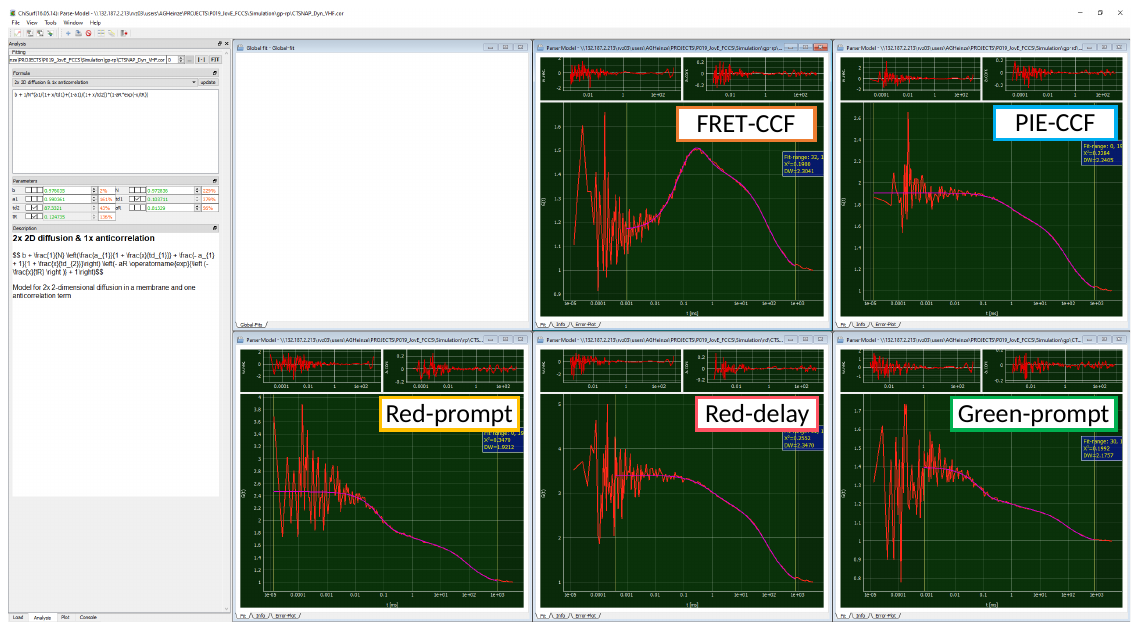

Figure 34 Global fit of all correlation curves of simulated data of a CT-SNAP construct with very high FRET contrast. No triplet blinking was added.

The amplitude of the anticorrelation has increased to *a_R_* = 0.66 compared to an *a_R_* = 0.21 from the previous case.

| Parameter | green-prompt | red-prompt | red-delay | FRET-CCF | PIE-CCF |
| --- | --- | --- | --- | --- | --- |
| *N(app)* | 2.45 | 0.664 | 0.397 | 0.973 | 1.05 |
| *t_D1_* [ms] | 91.8 | | | | |
| *a_1_* | 0.722 | 0.704 | 0.728 | 0.713 | 0.77 |
| *t_D2_* [ms] | 0.886 | | | | |
| *a_R_* | 0.39 | 0.392 | 0 | 0.662 | 0 |
| *t_R_* [µs] | 70.5 | | | | |

###### Influence of triplet blinking

Be aware that significant amount of triplet blinking in the µs time range of the donor and / or acceptor fluorophore might mask the anticorrelation induced due to FRET in the FRET-CCF.

In the example shown below (Figure 35), 16 % of additional triplet blinking at 5.5 µs was added to the example of LF(*E* = 0.2) <-> HF(*E* = 0.7).

Here, the FRET-CCF model has to be extended for a triplet-induced correlation term:

$$G_{CCF,2D}\left( t_{c} \right)=b+\frac{1}{N}\left[ \frac{a_{1}}{1+\frac{t_{c}}{t_{d1}}}+\frac{1-a_{1}}{1+\frac{t_{c}}{t_{d2}}} \right]\left[ 1-a_{R}\cdot\exp\left( -\frac{t_{c}}{t_{R}} \right) \right]\left[ 1-a_{T}+a_{T}\cdot\exp\left( -\frac{t_{c}}{t_{T}} \right) \right]$$

where *a_T_* and *t_T_* describe the amplitude and relaxation time of the triplet component.

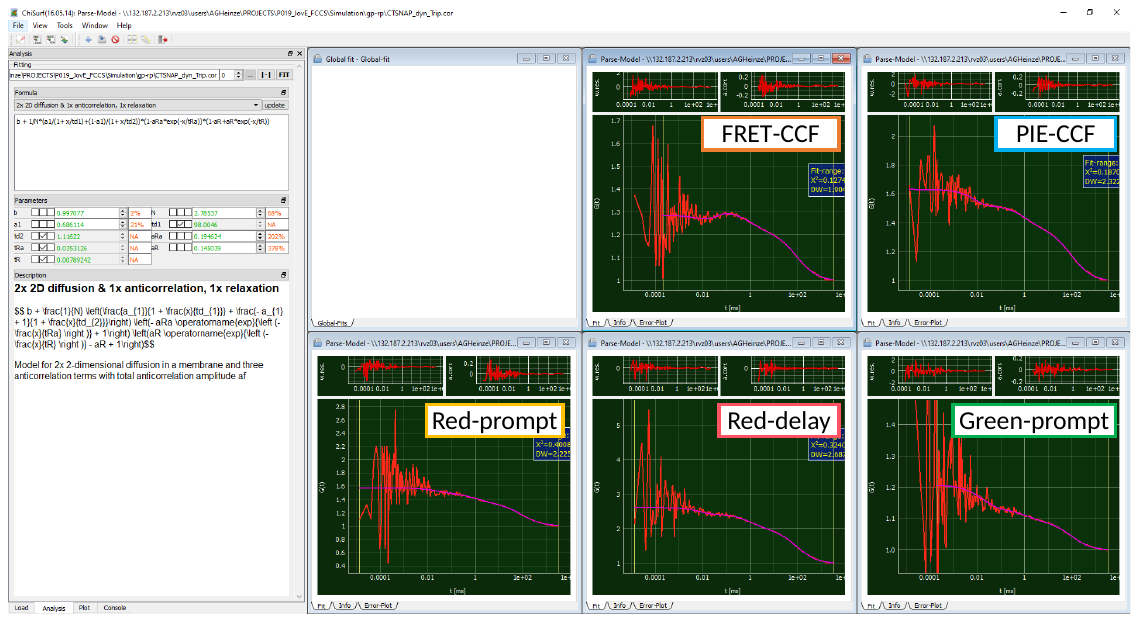

Figure 35 Global fit of all correlation curves of simulated data of a CT-SNAP construct with low FRET contrast and additional triplet blinking.

One can see quite nicely in the FRET-CCF how the “dip” in the curve due to the anticorrelation term is counteracted by the triplet component.

| Parameter | green-prompt | red-prompt | red-delay | FRET-CCF | PIE-CCF |
| --- | --- | --- | --- | --- | --- |
| *N(app)* | 4.79 | 1.73 | 0.611 | 1.73 | 1.57 |
| *t_D1_* [ms] | 98.0 | | | | |
| *a_1_* | 0.678 | 0.670 | 0.677 | 0.670 | 0.674 |
| *t_D2_* [ms] | 1.12 | | | | |
| *a_R_* | 0.210 | 0.115 | 0 | 0.115 | 0 |
| *t_R_* [µs] | 35.3 | | | | |
| *a_T_* | 0.146 | 0.015 | 0.113 | 0.015 | 0.171 |
| *t_T_* [µs] | 7.89 | | | | |

##### Species-filtered FCS to recover dynamics

A method to recover the triplet-masked anticorrelations in the FRET-CCF is to make use of the microtimes (i.e. the fluorescence decay histograms) encoded in the data. Here, instead of direct photon traces, an additional weighting function is introduced based on the fluorescence decay shape of the (i) IRF, (ii) the LF state, and (iii) the HF state. More details on how to generate these weighting functions can be found e.g. in the following literature ^6,7^.

In this species-specific or filtered FCS approach, four different correlation pattern are generated:

- Species-autocorrelation of the LF state (*sACF_LF-LF_*)
- Species-autocorrelation of the HF state (*sACF_HF-HF_*)
- Species-cross-correlation of the LF state to the HF state (*sCCF_LF-HF_*)
- Species-cross-correlation of the HF state to the LF state (*sCCF_HF-LF_*)

Below, exemplary the work flow and input for a filteredFCS analysis of the LF(*E* = 0.2) <-> HF(*E* = 0.7) example with additional triplet is shown (Figure 36).

| Normalized decays of species | Calculated filter for weighting | species-filtered FCS |
| --- | --- | --- |
| 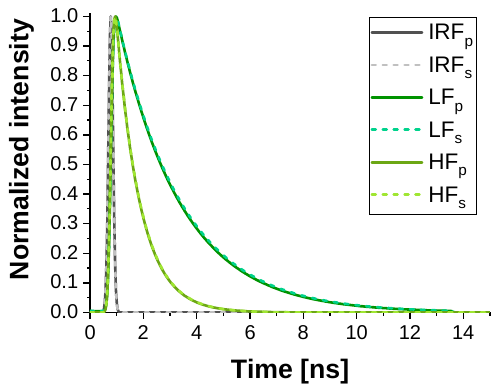 | 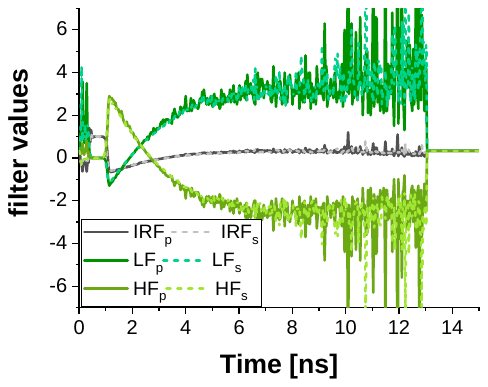 | 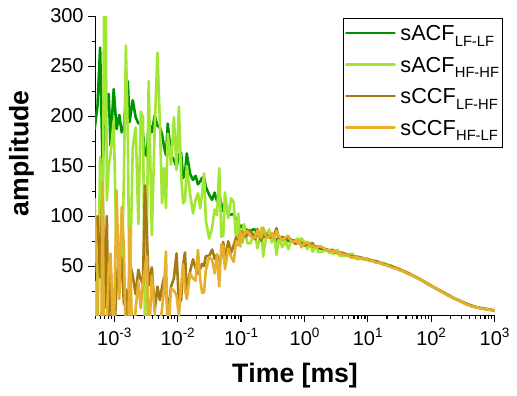 |
| Figure 36 Input and results for filteredFCS analysis of the data shown in Figure 35. Note: Suffix “p” and “s” are used to discriminate between the parallel (p) and perpendicular (s) channel here. | | |

For generation of the species-filtered FCS curves, the weights determined for the LF and HF species based on the normalized intensity decays are used during the correlation.

The resulting curves are fit to standard equations with bimodal membrane diffusion and relaxation and anticorrelation terms, respectively (Figure 37).

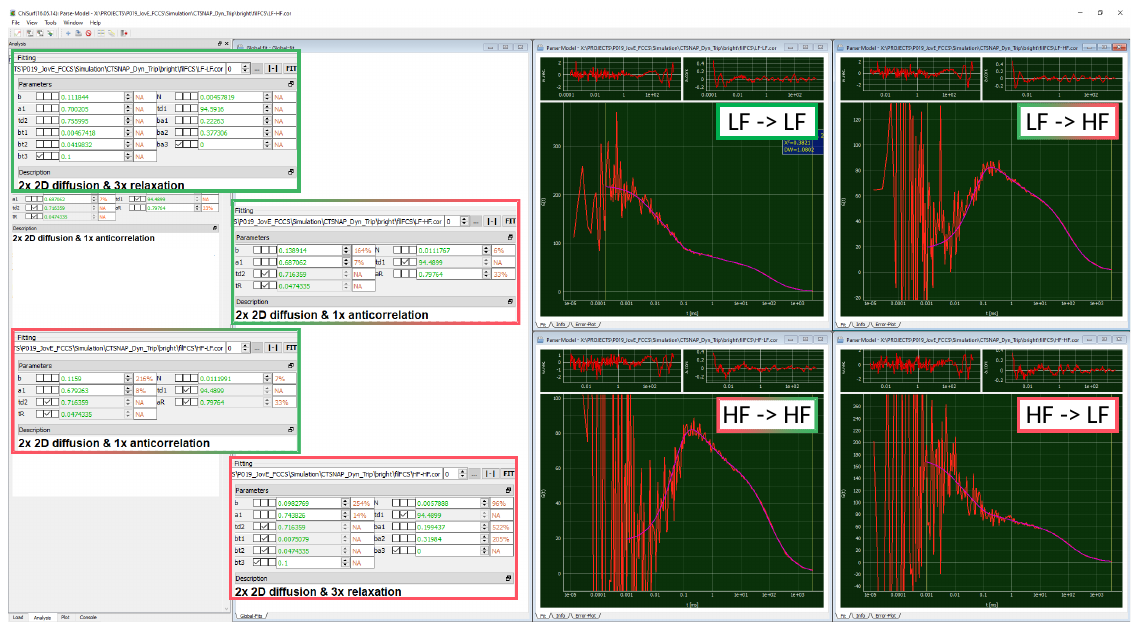

Figure 37 Global analysis of the generated filteredFCS curves

During the fit, both diffusion times *t_D1_* and *t_D2_* as well as the relaxation times are fit jointly and the FRET-induced relaxation of the *sACF_LF-LF_* and *sACF_HF-HF_* is linked to the anticorrelation term of the *sCCF_LF-HF_* and *sCCF_HF-LF_*.

*Be aware that the number of molecules in focus, N, is only an apparent number and does no longer relate to the concentration of molecules in the experiment / simulations!*

| Parameter | LF->LF | HF->HF | LF->HF | HF->LF |
| --- | --- | --- | --- | --- |
| *N(app)* | 0.0046 | 0.0058 | 0.011 | 0.011 |
| *t_D1_* [ms] | 94.6 | | | |
| *a_1_* | 0.700 | 0.687 | 0.687 | 0.679 |
| *t_D2_* [ms] | 0.756 | | | |
| *a_R_* | 0.378 | 0.320 | 0.798 | |
| *t_R_* [µs] | 42.0 | | | |
| *a_T_* | 0.223 | 0.199 | 0 | 0 |
| *t_T_* [µs] | 4.67 | | | |
