## Supplementary File 1-7 for "Dual-color Fluorescence Cross-Correlation Spectroscopy to study Protein-Protein Interaction and Protein Dynamics in Live Cells": SuppNote5_Fluorescence lifetime histograms.docx

### Fluorescence intensity of β_2_AR-IL3-eGFP, NT-SNAP-β_2_AR-IL3-eGFP and β_2_AR-IL3-eGFP-CT-SNAP

| 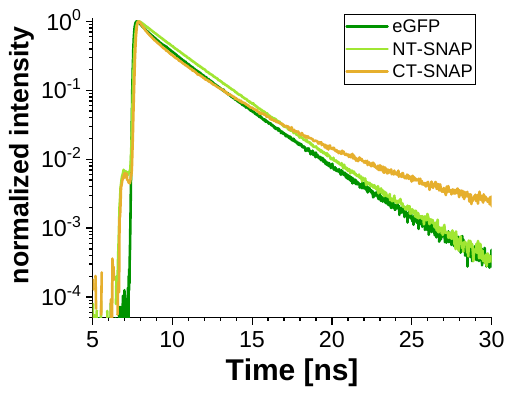 | 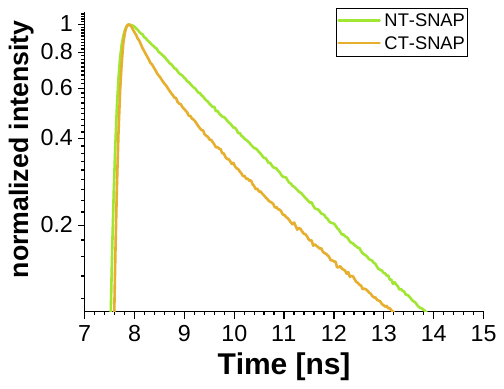 |
| --- | --- |
| Normalized photon arrival time histograms of β_2_AR-IL3-eGFP (green), NT-SNAP-β_2_AR-IL3-eGFP (light green) and β_2_AR-IL3-eGFP-CT-SNAP (orange). The Zoom-in on the right side shows clearly the “curved” course of the fluorescence intensity decay in the CT-SNAP induced by FRET while in contrast the NT-SNAP sample shows a nearly linear decay. | |
